# Sm-site containing mRNAs can accept Sm-rings and are downregulated in Spinal Muscular Atrophy

**DOI:** 10.1101/2024.10.09.617433

**Authors:** Anton J. Blatnik, Manu Sanjeev, Jacob Slivka, Benjamin Pastore, Caleb M. Embree, Wen Tang, Guramrit Singh, Arthur H. M. Burghes

## Abstract

Sm-ring assembly is important for the biogenesis, stability, and function of uridine-rich small nuclear RNAs (U snRNAs) involved in pre-mRNA splicing and histone pre-mRNA processing. Sm-ring assembly is cytoplasmic and dependent upon the Sm-site sequence and structural motif, ATP, and *Survival motor neuron* (SMN) protein complex. While RNAs other than U snRNAs were previously shown to associate with Sm proteins, whether this association follows Sm-ring assembly requirements is unknown. We systematically identified Sm-sites within the human and mouse transcriptomes and assessed whether these sites can accept Sm-rings. In addition to snRNAs, Sm-sites are highly prevalent in the 3’ untranslated regions of long messenger RNAs. RNA immunoprecipitation experiments confirm that Sm-site containing mRNAs associate with Sm proteins in the cytoplasm. In modified Sm-ring assembly assays, Sm-site containing RNAs, from either bulk polyadenylated RNAs or those transcribed *in vitro*, specifically associate with Sm proteins in an Sm-site and ATP-dependent manner. In cell and animal models of Spinal Muscular Atrophy (SMA), mRNAs containing Sm-sites are downregulated, suggesting reduced Sm-ring assembly on these mRNAs may contribute to SMA pathogenesis. Together, this study establishes that Sm-site containing mRNAs can accept Sm-rings and identifies a novel mechanism for Sm proteins in regulation of cytoplasmic mRNAs.

**GRAPHICAL ABSTRACT:** 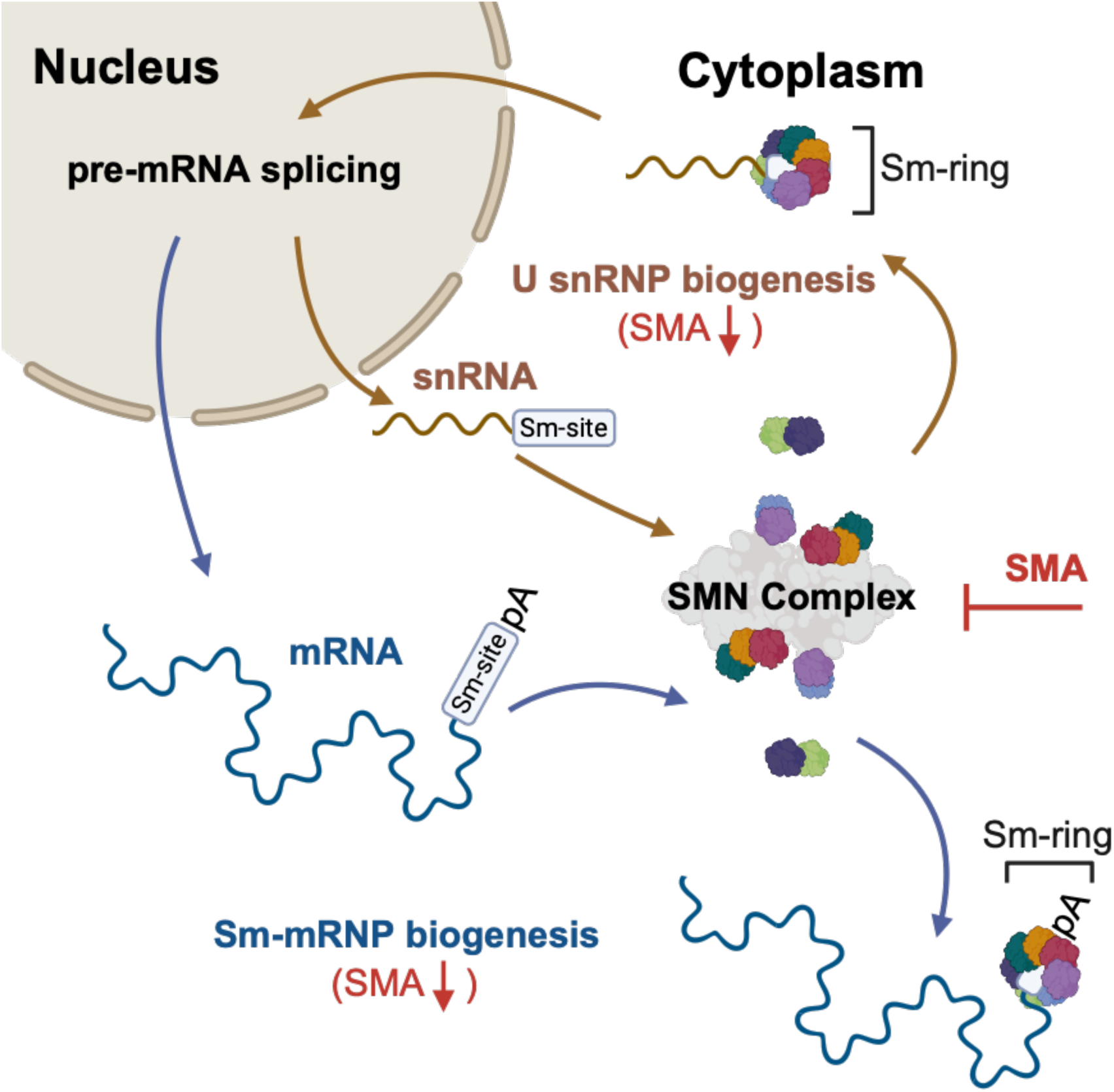

## INTRODUCTION

Sm-protein ring assembly is the process by which the seven Sm-proteins are assembled around a uridine-rich ribonucleic acid (RNA) sequence (1–6). This action is essential for the biogenesis and stability of spliceosomal, uridine-rich small nuclear RNAs (U snRNA) that function as the catalytic subunits of the spliceosome (7,8). The Sm-ring stabilizes U snRNAs and provides a handle for components of the spliceosome to interact with the assembled U snRNP (9,10). While RNAs beyond U snRNAs have been proposed to interact with Sm-ring-like assemblies (11), the prevalence of Sm-ring assembly sites in eukaryotic genomes and the possibility of Sm-ring assembly on RNAs beyond U snRNAs remains largely unexplored.

Sm-ring assembly is a highly choreographed process involving several trans-acting protein members. The Sm-ring is comprised of seven proteins—SmB/B’, SmD1, SmD2, SmD3, SmE, SmF, and SmG—that are preassembled by the PRMT5 complex into SmB/B’/D3 and SmD1/D2/E/F/G sub-complexes (5,12). These higher order Sm-subcomplexes are bound by pICln, kinetically trapping the Sm-proteins from assembling nonspecifically on RNA (5,12). The higher-order pICln:Sm blocks are then recruited to the SMN complex, composed of SMN, Gemins2-8, and Unrip, which then specifically assembles the full seven-member, heteromeric Sm-ring around a uridine-rich sequence motif within the U snRNA, known as an Sm-site (5,6,8,13–26).

The Sm-site is classically defined as 5’-AUUUUU(U)G-3’, found in the U2, U4, U4atac, U5, U11, and U12 snRNAs (2,4,22,27). The Sm-site in U1 snRNA is distinct, in which the fourth uridine is replaced by a guanosine to yield 5’-AUUUGUG-3’. Mutation of the third uridine to a cytosine does not alter Sm-ring formation (22). The U7 snRNA Sm-site is unique in that the sequence is longer —5’-AUUUGUCUAG-3’—, and results in the replacement of SmD1/D2 with Lsm10/11. This U7-specific hybrid Sm-ring is still assembled by the SMN complex, but results in a different protein association profile that facilitates the processing of histone pre-mRNAs (21,28,29). Apart from the uridine-rich sequence motif, the Sm-site includes a 3’ stem loop necessary for Sm-ring assembly (22,27). The Sm-ring forms a toroidal structure, in which individual bases directly interact with individual members of the Sm-ring, making a core that is stable under high salt, heparin, and urea conditions (1–7,30,31). Following Sm-ring assembly, U snRNA m^7^G-caps are hypermethylated (m^2,2,7^G), their 3’ ends are trimmed (25,32–36) and they are imported into the nucleus through a mechanism involving snurportin-1 and importin-beta (25,33,34,37–43). Within the nucleus, nucleotides of U snRNPs are pseudouridylated or 2’-O-methylated (44,45) and they associate with their intended RNA targets and protein effectors; pre-mRNA and the spliceosomal machinery in the case of spliceosomal U snRNPs, and histone pre-mRNA and processing factors for U7 snRNPs (46,47).

Sm-ring assembly is an ATP, SMN, and Sm-site dependent process in cell extracts (20,48). Though Sm-proteins will spontaneously form Sm-rings on RNA when in high concentration, the PRMT5/SMN complex system inhibits this promiscuous assembly in favor of the Sm-site (5). Sm-ring assembly requires the addition of ATP in cell and oocyte extracts (20,48), however reconstitution of ring assembly components *in vitro* indicate Sm-ring assembly is a Brownian activity involving a minimal SMN complex—SMN, Gemin2,6-8, Unrip, (12). It was recently reported that the DEAD-box helicase, Gemin3, confers the requirement for ATP, unfolding U snRNA secondary structure that can shield the Sm-site (49). Modification of the first and third uridine positions in the Sm-site, or removal of stem loop structures 3’ of the Sm-site sequence ablates Sm-ring assembly (22,27). Furthermore, SMN protein deficiency—causal of the human disease Spinal Muscular Atrophy (SMA)—results in a reduced capacity to assemble spliceosomal and U7 snRNPs, indicating Sm-ring assembly is SMN dependent (29,48,50).

Reductions in SMN protein abundance have been shown to perturb several RNA processing events including transcription (51,52), pre-mRNA splicing (53), U snRNP assembly (6,13,20,29,48,54–58), histone pre-mRNA processing (21,29), snoRNP assembly (43,59–61), telomerase activity (43,61), translation (62–65), signal recognition particle biogenesis (66), and mRNA trafficking (67–76). However, a direct role for SMN in these events has not been forthcoming. A previous report indicated many types of RNAs associate with Sm proteins but did not establish that these RNAs could accept an Sm-ring (11). Furthermore, Sm-ring assembly was found to be prerequisite to Lsm-ring assembly on the fission yeast telomerase RNA subunit (TER1) (77). We hypothesized that Sm-sites are present within many other RNAs beyond U snRNAs, that Sm-rings are assembled upon them, and this process may provide a direct link between accepted SMN function and the aforementioned RNA processing events. Our informatic search revealed that many human and mouse RNAs contain Sm-sites, but most notably, Sm-sites are enriched in mRNA 3’ untranslated regions (UTRs). To investigate whether prediction of an Sm-site informed enrichment upon Sm-protein immunoprecipitation, we modified U snRNP Sm-ring assembly assays to detect ring assembly on polyA-enriched RNA using anti-Sm RNA immunoprecipitation and next-generation sequencing (RIP-Seq). We report that many Sm-site containing mRNAs associate with Sm-rings and that Sm-protein ring assembly on these RNAs can occur in an ATP and Sm-site specific manner. Lastly, Sm-site containing mRNAs are downregulated in models of SMA, potentially identifying a link between SMN and metabolism of novel Sm-site-containing mRNAs.

## MATERIAL AND METHODS

### **I**nformatics pipeline to identify Sm-site containing RNAs

A miniconda3 environement was created to run SQlite3, SeqIO, pandas, numpy, and Bioconda packages Bio.Seq, and SeqUtils. A database was generated using SQlite3 from the NCBI Refseq and Gencode annotations of the human GRCh38 and mouse GRCm39 genomes that could easily recall gene ID, transcript ID, gene name, and coding sequence start and stop from the GFF files. Next, a python script was written to create a cursor that would search the supplied transcriptome fasta files for the presence of a U1 (5’-AUUUGUG-3’), U2 (5’-AUUUUUG-3’), U5 (5’-AUUUUUUG-3’), U7 (5’-AUUUGUCUAG-3’), or Noncanonical (5’-AUNUKUN-3’, where K is a G or U) sequence and capture the preceding 5’ 200 nucleotides proceeding 3’ 16-50 nucleotides into two separate output files. The aforementioned databases are used to populate information like the transcript ID, gene ID, and the region the Sm-site sequence was located within the transcript. This generates two output files for each Sm-site type, one for the preceding and one for the proceeding sequence. Next, these Sm-site output files were provided as input for RNAfold (ViennaRNA Package 2.0, version 2.6.4) (78) which predicts secondary structure for both the preceding and proceeding Sm-site sequences, resulting in a file providing the transcript ID, the queried sequence, and the structure prediction. A third python script was then used to determine if a secondary structure is predicted within 20 nucleotides 5’ or 10 nucleotides 3’ of the Sm-site using recognition. This script then discards all identified Sm-sites that are not flanked by secondary structures on both sides. This yields a single output file for each type of Sm-site identified with the associated transcript ID, gene name, the region the Sm-site is found in, and the proceeding 3’ sequence used in structure prediction. These output files are supplied in Supplementary File 1 and were then used to generate all following master tables. The python scripts used to generate this data are available in Github <https://github.com/ajblatnik/sm_ring_assembly_mrna.git>.

For the figures displayed within the study, frequency of Sm-sites is given for different regions and different transcripts on a gene-level basis. In this, each gene ID corresponds to a single transcript ID, and only Sm-sites identified for that transcript ID were used from the output files. To facilitate this, R (version 4.4.1) was used to cross-reference the shared and annotation-specific Sm-sites identified in the Refseq and Gencode Sm-site output files in Supplementary File 1, by first retaining those transcript IDs that have shared proceeding 3’ sequence and then appending sites specific to Refseq or Gencode. The frequency of Sm-sites identified for each transcript and transcript region was calculated and a master table was generated by inner-joining with a list of gene IDs pared down to a single transcript ID, prioritizing by MANE and Ensembl Transcript is Canonical notation as provided in BiomaRt (79,80). This was repeated for each type (U1, U2, U5, U7, and Noncanonical) to generate a single table that contained all information. Given that U1, U2, and U5 Sm-site types will satisfy the Noncanonical arguments, the frequency of true Noncanonical Sm-sites was determined by subtracting the frequency of noncanonical Sm-sites from the raw output by the number of U1, U2, and U5 Sm-site types. Canonical Sm-site frequencies were determined by adding the frequencies for U1, U2, and U5 Sm-site types. Transcript, 5’UTR, CDS, and 3’UTR lengths were then populated into these master tables which are available as Supplementary File 2. The R scripts and session info used to generate these tables are available in Github <https://github.com/ajblatnik/sm_ring_assembly_mrna.git> and these tables were then used to generate all the data and graphs within the study.

### Cell Culture

Neural stem cell NSC-34 cells were cultured in 12% FBS DMEM supplemented with L-glutamine and PenStrep (12% FBS Atlanta S11550, 10% PenStrep/L-Glutamine Gibco 10378-016, DMEM Gibco 11960077) under 37 °C, 5% CO_2_ in normoxic conditions. Cells were routinely passaged every 2-3 days by washing twice with Phosphate Buffered Saline lacking divalent cations, and then treating with 0.05% Trypsin-EDTA (Gibco 25300054). S3 fibroblasts were induced to a neural progenitor cell fate as described by Meyer *et al* (81). To culture neural progenitor cell S3-iNPCs, dishes were coated with fibronectin (Millipore FC010-10MG) and cells were cultured in DMEM/F12+ Glutamax (Gibco 10565-042) supplemented with 10% N2 (Gibco 17502-048), 10% B27 (Gibco 17504-044), 10% Anti-Anti (Gibco 15270-62), and 20 ng/mL FGF2 ( Peprotech 100-18B), under 37 °C, 5% CO_2_ in normoxic conditions. Cells were routinely passaged every 2-3 days by washing twice with PBS lacking divalent cations and treating with StemPro Accutase (ThermoFisher A1110501).

### Oligo-dT isolation of polyA-RNA

Four 15 cm dishes of NSC-34 or S3-iNPC cell were washed three times with PBS lacking divalent cations and treated with 8 mL TRIzol Reagent (Invitrogen 15596026) per 15 cm dish. TRIzol samples were pooled and a large scale total RNA isolation and precipitation was performed following manufacturer’s instructions. After purification, isolated RNA was treated with 200 units of Turbo DNase (Invitrogen AM2239) and incubated at 37 °C for 30 minutes with gentle rotation at 350 rpm in Eppendorf Thermomixer R. DNase was removed by phenol:chloroform extraction. RNA was precipitated with 1.5 M final concentration of LiCl and 100% ethanol, centrifuged at > 18,000 x g for 15 minutes. RNA was resuspended in 1 mL nuclease-free water and quantified using Qubit RNA Broad Range Assay (Invitrogen Q10211). PolyA-RNA was isolated using Dynabeads mRNA Purification Kit (Invitrogen 61006). Oligo-dT beads were reused post-polyA-RNA elution to isolate the magnitude of polyA-RNA required for modified Sm-ring assembly reaction and sequencing conditions. After isolation, polyA-RNA was quantified using Qubit RNA Broad Range Assay (Invitrogen Q10211) and RNA integrity (RIN) was quantified by TapeStation analysis performed by the Ohio State Comprehensive Cancer Center Genomics Core to determine removal of polyA-minus RNAs (Supplementary Figure 5C).

### Sm-ring assembly assays

Preparation of cytoplasmic cell extracts and standard Sm-ring assembly assays were performed as previously described in Wan *et al* 2005, and Blatnik *et al* 2020 (48,82). Cytoplasmic cell extracts were generated by scraping and collecting NSC-34 or S3-iNPC cells in cold phosphate buffered saline containing only monovalent cations. Cells were pelleted by centrifugation 400 x g for 4 min. Cells were resuspended in digitoxin lysis buffer (20 mM Tris pH 8, 50 mM KCl, 0.5 MgCl_2_, 1x eComplete, 200 units/mL RNase-OUT, 50 ug/ mL digitonin, in DEPC-treated water) and incubated on ice for a few minutes. Lysates were passaged 6 times through a 27 gauge needle with syringe then centrifuged at 1,500 x g for 1 minute at 4 °C. Supernatant was measured and transferred to a new microfuge tube and supplemented with NP-40 to achieve 0.01% final concentration. Lysates were then centrifuged at 4,000 x g for 15 minutes at 4 °C. Supernatants were then aliquoted into microfuge tubes and frozen in liquid nitrogen. Protein concentrations were determined by BCA assay (Pierce 23250). Protein-A DynaBeads (Invitrogen 10001D) were aliquoted 8.6 µL/rxn and washed three times in PBS + 0.1% NP-40. 0-3.2 µg/rxn Y12 antibody (Novus Biologicals NB600-546, EMD Millipore MABF2793, Invitrogen MA5-13449) or SmB/B’/N (12F5) (Santa Cruz sc-130670) was conjugated to Protein-A DynaBeads by incubating in PBS + 0.1% NP-40 for 1 hour at 4 °C with rotation. 1.6 µg/rxn Novus Biologicals NB600-546 Y12 antibody was used for all subsequent experiments. Beads were washed 3 more times with PBS + 0.1% NP-40, then three times with RSB-500 + 0.1% NP-40 (RSB-500: 10 mM Tris pH 7.4, 500 mM NaCl, 2.5 mM MgCl_2_). Beads were finally resuspended in 120 µL/rxn RSB-500 + 0.1% NP-40 supplemented with 2 mg/mL heparin. Sm-ring assembly was performed in 20 µL total volume and incubated for 1 hour at 30 °C with shake at 750 rpm (1 unit/µL RNAse-OUT, 0.25 ug/uL yeast tRNA, 0.01% NP-40, 2.5 mM ATP, 20 mM Tris pH 8, 50 mM KCl, 0.5 MgCl_2_, 25 µg cytoplamsic extract, and 10 nM biotinylated U4, U4ΔSm snRNA, mRNA, or mRNA 3’UTR). Assembly reactions were immunoprecipitated with 120 µL/rxn Y12-conjugated dynabeads, incubated for 1 hour at 30 °C with shaking at 750 rpm. Beads were washed 8 times with RSB-500 + 0.1% NP-40, utilizing magnets to sequester beads. Assembled snRNPs were detected by incubating the reactions with 120 uL RSB-500 + 0.1% NP-40 supplemented with 1:10000 dilution of NeutrAvidin-HRP, (Invitrogen A2664) for 1 hour at 30 °C with shake at 750 rpm. Beads were further washed 8 times with RSB-500 + 0.1% NP-40, utilizing magnets to sequester beads. Reactions were finally resuspended in 150 uL of 1:1 mixture of SuperSignal Fempto ELISA substrate (Pierce 37075), transferred to Nunclon white-bottom plate (Nunclon Delta Surface 136101). Luminescence was measured on Tecan Infinite F200 using I-Control software or Promega GloMAX plate reader, no attenuation, 1000 ms integration, 0 ms settle. Raw luminescence values were plotted for +ATP, -ATP, ΔSm + ATP, and ΔSm - ATP conditions in Supplementary Figure 15. Conditions were normalized to the ΔSm - ATP condition for Figure 4D-J. Graphs were plotted using GraphPad Prism Version 10.2.3 (347). For testing Sm-ring assembly directly on mRNA and mRNA 3’UTRs, prior to incubation with Y12 antibody, reactions were supplemented with 2 M urea and 5 mg/mL heparin and incubated for 15 minutes at 30 °C.

For modified Sm-ring assembly reactions, reactions were scaled up to 250 µg human S3-iNPC and 438 µg mouse NSC-34 total protein cytoplasmic extract, supplemented with 4 µg polyA-RNA (RIN 5-6, Supplementary Figure 5C) to capture sufficient RNA for sequencing library generation. 10 nM U4 snRNA is routinely used to perform assembly reactions in 25 µg total protein cytoplasmic lysate, however human U4 snRNA is only 141 nt in length. Therefore, approximately 18 nM polyA-RNA was used as input for the assembly reactions, assuming an average mRNA length of ∼3400 nt in humans and ∼3200 nt in mouse. Three replicates of the four conditions presented in Supplementary Figure 4B for both species combinations were used to generate sequencing libraries, using 50 ng of captured RNA as input for the Takara SMARTer Stranded Total RNA-Seq Kit v3-Pico Input Mammalian (Takara 634451), and sequenced using the NOVA-Seq platform, 150 base-pair paired-end reads. In words, the four conditions are: 1) the polyA-RNA input used for the assembly reactions, 2) anti-Sm-RIP of the cytoplasmic cell extract, 3) anti-Sm-RIP of cell extracts incubated with polyA-RNA, and 4) anti-Sm-RIP of cell extracts incubated with polyA-RNA and ATP but are also described in Supplementary Figure 4B.

### Informatic analysis of sequencing data

*For sequencing experiments discussed in Supplementary* Figure 4B: Table 1 lists the sequencing data generated in this study. Adapter sequences were trimmed using Cutadapt. Reads were aligned to both the human GRCh38 and mouse GRCm39 genome assemblies (Ensembl release 100) using STAR (83). Reads were counted using FeatureCounts from the subread package, using the fractional (—*fraction*) argument for reads mapped to multiple loci. Bash scripts used to process reads and MultiQC (84) html are provided to assess quality of reads, alignment, and features in Github <https://github.com/ajblatnik/sm_ring_assembly_mrna.git>, and percent uniquely mapped reads for alignment are provided in Supplementary Figure 5D. Wald-test comparisons of the FeatureCounts (85) outputs were performed using Deseq2 (86) in R (version 4.4.1). For all comparisons, the adjusted *p-*values from Deseq2 were used. Comparisons performed are discussed in Supplementary Figure 4B and Figure 4A. Principal component analysis was performed on the top 1000 genes to ensure proper clustering of samples (Supplementary Figure 5E). Custom R scripts were then written to analyze the data presented within this manuscript and the R session info are available in Github <https://github.com/ajblatnik/sm_ring_assembly_mrna.git>.

**Table 1:**
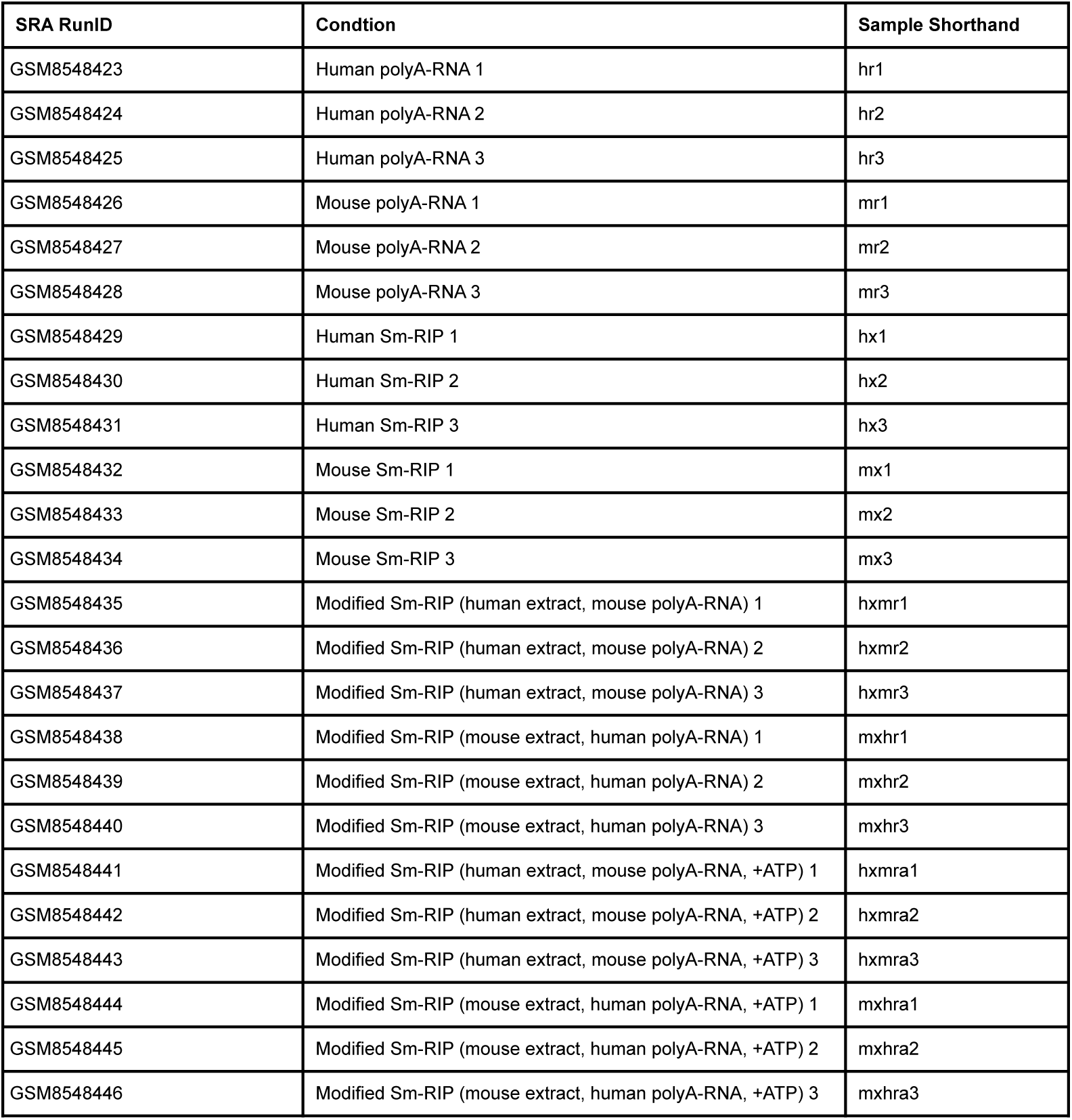
Sm-RIP sequencing data generated for this study (GSE278538)

#### For analyses of sequences discussed in Figures 5-6

Table 2 lists the publicly available sequencing data analyzed in the study. Data for each sample was retrieved from the NCBI Short Read Archive using their corresponding RunIDs with the fastqdump function of the SRA toolkit. The downloaded libraries were free from significant adapter content, eliminating the need for trimming. Sequencing reads were aligned to the genome (GRCh38/GRCm39, Ensembl release100) utilizing the STAR aligner (83). The aligned reads were then quantified using FeatureCounts (85) from the subread package. Differential expression was calculated for SMN deficient conditions by running DESeq2 (86) on the Featurecounts output. For each study, principal component analysis was performed with 2000 genes showing most variance to confirm that samples clustered by disease condition. Genes without a calculated *padj* value after DESeq2 were discarded. The cumulative distribution of log2 foldchanges was plotted after grouping genes based on the presence/absence of a canonical Sm-site.

**Table 2:**
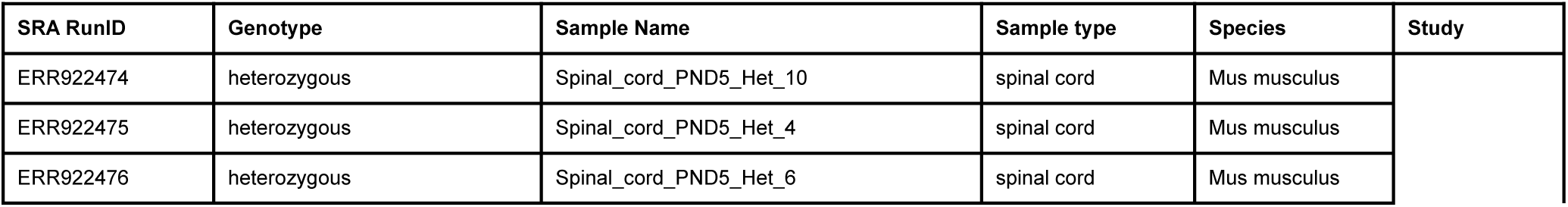
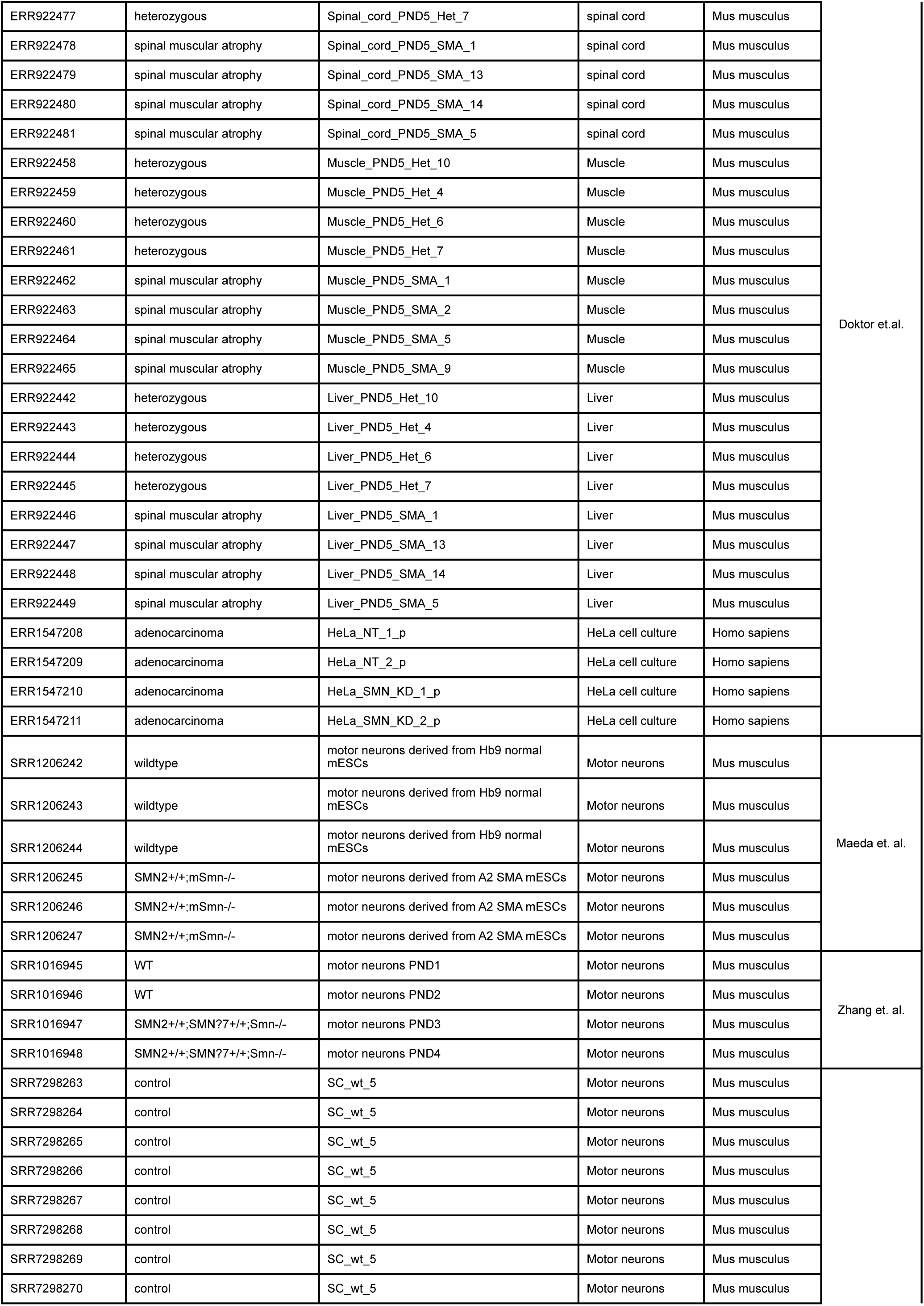
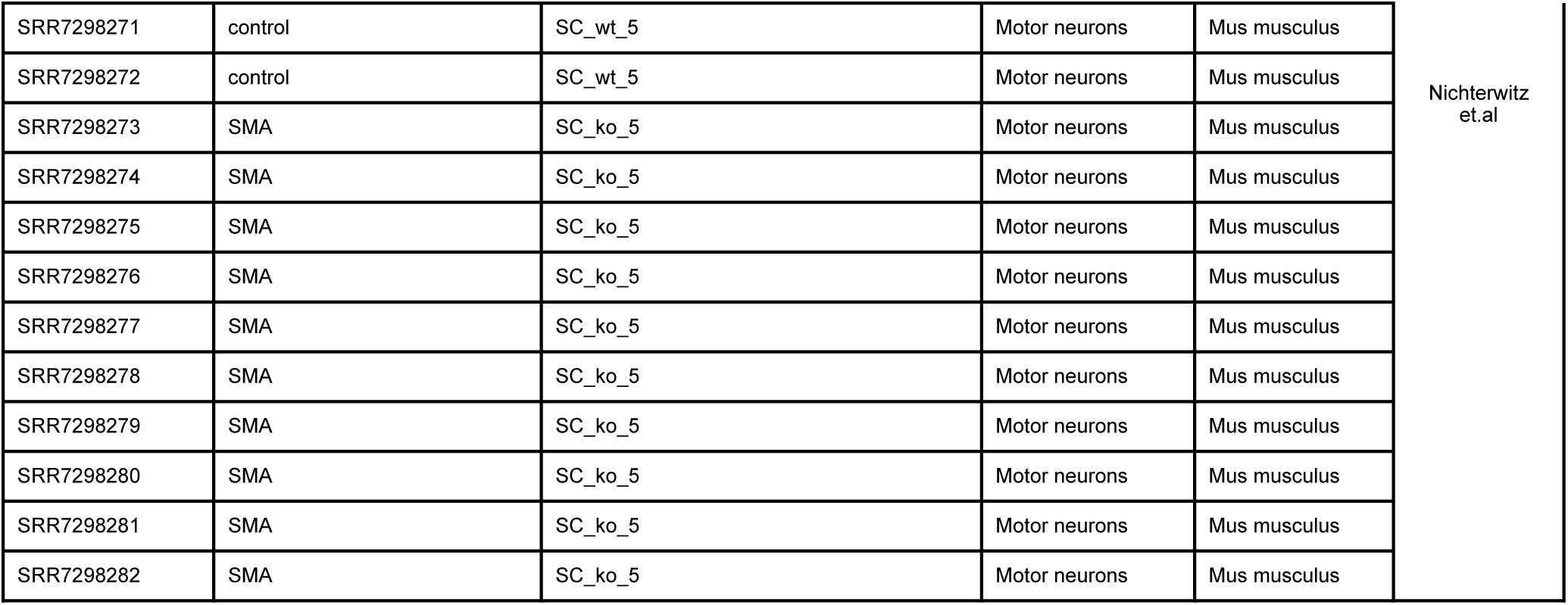
Publicly available sequencing data analyzed in this study.

### Cloning and *in vitro* transcription of mRNA and mRNA-3’UTRs

Candidate mRNA and mRNA-3’UTRs were selected as described within the results section. RNA sequences were imported from Ensembl, and all Sm-sites (canonical and noncanonical) were identified using Supplementary File 3 using a word processor. A duplicate sequence was generated in which each Sm-site (canonical and noncanonical) was mutated to 5’-ACCCCCG-3’ or 5’-ACUCUCG-3’ to provide the ΔSm sequence (Supplementary File 3). Sequences were then loaded into ApE-A plasmid Editor v2.0.61 to identify restriction enzymes that could be added to linearize plasmids for *in vitro* transcription. T7 promoter sequence was added to the 5’-end of the sequence, and PmlI or AflIII restriction sites were added to the 3’-end of the sequence. Invitrogen GeneART Synthesis services were used to generate plasmids (maps available in Supplementary File 4 and reagents can be made available upon request). Upon delivery of plasmids, sequences were confirmed by Sanger Sequencing. Plasmids were linearized using restriction digestion, followed by phenol:chloroform extraction and ethanol precipitation isolation methods. RNAs were *in vitro* transcribed using the HiScribe T7 High Yield Kit (New England Biolabs SE20405), substituting 25% of the UTP concentration with Bio-16-UTP (Invitrogen AM8452), following manufacturer’s instructions. Transcribed, labelled RNAs were purified using Urea-PAGE and correct band sizes were isolated using ultraviolet light shadow casting against a thin layer chromatography plate. RNAs were eluted from polyacrylamide slices in nuclease-free water overnight at 4 °C on a nutating mixer. RNAs were quantified using Qubit RNA Broad Range Assay (Invitrogen Q10211).

#### Reagents

Protein concentrations were determined by BCA assay (Pierce 23250). Sm-ring assembly reactions used Mouse mab Smith Antigen Ab (Y12) (Novus Biologicals NB600-546) or SmB/B’/N (12F5) (Santa Cruz sc-130670) antibodies bound to Protein-A Dynabeads (Invitrogen 10001D) for Sm-protein immunoprecipitation, and NeutrAvidin-HRP (Invitrogen A2664) for biotin detection. Other Y12 antibodies tested include SNRPB Monoclonal Antibody (Y12) (Invitrogen MA5-13449) and Anti-Smith Antigen Antibody, clone Y12 (Millipore Sigma MABF2793). Horse Radish Peroxidase (HRP) signal was detected using SuperSignal Fempto ELISA substrate (Pierce 37075). anti-Sm-RIP-Seq experiments were performed using anti-Sm(Y12) antibody (Novus Biologicals NB600-546). PolyA-RNA was isolated using Dynabeads mRNA Purification Kit (Invitrogen 61006). Sequencing libraries were prepared using Takara SMARTer Stranded Total RNA-Seq Kit v3-Pico Input Mammalian (Takara 634451). Plasmids used for *in vitro* transcription (Supplementary File 4) were restricted digested with PmlI (New England Biolabs R0532L), AfeI (New England Biolabs R0652L), or AflIII (New England Biolabs R0541L). Long RNAs were *in vitro* transcribed using the HiScribe T7 High Yield Kit (New England Biolabs SE20405). U4 snRNA was *in vitro* transcribed using MEGAshortscript T7 Transcription Kit (Invitrogen AM1354). Biotin labelling was performed by supplementing *in vitro* transcription reactions with with Bio-16-UTP (Invitrogen AM8452). RNA was quantified using Qubit RNA Broad Range Assay (Invitrogen Q10211).

#### Biological Resources

Cell lines used include neural stem cell-like NSC-34 and S3-induced neural progenitor cells (iNPC) (81). Plasmids for *in vitro* transcription and labelling of U4 and U4ΔSm were supplied by the lab of Dr. Dan Battle, formerly at The Ohio State University. Sequences for plasmids used for *in vitro* transcription and labeling of mouse and human candidate mRNA or mRNA 3’UTRs are provided in Supplementary Files 3 and 4. All biological reagents can be made available upon request.

### Statistical Analyses

Graphing was performed using ggplot2 in R (version 4.4.1) or GraphPad Prism (version 10.3.0 (461)). Statistics were performed in R (version 4.4.1). Wald-test comparisons were performed by DESeq2 (version 1.44.0). Pearson correlations were performed for continuous distribution comparisons. Kendell correlations were performed for rank-order comparisons. Wilcoxon tests were performed for all cumulative distribution functions, using the assumptions that the curves were either ‘greater’ or ‘less’ than the curve being compared to. Specific tests, their n, and data being compared are given in each specific plot and figure legend. R session info is provided in Github <https://github.com/ajblatnik/sm_ring_assembly_mrna.git>.

#### Novel Programs, Software, Algorithms

Informatics pipelines for identifying Sm-sites, for processing sequencing data, and generating figures are provided in Github <https://github.com/ajblatnik/ sm_ring_assembly_mrna.git>.

#### Data Base Referencing

Human and mouse NCBI RefSeq (87) reference genomes and transcript sequences were downloaded from <https://www.ncbi.nlm.nih.gov/datasets/>. Human and mouse Gencode (88) reference genomes and transcript sequences were downloaded from <https://www.gencodegenes.org/>. Genomic features, inlcuding transcript lengths, biotype, etc… were pulled using Ensembl BiomaRt (89) package installed in R (version 4.4.1). The RNA Binding Protein Data Base (RBPDB) <http://rbpdb.ccbr.utoronto.ca/> (90) was used to search for known proteins that bind sequences similar to the Sm-site. BioRender was used to generate some images in figures, all figures were assembled in Adobe Illustrator 2024 using an Adobe Creative Cloud license. Tables used to generate figures are available in Supplementary Files 1, 2 and 5. All scripts used to process data are available in Github <https://github.com/ajblatnik/sm_ring_assembly_mrna.git>.

## RESULTS

### Sm-sites are commonly found in the 3’UTRs of long mRNAs

To identify all RNAs in the human and mouse transcriptomes that have features of an Sm-site, we developed a custom algorithm to identify Sm-sites in the corresponding RefSeq and Gencode databases. First, the algorithm identifies transcripts that contain canonical or noncanonical Sm-site sequences. As the canonical Sm-protein ring is classically assembled on U1, U2, and U5-type Sm-site sequences, we defined canonical Sm-site sequences as those pertaining to U1, U2, and U5 snRNAs. Sm-sites from U4, U12 and U4atac snRNAs are classified within the U2-type, and the U11 snRNA site is classified in the U5-type (see Supplementary Figure 1 for a breakdown of the individual types of Sm-sites within snRNAs). The noncanonical Sm-site sequence was defined as—5’-AUNU(U/G)UN-3’—allowing for base substitutions that do not disrupt Sm-ring assembly (22). Transcript sequences containing the Sm-site sequence were further filtered to require an RNA secondary structure to start within the 20 nucleotides 5’ of the Sm-site sequence and 10 nucleotides 3’ of the sequence. A schematic describing the sequences and information supplied to the algorithm is provided in Figure 1A. The identified Sm-sites were cross-referenced between both NCBI RefSeq and Gencode analyses. The output for each type of Sm-site detected within the human and mouse transcriptomes is provided in Supplementary File 1. These data are summarized, providing counts of each Sm-site type on a per-gene basis in Supplementary File 2 (See methods for a more detailed description of these tables and their construction). Importantly, the identification of several known Sm-site containing snRNAs and the exclusion of the U6 snRNAs from the predictions demonstrates the validity of our approach for unbiased identification of Sm-site containing RNAs (Supplementary Figure 1). The inability of our algorithm to identify an Sm-site in human or mouse U4atac suggest its limitations (Supplementary Figure 1, Supplementary File 1-2).

**Figure 1.**
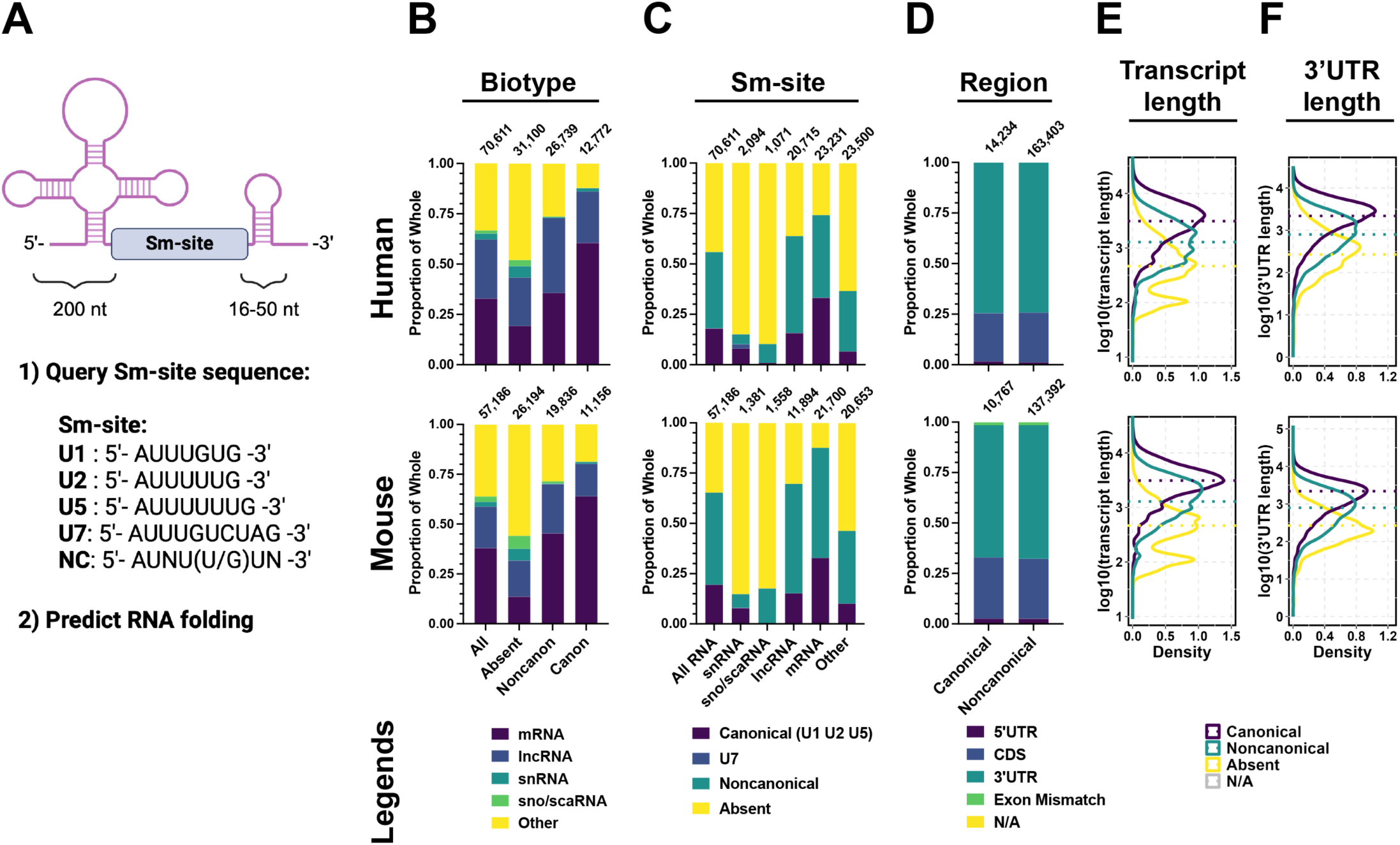
Sm-sites are commonly found in 3’UTRs of long mRNAs. **(A)** Generalized schematic of an snRNA, and the features queried by the search algorithm. First, the NCBI Refseq and Gencode transcriptomes were queried for presence of an Sm-site sequence—defined as U1, U2, U5, U7 or Noncanonical. Second, the 200 nt to the 5’ side of Sm site and 16-50 nt to its 3’ side were queried for thermodynamic favorability to fold using RNAfold. **(top)** for human genes, **(bottom)** for mouse genes. For all, bar graphs, top numbers are the number of genes represented within the bar. Data presented in **B-F** correspond to a single, unique transcript ID of a single, unique gene. Genes producing multiple transcripts are pared to only correspond to a single MANE or Ensembl Transcript is Canonical containing transcript. **(B)** Proportional bar graph of RNA biotypes. **All** is a breakdown of the different RNA biotypes within the annotated genome. **Absent** are RNAs not predicted to contain an Sm-site. **Noncanonical** are those RNAs only predicted to contain noncanonical Sm-site sequences. **Canonical** are those RNAs predicted to have at least one U1, U2, or U5 Sm-site sequence, but may have additional canonical or noncanonical sequences. **(C)** Proportional bar graph giving a breakdown of types of Sm-sites predicted in each of the following biotypes: All, snRNA, sno/scaRNA, lncRNA, mRNA, and Other. **(D)** Proportional bar graph depicting the region of an mRNA where Sm-sites are predicted. **(E)** Density plot of total transcript lengths categorized by type of Sm-site predicted. **(F)** Density plot of mRNA 3’UTR lengths categorized by the type of Sm-site predicted. See Supplementary Figure 2 for this analysis broken down for each individually identified U1, U2, U5, U7, and noncanonical Sm-site.

Many classes of RNAs were identified to contain Sm-sites, including snRNA, small nucleolar and Cajal-body specific RNAs (sno/scaRNA), long noncoding RNAs (lncRNA) and mRNAs (Figure 1B). Surprisingly, although mRNAs comprise about a third of all annotated transcripts (33% in human, 38% in mouse), as a group, they make up an outsized proportion of predicted canonical (60% in human, 64% in mouse) and noncanonical (36% in human, 45% in mouse) Sm-site-containing RNAs (Figure 1B). Similarly, only about a fifth of all genes are predicted to contain a canonical Sm-site (18% in humans and 19% in mouse, Figure 1C), but a third of all mRNAs are predicted to contain canonical Sm-sites (33% of human and 33% of mouse mRNAs) suggesting that Sm-sites may be preferentially enriched in mRNAs. The second most-represented group is lncRNAs, making up 26% of human and 16% of mouse canonical Sm-sites (Figure 1B). Noncanonical Sm-sites are predicted to occur in over a third of human (38%) and nearly half of mouse (46%) annotated genes. Noncanonical Sm-sites appear most frequently in longer RNAs—lncRNA and mRNA—in similar proportions (∼45% in human and 54% in mouse) (Figure 1C). Only 15% of snRNAs are predicted to contain an Sm-site. This is not surprising given nearly 71% of annotated human snRNAs are U6/U6atac, which does not have an Sm-site and does not receive an Sm-ring (91) (Supplementary Figure 1, Supplementary File 1-2). Of the remaining human snRNAs, only 46% are predicted to contain an Sm-site, suggesting many annotated snRNAs may not be functional, may not be Sm-class, or their Sm-site sequence remains to be defined (Supplementary Figure 1, Supplementary File 1-2). In all biotypes except snRNAs, U2-type Sm-sites are the most abundant, particularly in lncRNA and mRNAs (Supplementary Figure 2A, 2B). U7-type sites are primarily identified in U7 snRNAs and only identified in a handful of other transcripts, supporting our argument to exclude these from our canonical Sm-site definition. These data suggest that many RNAs contain features of snRNAs, and that many of these RNAs are mRNAs.

To assess whether Sm-site prediction is a function of transcript length, we stratified RNAs by the presence (≥ 1 canonical or noncanonical) or absence of an Sm-site, and plotted them by their transcript length (Figure 1E). Interestingly, canonical Sm-sites are found in longer transcripts (median transcript length, human = 3,092 nt; mouse = 2,936 nt) than those that have noncanonical Sm-sites (human = 1,281 nt; mouse = 1,531 nt) or that do not have Sm-sites (human = 466 nt; mouse = 407 nt) (Figure 1E). The frequency of canonical Sm-sites is moderately correlated with transcript length (human: Kendell τ = 0.35, *p* < 2.2 x 10^-16^; mouse: Kendell τ = 0.35, *p* < 2.2 x 10^-16^) (Supplementary Figure 3A). The frequency of noncanonical Sm-sites is more strongly correlated with transcript length (human: Kendell τ = 0.54, *p* < 2.2 x 10^-16^; mouse: Kendell τ = 0.61, *p* < 2.2 x 10^-16^) (Supplementary Figure 3B). The finding that the most stringent predicted Sm-sites are more commonly found in longer transcripts than the more promiscuous noncanonical sites suggests that canonical Sm-sites are a *bona fide* feature of these long transcripts and their detection is not simply a function of RNA length.

mRNAs are known to contain regulatory elements within their 5’ and 3’ untranslated regions (UTR) (92,93). To test if Sm-sites are more prevalent in one or more of these regions, Sm-sites identified in mRNAs were binned by their location: 5’UTR, Coding Sequence (CDS), and 3’UTR (Figure 2A). Additionally, Sm-sites identified in exonic sequences (Exon Mismatch) or other sequences (N/A) that do not clearly match the reference genome were also included (Figure 1D). We observe that nearly three-fourths of Sm-sites, irrespective of the type of Sm-site, are found in the 3’UTRs of mRNAs (Figure 1D and Supplementary Figure 2C). An estimate of the density of Sm-sites showed that canonical Sm-sites are three times more likely to be detected in an 3’UTR than in any other mRNA region (in units of sites/million bases, human: 5’UTR 60.2 CDS 87.7, 3’UTR 291.4; mouse 5’UTR 70.3, CDS 90.7, 3’UTR 260.0). Not only are canonical Sm-sites more likely to be found in mRNA 3’UTRs, but they are also more commonly detected in long 3’UTRs (human: n = 7,717 mRNAs, median 2,181 nt; mouse: n = 7,121 mRNAs, median 1,478 nt) (Figure 1F) whereas noncanonical Sm-sites are found in shorter 3’UTRs (human: n = 9,529 mRNAs, median 794 nt; mouse: 11,878 mRNAs, median 645 nt). These observations further strengthen the argument that canonical Sm-sites are preferentially found in mRNA 3’UTRs. Lastly, like overall transcript lengths, 3’UTR length is moderately correlated with the number of canonical sites (human: Kendell τ = 0.44, *p* < 2.2 x 10^-16^ ; mouse: Kendell τ = 0.36, *p* < 2.2 x 10^-16^) (Supplementary Figure 3C) whereas the correlation between 3’UTR length and noncanonical Sm-site frequency is stronger (human: Kendell τ = 0.62, *p* < 2.2 x 10^-16^ ; mouse: Kendell τ = 0.60, *p* < 2.2 x 10^-16^) (Supplementary Figure 3D). Taken with the aforementioned results, these data suggest that Sm-sites are more commonly found in the 3’UTRs of longer mRNAs over other RNAs.

**Figure 2:**
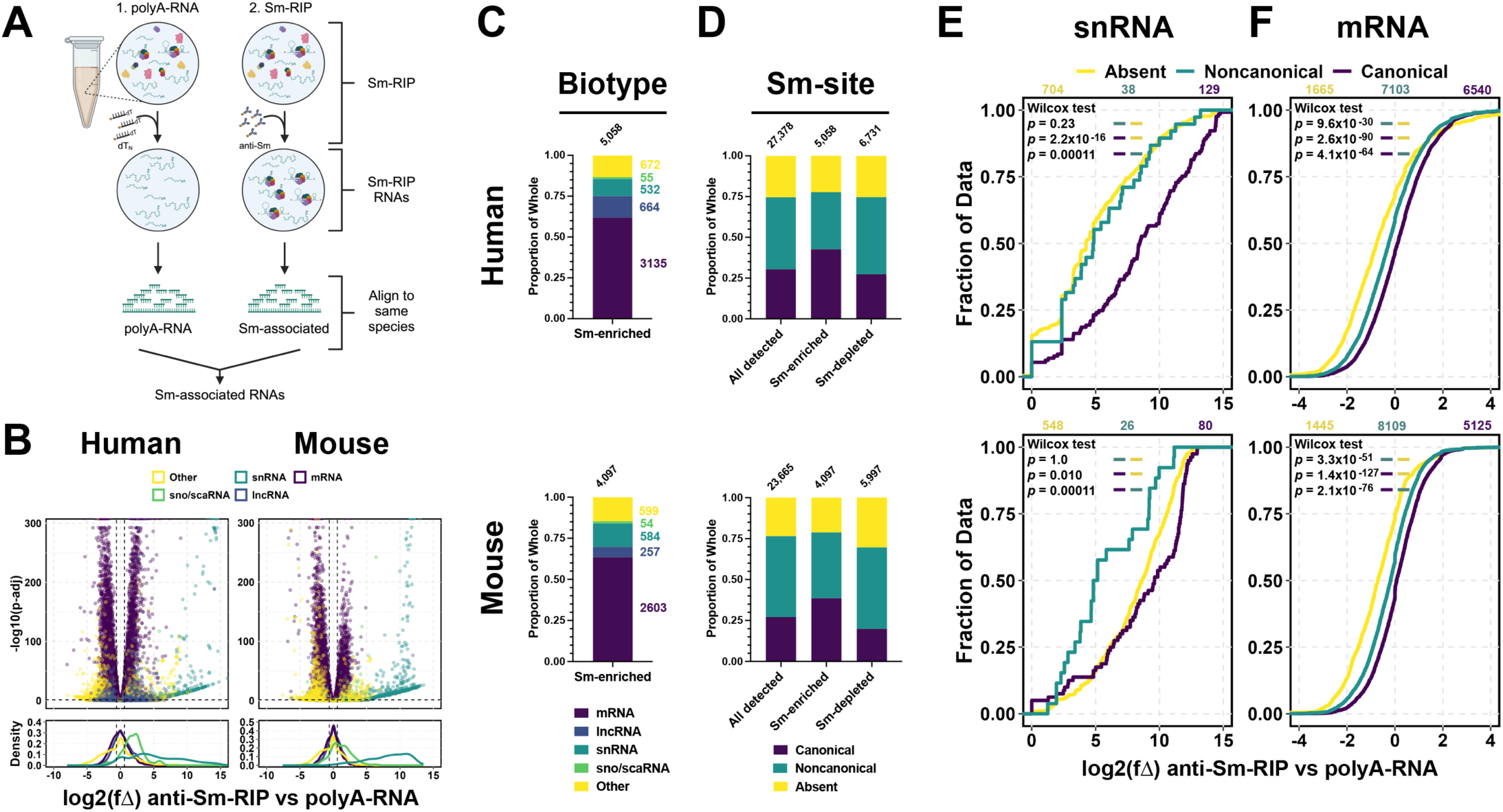
Sm-site containing RNAs are specifically enriched with Sm-proteins. **(A)** Schematic representation of experimental conditions. **(B)** Volcano plot and accompanying density plot indicating the different types of RNAs that are enriched in an anti-Sm-RIP experiment when compared to the polyA-RNA enriched transcriptome. X-axis for volcano and density plots are shared—log2 fold change (log2(fΔ)) of anti-Sm-RIP vs polyA-RNA. For all bar graphs, at the top are the number of genes represented within the bar. Values, to the side of the bar are the number of genes contained within the portion corresponding to the represented color. For **C-F**, **(top)** human analysis and **(bottom)** mouse analysis. U7 Sm-site-containing RNAs were removed from analsysis as this site is both infrequent and receives a specialized Sm-ring differentiating it from U1, U2, and U5 type Sm-sites. **(C)** Proportional bar graph of RNA biotypes enriched (log2(fΔ) > 0.6, *padj* < 0.05) in anti-Sm-RIP vs the polyA-RNA transcriptome. **(D)** Proportional bar graph giving a breakdown of the Sm-sites. **All** indicates the total different transcripts mapped in the sequenced library, **Sm-enriched** are those transcripts enriched (log2(fΔ) > 0.6 *padj* < 0.05) in the anti-Sm-RIP. **Un-enriched** are those transcripts that were not enriched (log2(fΔ) < -0.6, *padj* < 0.05) in the anti-Sm-RIP. For **E-F**, Cumulative distribution plots for given RNA biotypes, plotting the log2 fold change (log2(fΔ)) by increasing value. log2(fΔ) was calculated between the anti-Sm-RIP vs polyA-RNA conditions. **(E)** snRNA, **(F)** mRNA. Colors designate whether an Sm-site is **Absent** (yellow), **Noncanonical** (green), or **Canonical** (purple). Values above CDFs indicate the number of genes plotted for each condition in color. *P-values* generated from Wilcox tests for the left color being greater than the right color are provided in the upper lefthand corner of the graphs.

### Biochemical identification of RNAs that associate with Sm-proteins in an ATP-dependent manner

The prevalence of Sm-sites in mRNAs and other non-snRNAs suggest that Sm-protein rings may associate with RNAs beyond snRNAs. Although Lu *et al* (11) previously reported such Sm-protein:mRNA association, it is not known if this interaction is dependent on Sm-sites. To identify RNAs that associate with Sm-proteins at equilibrium in human and mouse cell cytoplasmic extracts, we performed anti-Sm (Y12) RNA immunoprecipitation followed by high-throughput sequencing (RIP-Seq) from cytoplasmic extracts of human S3 induced neuronal progenitor cells (iNPC) and immortalized mouse neuronal stem cells (NSC-34). Previous work from multiple laboratories has shown that Sm-protein rings can be assembled onto the Sm-site of synthetic, labelled U snRNAs by the SMN complex in an ATP-dependent manner in cytoplasmic cell extracts (20,22,27,48). Such newly assembled U snRNPs can be enriched and quantified by anti-Sm immunoprecipitation in a stringent manner that selects for RNAs that assemble a high-salt and heparin-resistant Sm-ring (20,22,27,48) (Supplementary Figure 4A). We also sought to determine if Sm-protein rings can be assembled on RNAs other than snRNAs. To this end, a library of polyA-enriched RNA (polyA-RNA) was supplemented into cytoplasmic cell extracts to be used as a substrate (instead of labelled U snRNA) in modified Sm-ring assembly reactions, which were performed in the presence or absence of ATP (ATP+ or ATP-, respectively) (Supplementary Figure 4B). The cytoplasmic cell extracts and polyA-RNA were prepared from human iNPCs and mouse NSC-34 cell lines, and the assembly reactions were performed in a cross-species manner, i.e. human polyA-RNA incubated with mouse extract and vice versa. This allowed us to identify newly assembled Sm-RNPs from the Sm-RNPs already present in the cytoplasmic extract (Supplementary Figure 4B). All Sm-RNPs were then identified via anti-Sm RIP-Seq. Finally, the input polyA-RNA libraries from the two cell types were also sequenced to serve as a baseline for the enrichment analysis (Supplementary Figure 4B).

To ensure robust, reproducible and equivalent Sm-ring assembly across experimental conditions, several key controls were performed. First, snRNP assembly capacity was normalized between cytoplasmic extracts used for the RIP-seq experiments by empirically determining the amounts of extracts that yield equivalent Sm-ring assembly on a biotinylated *in vitro* transcribed human U4 snRNA (Supplementary Figure 5A). Second, multiple anti-Sm (Y12) antibodies were tested to determine the source and amount required for saturation of capture (Supplementary Figure 5B). Third, polyA-RNA after oligo-dT enrichment was confirmed to be depleted of ribosomal RNAs (Supplementary Figure 5C). Fourth, reactions were scaled up to contain 400 µg total protein in the cytoplasmic extracts and 4 µg of polyA-RNA to provide sufficient yields for sequencing library preparation. Finally, sequencing libraries were generated and sequenced for three biological replicates for each condition presented in Supplementary Figure 4B.

Alignment of sequencing reads to both the human GRCh38 and mouse GRCm39 genome assemblies using STAR (83) showed that, reads from samples consisting of only human or mouse RNAs align primarily to their corresponding genomes, but poorly to the cross-species genome, as expected (Supplementary Figure 5D). Furthermore, samples containing RNA from both species show greater read mapping to the genome corresponding to the input polyA-RNA (Supplementary Figure 5D), suggestive of higher abundance of new Sm:RNA interactions. Less than 10% of genes align to both human and mouse genomes (Supplementary Figure 5D). These genomically ambiguous RNAs were removed from further analysis. Principal component analysis shows that replicates for each condition are very tightly clustered (Supplementary Figure 5E). The largest variation between samples is conferred by species and the second largest variation is between the input polyA-RNA condition and Sm-RIP conditions (Supplementary Figure 5E). All modified Sm-ring assembly reaction conditions cluster together, most closely resembling the Sm-RIP-Seq samples isolated from extracts of the same species the samples were aligned to (Supplementary Figure 5E). Thus, our libraries are reproducible and specific, and the data indicates robust signal for polyA-RNAs receiving a newly assembled Sm-ring. We will first discuss what RNAs are enriched with Sm-proteins from unsupplemented cytoplasmic extracts and the degree to which the prediction of Sm-sites informs this enrichment. Then we will discuss the degree to which we detect newly assembled Sm-RNPs on exogenously supplemented polyA-RNAs.

### Many RNAs including mRNAs with predicted Sm-sites associate with Sm-proteins

Many different species, or biotypes, of RNA have been shown to co-immunoprecipitate with Sm-proteins (Supplementary Figure 6A) (11). However, it was not determined if the enriched RNAs contained features of U snRNAs or whether Sm-rings were assembled upon these Sm-associating RNAs (11). Using our Sm-site prediction method, we determined 28% of the RNAs identified by Lu *et al* were predicted to contain a canonical Sm-site and 37% were predicted to contain noncanonical Sm-sites (Supplementary Figure 6B). The Sm-enriched dataset provided by Lu *et al* comprises a small list of genes, determined by comparing an anti-Sm-RIP to an isotype control RIP in HeLa total cell extracts. To expand upon this study and provide the transcriptome context for the RIP, we investigated the type of RNAs enriched in Sm-RIP-Seq samples from NCS-34 and S3 iNPC cytoplasmic cell extracts (without addition of exogenous polyA-RNA and ATP) and compared them to the polyA-enriched RNAs from the same cell line (Figure 2A).

Many RNA biotypes, other than snRNAs, are represented in the Sm-enriched fraction (log_2_(fΔ) > 0.6, *p* < 0.05) (Figure 2B-C). mRNAs comprise 62% of human and 64% of mouse RNAs detected in Sm-RIPs, with the second largest biotypes being lncRNAs and the catch-all “Other” category (Figure 2C). The proportion of RNAs with a canonical Sm-site is the highest among the Sm-enriched RNAs as compared to RNAs that are not enriched (Un-enriched) and all the total RNAs detected in the analysis (Figure 2D). On the other hand, the proportion of noncanonical Sm-site containing RNAs is reduced in the anti-Sm-RIP condition, suggesting that noncanonical sites are less deterministic for RNA association with Sm-proteins. Indeed, the proportion of noncanonical sites within the un-enriched gene list is similar to all RNAs detected in both humans and mouse (Figure 2D). Furthermore, RNAs predicted to have a canonical Sm-site are more enriched as compared to those predicted to have a noncanonical Sm-site (Supplementary Figure 7A), further suggesting the noncanonical condition is permissive and likely comprises RNAs that exhibit reduced specificity in Sm-protein association. Surprisingly, only 22% of the RNAs identified by Lu *et al* are shared in our anti-Sm-RIP dataset, likely due to the differences in cell type and conditions of the comparisons.

To further test if Sm-site prediction specifically informs Sm-protein-RNA association, we examined Sm-RIP enrichment of RNAs with and without Sm-sites from different RNA biotypes (Figure 2E-F, Supplementary Figure 7B-C). The presence of an Sm-site in sno/scaRNAs, snRNAs and mRNAs (and less so in the case of lncRNAs) leads to enrichment with Sm-proteins. For snRNAs and mRNAs, presence of canonical Sm-site as compared to a noncanonical site leads to their higher enrichment (Figure 2E-F). Surprisingly, mRNAs are the largest contributor to the Sm-site containing RNAs enriched in Figure 2F (compare gene numbers in Figure 2E-F, Supplementary Figure 7B-C). Although sample sizes for snRNAs and sno/scaRNAs are low, perhaps due to poor annotation of these biotypes within the Refseq and Gencode databases, human snRNA—which are better annotated than their mouse counterparts—do indicate that Sm-site presence is directly correlated with RNA enrichment with Sm-proteins (Figure 2E). Surprisingly, mouse snRNAs devoid of any detectable Sm-sites are nearly as enriched as canonical Sm-site containing snRNAs, suggesting the Sm-site definition used may not fully represent mouse snRNAs.

Importantly, all types of Sm-sites predict further enrichment in an Sm-RIP with U2 and U7-type Sm-site-containing RNAs exhibiting the largest shift in comparison to RNAs lacking Sm-sites (Supplementary Figure 8). Furthermore, the frequency of Sm-sites within an RNA is directly correlated to RNA enrichment with Sm-proteins (Supplementary Figure 9), suggesting specificity of Sm-proteins for the Sm-site. RNAs that contain 3 or more predicted canonical Sm-sites are further enriched in Sm-RIPs than those that have only 2, 1, or no predicted Sm-sites (Supplementary Figure 9A). Additionally, the increased frequency of noncanonical Sm-sites also enhances enrichment in Sm-RIPs of transcripts that contain at least one canonical Sm-site (Supplementary Figure 9B) or that do not contain a canoncial Sm-site (Supplementary Figure 9C). Transcript length and 3’UTR length does not correlate with Sm-protein enrichment, suggesting that enrichment is due to the presence of the predicted Sm-site and not simply dictated by length of the RNA (Supplementary Figure 10A and 10B, respectively). Together, these data support a conclusion that RNAs predicted to contain Sm-sites are specifically and preferentially associated with Sm-proteins in human and mouse extracts under physiological conditions.

Based on our findings that RNAs containing canonical (and noncanonical) Sm-sites are enriched in anti-Sm-RIP-Seq experiments, we predicted that such RNAs will also show preferential binding to Sm-ring components in the published SmB/B’ CLIP-Seq dataset (94). Indeed, as compared to RNAs lacking Sm-sites, the RNAs predicted to contain Sm-sites are also enriched in the published SmB/B’ CLIP-Seq dataset (Supplementary Figure 6D-G) (94). Importantly, among the highest quartile of RNAs enriched in the SmB/B’ CLIP-Seq dataset, 578 transcripts are shared with our Sm-RIP-Seq datasets (Supplementary Figure 6H). 97% of these shared transcripts are mRNAs and 97% are predicted to contain an Sm-site (canonical + noncanonical) (Supplementary Figure 6H-J). Thus, we conclude that Sm-ring assembly can occur on Sm-sites contained in RNAs other than snRNAs, including mRNAs.

### Sm-site containing polyadenylated RNAs associate with Sm-proteins in an ATP-dependent manner

Sm-protein ring assembly is an ATP-dependent process inside cells as well as in cell extracts (20,48). To identify the most likely candidates that undergo Sm-protein ring assembly, we compared RNAs enriched in Sm-RIPs from extracts supplemented with both polyA-RNA and ATP (ATP+) to those without the addition of ATP (ATP-) (Figure 3A). Notably, in these comparisons, sequencing reads were aligned to the genome reference corresponding to the species from which the polyA-RNA was isolated. For example, when mouse extract was supplemented with human polyA-RNA + ATP and compared to mouse extract supplemented with human polyA-RNA only, reads were aligned to the human genome (Figure 3A). The capture of several transcripts is significantly different (log_2_(fΔ) > 0.6, *p* < 0.05 or log_2_(fΔ) > -0.6, *p* < 0.05) between ATP+ and ATP-conditions, with 485 human and 883 mouse RNAs significantly enriched, and 63 human and 315 mouse RNAs significantly depleted (Figure 3B-C). The difference in the number of transcripts enriched following ATP-addition between mouse and human is likely contributed by the 1.8 fold higher amount of mouse extract (438 µg) used in the RIP-Seq experiments as compared to the human extract (250 µg), which was done to accommodate the reduced capacity of mouse extracts to assemble Sm-rings (Supplementary Figure 5A). Strikingly, the largest fraction of the ATP-induced Sm-enriched RNAs are protein coding (49% for human and 70% for mouse) (Figure 3B). snRNAs comprise 25% of the human and 11% of the mouse ATP-induced Sm-enriched RNAs, possibly due to their depletion in the polyA-RNA libraries. A vast majority of enriched human (87%) and mouse (99%) mRNAs contain a predicted Sm-site (Figure 3D) and Sm-site containing mRNAs are globally enriched in extracts supplemented with ATP (Figure 3E). Akin to the anti-Sm-RIP in unsupplemented extracts (Figure 2D), noncanonical Sm-sites are proportionally reduced in the ATP and polyA-RNA supplemented condition, suggesting the definition of a noncanonical Sm-site used is too permissive. Addition of ATP also induces Sm-enrichment of Sm-site containing sno/scaRNA, snRNAs, and lncRNAs (Supplementary Figure 11B-D). Further, mRNAs with canonical Sm-sites are significantly more enriched upon the addition of ATP than those with noncanonical or absent Sm-sites, suggesting Sm-protein rings are specifically assembled on canonical Sm-site-containing polyA-RNAs (Figure 3D-E). The addition of ATP also enriches U2 and U5-type Sm-site containing mRNAs more than U1, U7, or noncanonical Sm-site containing mRNAs (Supplementary Figure 12C). The frequency of Sm-sites identified within mRNAs is predictive of further enrichment upon the addition of ATP (Figure 3F, Supplementary Figure 13) whereas transcript and 3’UTR length does not correlate with ATP-inducible enrichment (Supplementary Figure 14 A and B, respectively), underscoring that enrichment is due to the presence of *bona fide* Sm-sites. Together, hallmarks of Sm-ring assembly—Sm-site and ATP-dependence—correlate with enrichment of specific RNAs in *in vitro* Sm-ring assembly reactions, suggesting that Sm-protein rings can be more broadly assembled on Sm-site containing RNAs.

**Figure 3.**
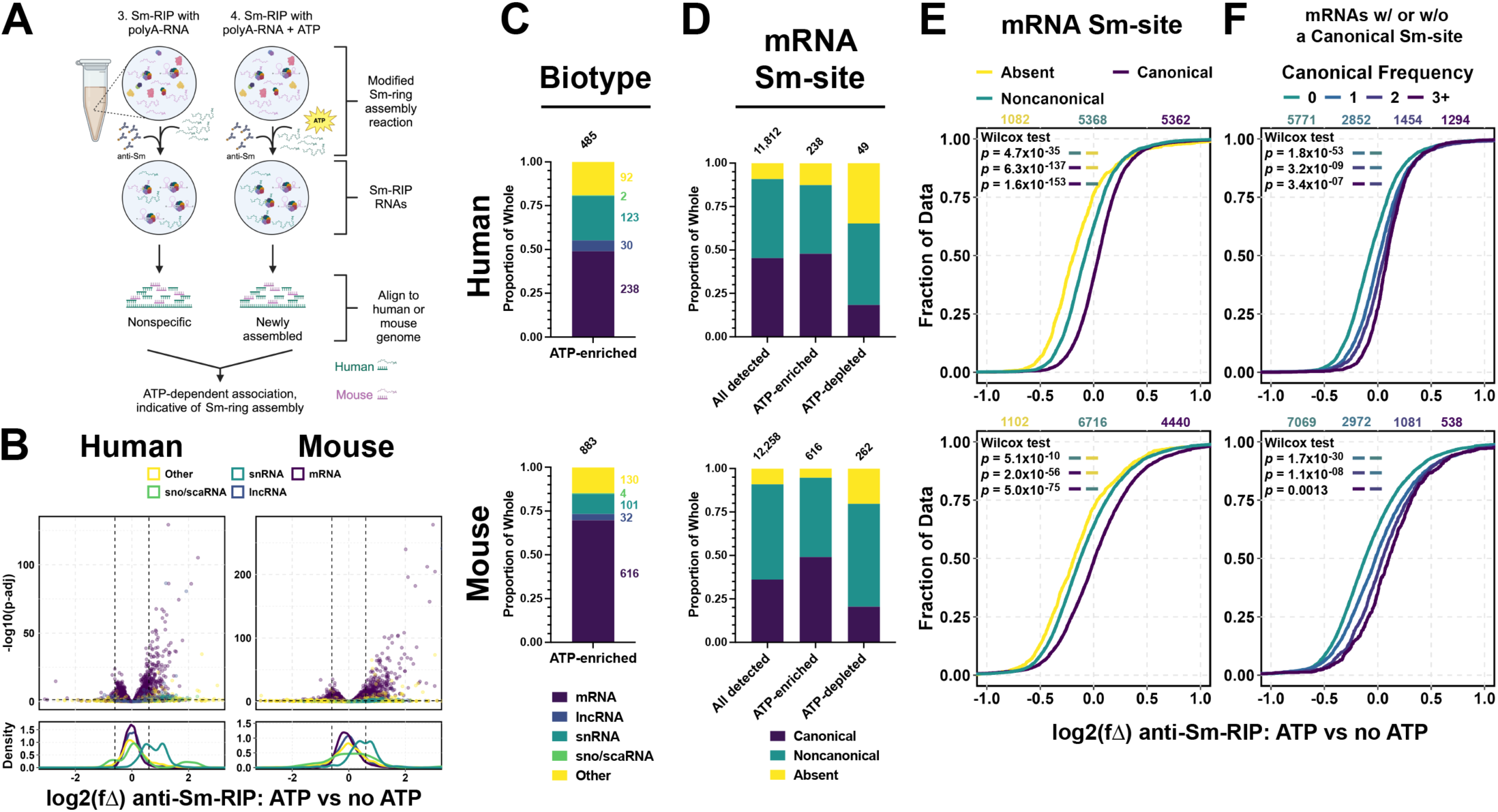
Sm-site containing mRNAs are specifically enriched in anti-Sm-RIPs in an ATP-dependent manner. **(A)** Schematic representation of the experiment. **(B)** Volcano plot and accompanying density plot indicating the different types of RNAs that are enriched in a modified Sm-RIP experiment supplemented with polyA-RNA and ATP vs. a modified Sm-RIP supplemented polyA-RNA only. X-axis for volcano and density plots are shared—log2 fold change (log2(fΔ)) of anti-Sm-RIP: ATP vs no ATP. For **(C-E)**, **(top)** for human analysis and **(bottom)** for mouse analysis. For all bar graphs, top numbers are the number of genes represented within the bar and values to the side correspond to the number of genes represented in each color of the graph. U7 Sm-site-containing RNAs were removed from analsysis. **(C)** Proportional bar graph of RNA biotypes enriched (log2(fΔ) > 0.6, *padj* < 0.05) in anti-Sm-RIP supplemented with ATP vs anti-Sm-RIP not supplemented with ATP. **(D)** Proportional bar graph giving a breakdown of types of Sm-sites in mRNAs represented in the analysis (All), represented in those mRNAs enriched following anti-Sm-RIP (ATP-enriched, log2(fΔ) > 0.6, *padj* < 0.05), and those not-enriched (Un-enriched, log2(fΔ) < -0.6, *padj* < 0.05). For **(D-E)**, Values above plots are the number of genes plotted for each group in color. *p-values* for Wilcox tests using the alternative that the left color is greater than the right color are provided in the upper-lefthand corner of the plots. **(D)** Cumulative distribution plot of only mRNAs, delineating by Sm-site prediction—**Absent** (yellow), **Noncanonical** (green), **Canonical** (purple). **(E)** Cumulative distribution plot delineating the frequency of canonical Sm-site prediction in an mRNA transcript by log2 fold change in anti-Sm-RIP supplemented with polyA-RNA and ATP vs anti-Sm-RIPs supplemented only with polyA-RNA. mRNAs predicted to solely contain noncanonical Sm-sites are not plotted in this graph.

### Sm-proteins can be directly assembled on *in vitro*-transcribed Sm-site containing mRNA 3’UTRs

Though Sm-site containing RNAs are associated with Sm-proteins and are specifically enriched with Sm-proteins when induced by the addition of ATP, this does not indicate Sm-protein rings specifically assemble at the predicted Sm-sites within these mRNAs. Therefore, to identify the best candidate mRNAs to directly test Sm-protein ring assembly on Sm-sites of representative polyA-RNAs, we compared different experimental conditions indicated in Supplementary Figure 4B and applied a series of filtering steps (Figure 4A). Top candidate RNAs were expected to be enriched upon the addition of ATP, genomically unambiguous, enriched over the polyA-RNA input library, and to associate with Sm-proteins. Enrichment within a particular experimental comparison was defined as a > 1.5-fold change with an adjusted *p*-value less than 0.05 (log_2_(fΔ) > 0.6, *p* < 0.05). 59 human and 66 mouse candidate RNAs satisfy the full set of conditions (Figure 4A, Supplementary File 2). Of these candidate RNAs, the majority are mRNAs (69% in human and 88% in mouse) (Figure 4B). Indeed, 71% of human and 86% of mouse candidate RNAs contain an Sm-site (Figure 4C). This data further underscore that Sm-site containing polyA-RNAs, particularly mRNAs, are robust candidates to accept an Sm-protein ring in an ATP-dependent manner.

**Figure 4.**
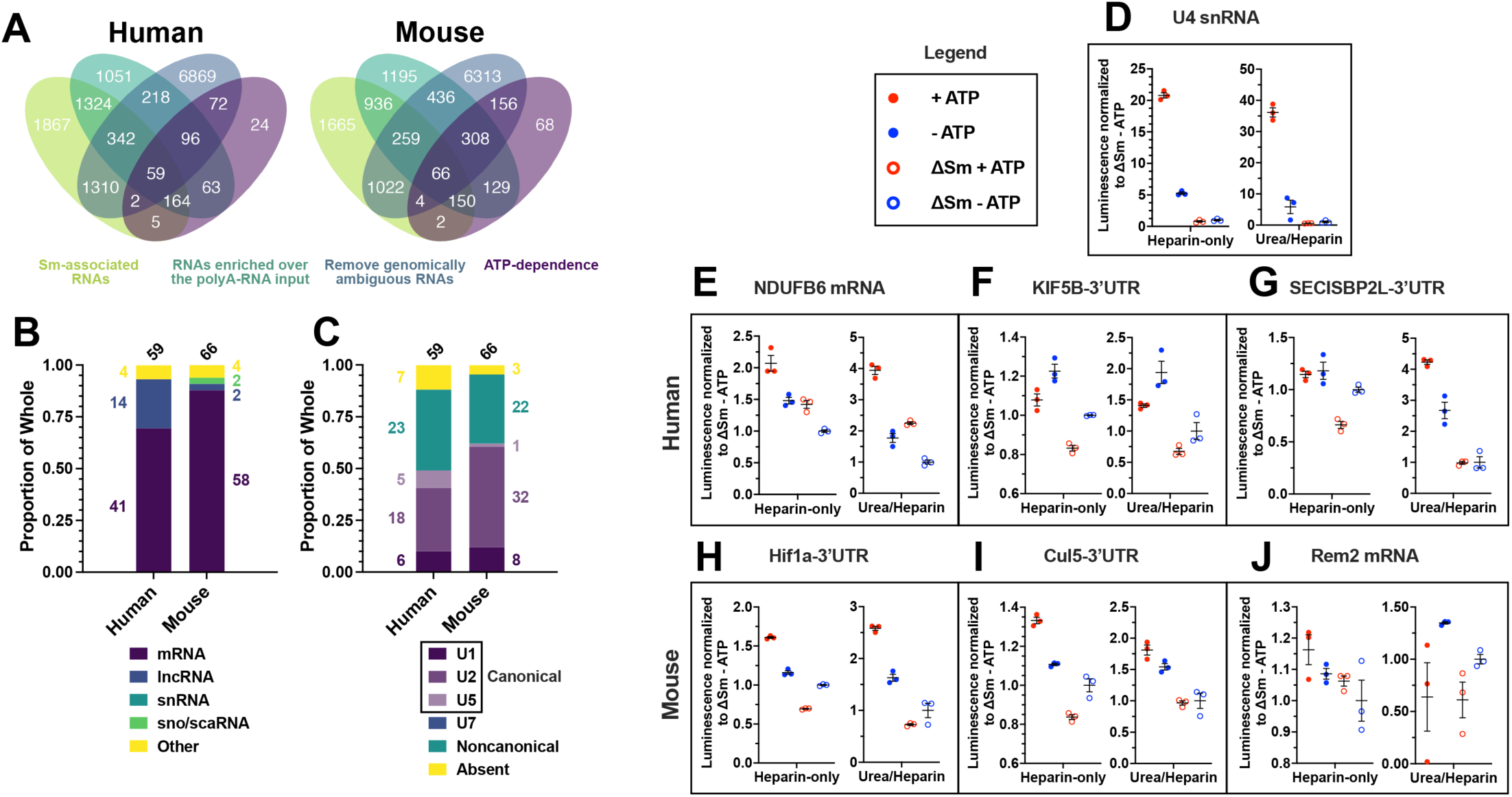
Sm-protein rings are directly assembled on mRNAs in an ATP and Sm-site dependent manner. **(A)** Venn diagrams depicting the comparisons made to select candidate mRNAs to test for Sm-ring assembly directly. Each ellipse is representative of gene products enriched (log2(fΔ) > 0.6, *padj* < 0.05) in the noted comparison. The ∼60 genes in the center are enriched in each comparison, comprising the candidate list. For **(B-C)**, values above bars indicate the number of genes contributing to the plots. Numbers in color to the sides of bars indicate the number of genes contributing to the specified group within the bar. **(B)** Proportional bar graph of the RNA biotypes for gene products enriched (log2(fΔ) > 0.6, *padj* < 0.05) within each comparison depicted in **(A)**. **(C)** Proportional bar graph giving a breakdown of types of Sm-sites predicted in RNA enriched (log2(fΔ) > 0.6, *padj* < 0.05) in all comparisons, depicted in **(A)**. **(D-J)** Luminescence results from detection of *in vitro* transcribed, biotin-labelled human U4 snRNA **(D)**, mRNAs (NDUFB6 **(E)** and Rem2 **(J)**), and mRNA 3’UTRs (KIF5B **(F)**, SECISBP2L **(G)**, Hif1a **(H)**, and Cul5 **(I)**) enriched following anti-Sm-RIP. Four conditions were performed for each RNA: (solid red dot) cytoplasmic cell extract supplemented with wild-type RNA and ATP, (solid blue dot) cytoplasmic cell extract supplemented with wild-type RNA but not with ATP, (open red circle) cytoplasmic cell extract supplemented with ATP and RNA mutated to remove the Sm-site sequence, and (open blue circle) cytoplasmic cell extract supplemented with RNA mutated to remove the Sm-ste sequence but not ATP. Right graphs are the results obtained following 15 min treatment with 2M urea and 5 mg/mL heparin, followed by immunoprecipitation in 2 mg/mL heparin, RSB-500 + 0.1% NP-40 and washed 8 times with RSB-500 + 0.1% NP-40. Raw luminescence values were normalized to the ΔSm -ATP condition for each individual experiment. All shown RNAs have a canonical Sm-site except Rem2. Left graphs are the results obtained by performing the immunoprecipitation in 2 mg/mL heparin, RSB-500 + 0.1% NP-40 and washed 8 times with RSB-500 + 0.1% NP-40. The raw values are plotted in Supplementary Figure 15.

For our *in vitro* validation studies, we focused on candidate mRNAs that 1) contain a canonical Sm-site within the 3’UTR at a similar locus of the mRNA in both mouse and human orthologs, 2) do not have multiple canonical Sm-sites at vastly different locations of the mRNA, and 3) are devoid of noncanonical Sm-sites. Candidate mRNAs and 3’UTRs were *in vitro* transcribed in the presence of biotin-UTP. As controls, Sm-sites, canonical and/or noncanonical were mutated in which all uridines were changed to cytosines or alternating cytosines (i.e.: from 5’-AUUUUUG-3’ to 5’-ACCCCCG-3’ or 5’-ACUCUCG-3’). Both candidate RNAs containing and lacking their respective Sm-sites were then assayed for Sm-protein ring assembly (48), testing for ATP-inducibility and Sm-site dependence of assembly, mimicking that of a similarly labeled human U4 snRNA.

Some mRNAs and 3’UTRs containing canonical Sm-sites show both an ATP and Sm-site dependence in Sm-protein ring assembly under conditions previously shown to select for RNAs with assembled Sm-rings (500 mM NaCl and 2 mg/mL heparin), as described by Wan *et al* (Figure 4D-J left, raw luminescence in Supplementary Figure 15A-G left). These include the human NDUFB6 mRNA, and the mouse Hif1a and Cul5 3’UTRs (Figure 4E, 4H-I left, Supplementary Figure 15B, 15E-F left). Despite exhibiting the highest luminescence, the human KIF5B-3’UTR only shows an Sm-site dependence and not an additional ATP-inducible increase in luminescence signal (Figure 4F left, Supplementary Figure 15C left). Furthermore, the only noncanonical Sm-site containing mRNA tested, mouse Rem2, showed only a modest ATP and Sm-site dependence (Figure 4J left, Supplementary Figure 15G left).

To test Sm-ring assembly under even more stringent conditions (29,50), we incubated the reactions with 2 M urea and 5 mg/mL heparin, followed by washes in 500 mM NaCl and 2 mg/mL heparin (Figure 4D-J right, Supplementary Figure 15A-G right). Under these more stringent conditions, the maximal luminescence signal detected was reduced, but the signal between the ATP with intact Sm-site condition from the other conditions was widened, indicating a more stable and specific Sm-ring assembly (Figure 4D-J right, Supplementary Figure 15A-G right). Urea/heparin treatment confirmed specific Sm-protein ring assembly on human NDUFB6 mRNA, on mouse Hif1A and Cul5 3’UTRs, and additionally on the SECISBP2L 3’UTR (Figure 4E, 4G-I right, Supplementary Figure 15B, 15D-F right). Sm-site dependence but a lack of ATP-dependence was likewise confirmed for the human KIF5B 3’UTR (Figure 4F right, Supplementary Figure 15C right). The mouse Rem2 mRNA showed no increase in luminescence signal following urea/heparin treatment, indicating that Sm-ring assembly on this candidate mRNA is not specific (Figure 4J right, Supplementary Figure 15G right). Lastly, stable Sm-ring assembly on NDUFB6 mRNA and Hif1a-3’UTR can also be detected via immunoprecipitation using an anti-SmB/B’ specific antibody (Supplementary Figure 16). These data confirm that Sm-protein rings can be assembled on RNAs other than snRNAs if they contain a canonical Sm-site.

### Sm-site containing mRNAs have a reduced abundance in SMA animal and cell models

A reduced capacity to assemble U snRNPs has been shown to result in reduced stability of U snRNAs, thus we hypothesized that this may also be the case for other RNAs predicted to contain Sm-sites. We mined total RNA-seq datasets from four studies (95–98) in three mouse models of SMA (99–101) to determine if there was a correlation between the abundance of mRNAs containing Sm-sites and the SMA condition. In this analysis, we included the three RNA-seq datasets where the control and disease samples differentiate along the first component of a principal component analysis (PC1): mouse embryonic stem cell derived motor neurons from severe SMA (*Smn^-/-^;tg89SMN2^+/+^*) mice (96), and the spinal cord and liver datasets generated from post-natal day 5 ‘Taiwanese’ SMA (*Smn^-/-^;SMN2^+/-^*) mice (97) (Supplementary Figure 17A-C). We find that, canonical Sm-site containing mRNAs are globally reduced in abundance versus mRNAs lacking Sm-sites in severe SMA mouse embryonic stem cells differentiated to motor neuron fates (Figure 5A, top). Furthermore, this effect is shared, though milder, in spinal cord samples generated from post-natal day 5 ‘Taiwanese’ SMA mice (Figure 5A, middle). Though significantly reduced in abundance, the shift is less apparent for Sm-site containing mRNAs in the liver (Figure 5A, bottom), possibly suggesting cell type specificity. Importantly, mRNAs enriched in our Sm-RIP experiments mimick the reductions seen for Sm-site containing mRNAs (Figure 5B). Perhaps surprisingly, only 81 significantly down-regulated (log2fΔ < 0, *padj* < 0.05) genes are shared across the three datasets and nearly all of these are mRNAs (Figure 5C). Of these 81 genes, 95% contain a predicted Sm-site, potentially identifying RNAs particularly susceptible to low levels of SMN protein abundance (Figure 5D). Lastly, the majority of the canonical Sm-site containing mRNAs that are downregulated are also associated with Sm-proteins (Figure 5E). Taken together, these data suggest Sm-site containing RNAs, including lncRNAs and mRNAs, are less stable when SMN is deficient, mimicking the known physiology for U snRNAs and thereby underscoring the relevance for our findings to physiological states in SMA.

**Figure 5.**
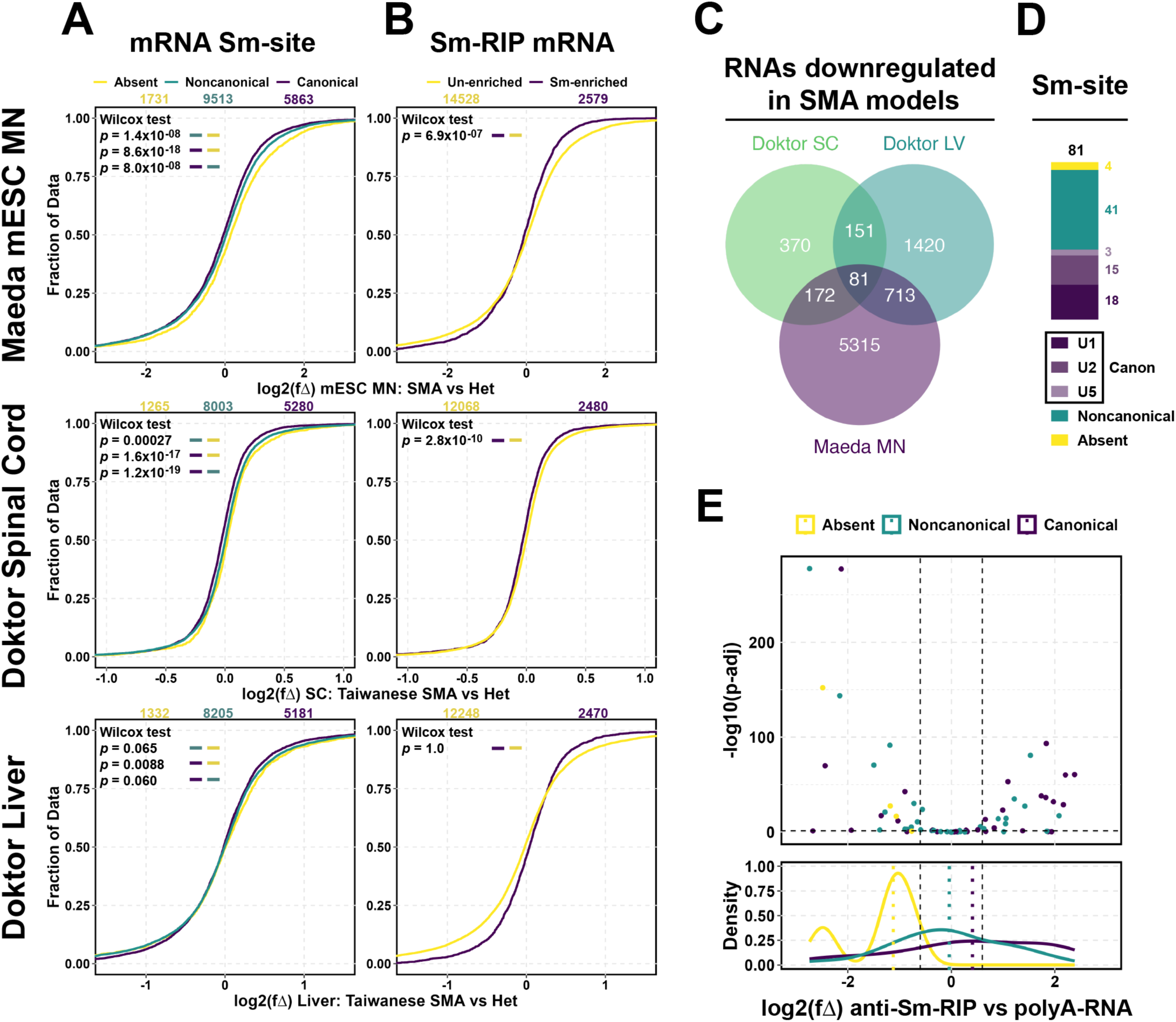
mRNAs predicted to have canonical Sm-sites have a reduced abundance in low SMN conditions. For **(A-B)**, Cumulative distribution function plots of mRNA fold changes in low SMN conditions compared to normal SMN conditions: **(top)** comparison of mESC differentiated motor neurons derived from SMA (*Smn^+/-^*;*SMN2^+/+^*) and normal (*Smn^+/+^*;*SMN2^+/+^*) as published by Maeda *et al PLoS ONE* 2014(96), **(middle)** comparison of spinal cord lysates collected from post-natal day 5 Taiwanese SMA (*Smn^-/-^*;*SMN2^+/+^*) and Taiwanese Het (*Smn^+/-^*;*SMN2^+/+^*) mice as published by Doktor *et al NAR* 2017 (97), and **(bottom)** comparison of liver lysates collected from post-natal day 5 Taiwanese SMA (*Smn^-/-^*;*SMN2^+/+^*) and Taiwanese Het (*Smn^+/-^*;*SMN2^+/+^*) mice as published by Doktor *et al NAR* 2017 (97). Values above plots are the number of genes plotted for each group in color. *P-values* for Wilcox tests using the alternative that the left color is less than the right color are provided in the upper-lefthand corner of the plots. **(A)** Plots delineated by Sm-site prediction—**Absent** (yellow), **Noncanonical** (green), **Canonical** (purple). **(B)** Plots delineated by whether those transcripts were enriched (log2(fΔ) > 0.6, *padj* < 0.05) in anti-Sm-RIPs (**Sm-enriched**, purple) or **Un-enriched** (yellow). **(C)** Venn diagram indicating the shared transcripts with reduced abundance (log2(fΔ) < 0, *padj* < 0.05) when SMN is deficient. **(D)** Proportional bar graph giving a breakdown of types of Sm-sites predicted in RNAs shared between each SMN-deficient dataset. Values above bars indicate the number of genes contributing to the plots. Numbers in color to the sides of bars indicate the number of genes contributing to the specified group within the bar. **(E)** Volcano and density plot for 81 shared genes with reduced abundance in SMA-models as detected in the anti-Sm-RIP vs the polyA-RNA comparison.

### Sm-ring associated mRNAs are strongly correlated with mRNAs enriched in motor neuron soma and alpha-COP vesicles

With the finding that Sm-site containing RNAs, in particular mRNAs, are less abundant in SMA models, we asked whether these RNAs are also identified in cellular processes shown to be affected by SMN-deficiency. One hypothesis in the SMA field is that SMN-deficiency affects mRNA trafficking for translation at specific subcellular locations, particularly down neuronal axons for translation at the axon terminal. To test whether the Sm-site affects this pathway, we analyzed enrichment of Sm-site containing RNAs in axonal versus soma sequencing dataset generated from mouse embryonic stem cell derived motor neurons (102). This dataset determined the axonal transcriptome is distinct from the soma transcriptome, and identifies mRNAs trafficked to be locally translated at axon terminals. In rejection of our initial hypothesis, Sm-site containing mRNAs are massively enriched in motor neuron somas (Figure 6A). Furthermore, Sm-associated mRNAs are also more enriched in motor neuron somas than axons (Figure 6B). These data reject the hypothesis that Sm-sites and Sm-ring assembly are a signal for mRNA axonal trafficking.

**Figure 6.**
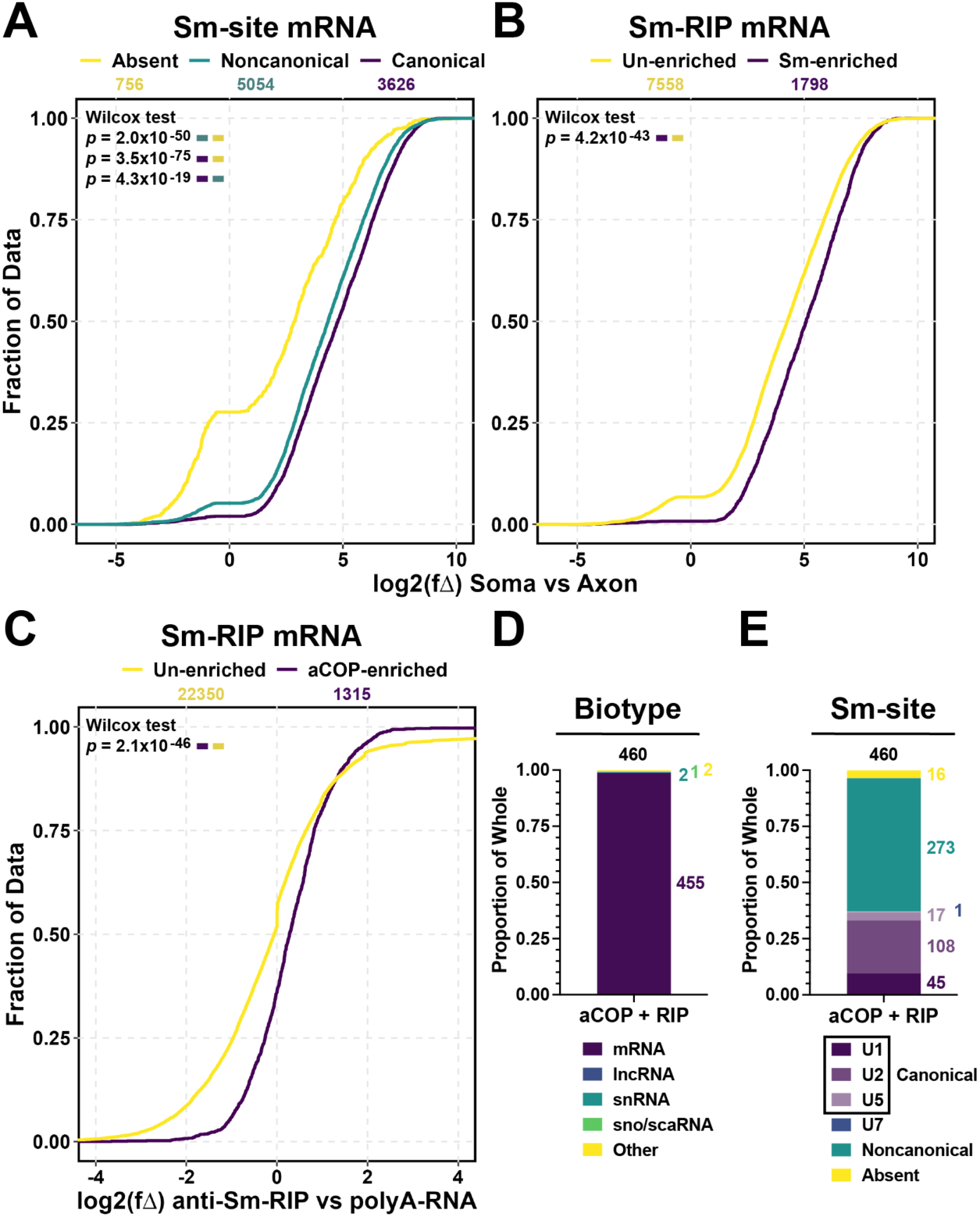
Sm-site containing RNAs are associated with intracellular vessicle trafficking machinery. For **A-C,** Values above plots are the number of genes plotted for each group in color. *P-values* for Wilcox tests using the alternative that the left color is greater than the right color are provided in the upper-lefthand corner of the plots. **(A)** Cumulative distribution function plots comparing log2 fold change between **Soma** or **Axon** specific transcripts as published by Nijssen *et al* (102). Plots are delineated by Sm-site prediction—**Absent** (yellow), **Noncanonical** (green), **Canonical** (purple). **(B)** Cumulative distribution function plots comparing log2 fold change between **Soma** or **Axon** specific transcripts as published by Nijssen *et al* (102). Plots are delineated by whether the transcripts were found to be enriched with Sm-proteins **(Sm-enriched**, purple)—log2(fΔ) ≥ 0.6, *padj* < 0.05 in the anti-Sm-RIP vs polyA-RNA comparison—or **Un-enriched** (yellow) in the anti-Sm-RIP vs polyA-RNA comparison. **(C)** Cumulative distribution function plot comparing log2 fold change in the anti-Sm-RIP vs polyA-RNA comparison, delineating by RNAs previously found to associate with alpha-COP (**aCOP-enriched**, purple), or not (**Un-enriched**, yellow), as found by Todd *et al* (72). For **D-E**, Values above bars indicate the number of genes contributing to the plots. Numbers in color to the sides of bars indicate the number of genes contributing to the specified group within the bar. **(D)** Proportional bar graph giving a breakdown of types of RNAs associated with alpha-COP and physiologically enriched in anti-Sm-RIP (log2(fΔ) ≥ 0.6, *padj* < 0.05). **(E)** Proportional bar graph giving a breakdown of types of Sm-sites predicted in RNAs associated with alpha-COP and physiologically enriched in anti-Sm-RIP (log2(fΔ) ≥ 0.6, *padj* < 0.05).

Additionally, RNAs have been shown to associate with vesicle complexes of the endoplasmic reticulum and Golgi apparatus, mediated through the alpha subunit of the coatamer protein I complex— alpha-COP (72). Not only does SMN and alpha-COP inhabit a similar subcellular locale, but over-expression of alpha-COP was shown to suppress the effects of SMN-deficiency in SMA zebrafish and mouse models (103,104). Previously, another group identified alpha-COP associated RNAs using formaldehyde crosslinking (72). Thus, we asked if these alpha-COP associated RNAs were enriched in our Sm-RIP dataset. Of the 1,519 RNAs enriched in alpha-COP RIPs, 1,315 of them were present in our anti-Sm-RIP dataset and were more globally associated with Sm-proteins (Figure 6C). 460 of the alpha-COP associated RNAs are significantly enriched with Sm-proteins, and almost all of them are mRNAs predicted to contain canonical or noncanonical Sm-sites (Figure 6D-E). These correlations between Sm-sites, Sm-association, and COPI-mediated trafficking of these RNAs suggest a mechanism for alpha-COP mediated suppression of SMN-deficiency in SMA.

## DISCUSSION

### Sm-sites are not restricted to snRNAs and are prevalent in mammalian transcriptomes

Our algorithm searches for Sm-sites modeled on established features of the human U snRNAs (22,27) and identified Sm-sites in all major RNA biotypes, including snRNAs and mRNAs. Multiple observations regarding the detection of Sm-sites in snRNAs indicate the specificity and precision of the algorithm. First, we detect Sm-sites in all snRNAs known to contain these sequences with the exception of U4atac. U4atac is particularly unique in that there are only 8 nucleotides 3’ of the Sm-site which are not predicted to fold (91). As our algorithm requires both a 5’ and 3’ stem loop structure, it would not be expected to identify an Sm-site in U4atac. Second, the algorithm rarely detects Sm-sites in U6 snRNAs (2.6% human, 0% mouse), which lack Sm-sites, are not exported from the nucleus, and do not receive an Sm-ring (105–107). Third, our anti-Sm-RIP-Seq experiments show that human snRNAs detected by the algorithm to contain canonical Sm-sites are enriched over snRNAs predicted to contain a noncanonical Sm-site, or those snRNAs with no prediction for an Sm-site (Figure 2E, top). Thus, Sm-site containing RNAs identified across the transcriptome by our algorithm are highly likely to be targets of Sm-ring assembly.

We included noncanonical Sm-sites in our study to account for the well-known base-substitutions that still result in Sm-association and ring assembly (22). We find that while noncanonical Sm-site containing snRNAs show a lower association with Sm-proteins than canonical Sm-site containing snRNAs (Figure 2C, top), they are further enriched over snRNAs lacking Sm-sites in reactions supplemented with ATP (Supplementary Figure 11C, top). These observations suggest that noncanonical Sm-sites do accept Sm-rings albeit at a lower affinity. The inclusion of noncanonical sites in our search algorithm allowed us to make a comprehensive list of transcripts that exhibit a potential for Sm-ring assembly. Given the increased number of possible sequences that can satisfy the noncanonical Sm-site definition, it is not surprising that nearly twice as many transcripts contain noncanonical Sm-sites than canonical Sm-sites (Figure 1B), noncanonical Sm-sites are found more frequently than canonical Sm-sites (Figure 1A), and noncanonical Sm-site frequency is strongly correlated with transcript and 3’UTR length (Suppleemntary Figure 3). Like snRNAs, mRNAs containing noncanonical Sm-sites exhibit a higher affinity for Sm-proteins than mRNAs lacking an Sm-site, when assembled endogenously and after supplementation with ATP (Figure 2F, 3D, Supplementary Figures 9B-C, 13). Thus, we propose that the noncanonical Sm-sites are likely comprised of both *bona fide* Sm-sites and sites that do not accept Sm-protein ring assembly. In support of this, the Rem2 mRNA, which is predicted to contain only noncanonical Sm-sites, does not show an ATP-dependent or Sm-site dependent association with Sm-proteins (Figure 4J, Supplementary Figure 15G).

A large fraction of annotated snRNAs are not predicted to contain an Sm-site. These include U6/U6atac snRNA that comprise approximately 71% of the human annotated snRNAs. Of the remaining human spliceosomal and U7 snRNAs, 54% are not predicted to contain an Sm-site (Figure 1C, Supplementary Figure 1 top). Our data suggests that human snRNAs lacking an Sm-site likely do not receive an Sm-ring (Figure 2E, Supplementary Figure 11C), suggesting these variants may be nonfunctional forms of the classical snRNAs. In contrast, mouse snRNAs predicted to lack an Sm-site exhibit similar affinity for Sm-proteins as canonical Sm-site containing snRNAs in our standard anti-Sm-RIPs (Figure 2E, bottom), but do not exhibit this same affinity in response to supplementing reactions with ATP (Supplementary Figure 11C, bottom). Provided the annotations for mouse snRNAs are correct, these data suggest that the current definition of a mouse Sm-site may be incomplete. This may not be surprising as mouse snRNAs are very poorly annotated (Supplementary Figure 1, bottom). Also, our program may be unable to correctly identify all mouse snRNAs that receive an Sm-ring as we defined Sm-sites based on the human sequences. Alternatively, it is also possible, though unlikely, that a large fraction of mouse snRNAs may not require ATP to receive an Sm-ring. Overall, our algorithm specifically identifies Sm-sites in the expected snRNAs and can additionally inform the Sm-ring potential of specific variants.

### Evidence for Sm ring assembly on Sm-site containing mRNAs

Our findings that Sm-site containing RNAs in addition to snRNAs are significantly associated with Sm-proteins (Figure 2C, 2F, Supplementary Figure 7) suggests that Sm-ring containing assemblies are much more widespread in the mammalian transcriptomes than previously known. It has been shown that mRNAs and sno/ scaRNAs can associate with Sm-proteins in HeLa cells (11) and in anti-SmB/B’-CLIP experiments (94). For both of these studies, Sm-protein association was suggested to occur through base-pair complementarity between the 5’ end of U snRNAs in already assembled snRNPs and the target mRNA (11). These studies could only assess those RNAs that associate with Sm-proteins in total cell lysates via immunoprecipitation of the endogenous Sm-RNPs. In comparison, our study has two key advantages for assessing whether Sm-rings are, indeed, assembled on RNAs other than U snRNAs. First, we perform Sm-ring assembly and immunoprecipitations from cytoplasmic cell extracts, identifying Sm-protein associations in the environment Sm-ring assembly occurs within the cell. Restricting to the cytoplasm limits capture of RNAs that associate with Sm-proteins through the spliceosome and the pre-mRNA splicing process. Second, akin to U snRNA *in vitro* assembly assays, we supplemented cytoplasmic extracts with exogenous polyA-RNA and ATP to drive assembly of new Sm-protein rings on RNAs that contain putative Sm-sites. These RIPs were performed in stringent conditions that have been shown to contain only snRNAs that have assembled high-salt and heparin-resistant Sm-rings (48). Additionally, several lines of evidence support that the Sm-RNA interactions we observe on mRNAs, and other RNAs beyond snRNAs, are indeed authentic. We show that association between Sm-proteins and many types of Sm-site containing RNAs are ATP-dependent and Sm-site-dependent, and thus conform to the established rules for Sm-ring assembly on U snRNAs (Figure 4, Supplementary Figure 15-16). We validate this ATP and Sm-site dependence using *in vitro* Sm-ring assembly reactions on NDUFB6, SECISBP2l, Hif1a, and Cul5, mRNA 3’UTRs, providing evidence these mRNAs can indeed accept Sm-rings (Figure 4E, 4G-I, Supplementary Figure 15B, 15D-F). Next, Sm-rings assembled on 4 out of 6 newly identified targets (NDUFB6, SECISBP2l, Hif1a, and Cul5, mRNA 3’UTRs) are stable following a 2 M urea treatment (Figure 4D-J, Supplementary Figure 15), a higher stringency condition that favors an assembled Sm-ring (29,48). Not only are the novel Sm-mRNP interactions we report captured with the Y12 anti-Sm antibody, but they are also captured with an antibody specific to SmB/B’ (Supplementary Figure 16). Lastly, despite differences in the RNA-immunoprecipitation methods and cell types used, the Sm-enriched RNAs from our study show a 23% overlap with Sm targets identified by Lu *et al* 2014 (11) and 40% overlap with those described in Briese *et al* 2019 (94) (Supplementary Figure 6C, 6H). Notably, the majority of these RNAs shared between these studies contain Sm-sites (Supplementary Figure 6B, 6J). Thus, short or long RNAs as well as non-coding or protein-coding RNAs that contain Sm-sites contain the potential to associate with Sm-proteins.

Curiously, a fraction of Sm-site containing RNAs identified in the Sm-RIP experiments, including the KIF5B-3’UTR and Rem2-mRNA, do not exhibit both ATP or Sm-site dependence (Figure 4F, 4J, Supplementary Figure 15C, 15G). Though the KIF5B 3’UTR shows an Sm-site dependent association, stable under the urea conditions, it does not show an ATP-dependent increase for the wild-type RNA (Figure 4F, Supplementary Figure 15C). Among the various reasons for ATP and/or Sm-site independence, one possibility is that some of these sites may resemble recently reported sites that can receive Sm-ring in an ATP independent manner (49). This report provided evidence that U snRNAs predicted to form secondary structures across the Sm-site require ATP and the DEAD-box helicase Gemin3 to be imported into the nucleus, whereas those mutated to reduce Sm-site folding could be imported into the nucleus even when Gemin3 abundance was reduced (49). Additionally, as discussed above, our noncanonical Sm-site definition may be comprised of both *bona fide* Sm-sites, as well as false Sm-sites, potentially explaining why Rem2-mRNA did not show Sm-site or ATP-induced Sm-ring assembly (Figure 4J, Supplementary Figure 15G). It is also possible that *in vitro* transcribed mRNAs used in Sm-ring assembly reactions may be lacking certain base-modifications that may be present in the polyA-RNA libraries used to perform the modified Sm-ring assembly reactions. While it is currently unknown if base-modifications affect Sm-ring assembly, they can influence RNA secondary structure (108,109), which can in turn affect Sm-ring assembly (49). Lastly, the prior proposed mechanisms involving base-pairing between assembled U snRNPs and a target RNA also remains a possibility for ATP and/ or Sm-site independent association between Sm-proteins and RNAs (11). Nonetheless, given the specificity prediction of an Sm-site has on driving enrichment with Sm-proteins, our data suggests the Sm-site is a main contributor to Sm-association for our Sm-site containing mRNAs.

### Impact of Sm-ring assembly on expression of Sm-site containing mRNAs

Although it is largely unknown how SMN-deficiency results in SMA, it is quite clear that reduced SMN in cell or animal models of SMA results in the reduction of U snRNP assembly (48,50,110–114), resulting in degradation of the assembly reactants—U snRNAs and Sm-proteins (9,50,115). Furthermore, correction of the SMA phenotype strongly correlates with improved U snRNP assembly (50,110,113,114,116). In congruence with these studies, we show that RNAs, in particular mRNAs, predicted to contain Sm-sites are globally reduced in abundance in mouse models of SMA as compared to other RNAs (Figure 5A). Additionally, we show that 95% of the downregulated transcripts shared across the three independent datasets contain Sm-sites (Figure 5D). Indeed, canonical Sm-site containing transcripts downregulated in these datasets are enriched in anti-Sm-RIPs (Figure 5E), providing further evidence of a correlation between Sm-site containing mRNAs, Sm-ring assembly, and SMN-deficiency. Therefore, our data supports a proposal that SMA results in reduced capacity of SMN to assemble Sm-rings on mRNAs, resulting in their reduced abundance. Moreover, such a mechanism could contribute along with other proposed mechanisms of SMA pathogenesis including altered pre-mRNA splicing. Thus, the critical targets in SMA could be both mis-spliced and downregulated.

How does the U-rich Sm-site, prominently found in the 3’UTRs of long mRNAs, and the binding of Sm-proteins to these sites, impact expression of these mRNAs? One possibility is that assembly of Sm-rings onto these mRNA interplays with other RNA-binding proteins (RBPs) that bind 3’UTRs to regulate gene expression (92,93). Within the RNA-Binding Protein Database (RBPDB), an experimentally validated collection of RBPs and their binding sites (90), we found that a handful of RBPs are annotated to bind elements that contain the Sm-site sequence. These RBPs include SFRS1, Tia1, ELAVL1/4, gld-1, pos-1, ZFP36, and Cus2. Two of these —SFRS1 and Tia1—are splicing factors that bind U-rich elements in introns to affect pre-mRNA splicing (117,118). Cus2 interacts with the U2 snRNA (119). The rest—ELAVL1/4, gld-1, pos-1, and ZFP36—promote stability of their target mRNAs (117,120–127), some by inhibiting target mRNA translation (122). The most interesting of these are ELAVL1/4, also known as HuR and HuD. HuD is specifically expressed in neurons and it’s overexpression has been shown to partially suppress the SMA phenotype in zebrafish and mouse SMA models, and neurite outgrowth defects in SMN-deficient cells (69,70,76). While the mechanism of SMA suppression by HuD has primarily focused on SMN binding (69,70,76), we suggest that a potential mechanism for HuD-mediated suppression of SMA may occur by stabilizing Sm-site containing mRNAs that do not recieve an Sm-ring as a result of reduced Sm-ring assembly by the SMN complex. Another possibility stems from observations that Sm-ring assembly occurs on immature yeast telomerase RNA, but to then be replaced by the Lsm-ring in mature telomerase (77). It is possible that such a mechanism could be adopted by Sm-site containing mRNAs and other RNAs, whereby Sm proteins may assist or antagonize Lsm-ring assembly within 3’UTRs.

SMN-deficiency has been shown to effect the trafficking of mRNAs in motor neurons. Our findings indicate that Sm-site containing RNAs, and Sm-associated RNAs, are almost exclusively found in motor neuron somas, rejecting the hypothesis that the Sm-site is used to traffic mRNAs to be translated locally at axon terminals (Figure 6A). However, this could raise a new possibility that the Sm-site acts as a signal for retention of the mRNA in the soma. Additionally, our finding that RNAs associated with COPI-mediated intracellular vesicle trafficking are also associated with Sm-proteins, and that the majority of these are predicted to contain Sm-sites (Figure 6C, 6E), suggests yet another mRNA transport mechanism involving 3’UTR Sm-sites. It has previously been shown that overexpression of alpha-COP, the large subunit of the COPI particle, can partially suppress the SMA phenotype in zebrafish, mice, and the neurite outgrowth defects in SMN-deficient neuronal cell cultures (103,104,128). However, like in the case of HuD, the mechanism for this suppression has only focused on alpha-COP association with SMN. Our data suggests a link between alpha-COP mRNA trafficking and Sm-site containing mRNAs. Could the Sm-site confer a signal to incorporate an mRNA into a COPI vesicle for intracellular trafficking, and might the partial suppression of the SMA phenotype by overexpression of alpha-COP occur by increasing the trafficking of these mRNAs?

In summary, we have dramatically expanded the list of RNAs that contain classical and putative Sm-sites and provided stringent criteria for determining Sm-ring assembly potential for these RNAs. Our experiments suggest a novel role for Sm-ring assembly in the regulation of cytoplasmic RNAs, and in particular mRNAs. Exactly how this regulation is controlled and the magnitude of it’s effect on the physiology of Sm-site containing RNAs is an intriguing question. Furthermore, how this novel mechanism fits into the pathogenesis of SMN-deficiency in SMA and whether it directly ties SMN function to affected RNA processing events will be beneficial to future progress on understanding the disease.

## Supporting information

Table of predicted Sm-sites

Summary table of Sm-sites with frequency per single gene product

Sm-RIP Wald test comparisons used to generate figures

Sequences for mRNA candidates used to test Sm-ring assembly

Plasmid maps for all in vitro transcribed candidate RNAs

## DATA AVAILABILITY

Data supporting findings for this study are available in this manuscript and associated Supplementary Data and Files. Raw sequencing data are uploaded to NCBI GEO278538. Custom scripts and master tables used for generating figures are available in Github <https://github.com/ajblatnik/sm_ring_assembly_mrna.git>.

## SUPPLEMENTARY DATA

Supplementary Data and Files are available at NAR online.

## AUTHOR CONTRIBUTIONS

AHMB, AJBIII, and GS jointly conceived the project with design input from MS and WT. Informatic pipeline for identifying Sm-sites was created by AJBIII, BP, and JS. Biochemical and sequencing experiments were performed and analyzed by AJBIII with conceptual and design input from AHMB, BP, CE, GS, MS, and WT. Sequences for *in vitro* transcription plasmids were designed by AJBIII and cloning was performed by Invitrogen GeneArt Gene Synthesis. Informatic analysis of published datasets was performed by MS with conceptual and design input from AJBIII. The manuscript was written by AJBIII and edited by AHMB, GS, MS, and WT. All authors reviewed and approved final submission.

## ACKNOWLEDGEMENTS

We thank Dr. Jill Weimer and Sandford Research for the S3 fibroblast line. Gratitude is extended to Dr. Kathrin Meyer at The Abigail Wexner Research Institute at Nationwide Children’s Hospital for technical advice in inducing the S3 cells to a neural progenitor fate. We would like to acknowledge the Ohio Super Computer, Vienna RNAfold, The Ohio State University Comprehensive Cancer Center Genomics Core, NovoGene, BioRender (Blatnik, A. (2024) BioRender.com/x41i231), and Invitrogen GeneArt Gene Synthesis for their services.

## REFERENCES

1. Kambach, C., Walke, S., Young, R., Avis, J.M., de la Fortelle, E., Raker, V.A., Luhrmann, R., Li, J. and Nagai, K. (1999) Crystal structures of two Sm protein complexes and their implications for the assembly of the spliceosomal snRNPs. Cell, 96, 375–387.

2. Raker, V.A., Hartmuth, K., Kastner, B. and Luhrmann, R. (1999) Spliceosomal U snRNP core assembly: Sm proteins assemble onto an Sm site RNA nonanucleotide in a specific and thermodynamically stable manner. Mol. Cell. Biol., 19, 6554–6565.

3. Stark, H., Dube, P., Luhrmann, R. and Kastner, B. (2001) Arrangement of RNA and proteins in the spliceosomal U1 small nuclear ribonucleoprotein particle. Nature, 409, 539–542.

4. Urlaub, H., Raker, V.A., Kostka, S. and Luhrmann, R. (2001) Sm protein-Sm site RNA interactions within the inner ring of the spliceosomal snRNP core structure. EMBO J., 20, 187–196.

5. Chari, A., Golas, M.M., Klingenhager, M., Neuenkirchen, N., Sander, B., Englbrecht, C., Sickmann, A., Stark, H. and Fischer, U. (2008) An assembly chaperone collaborates with the SMN complex to generate spliceosomal SnRNPs. Cell, 135, 497–509.

6. Zhang, R., So, B.R., Li, P., Yong, J., Glisovic, T., Wan, L. and Dreyfuss, G. (2011) Structure of a key intermediate of the SMN complex reveals Gemin2’s crucial function in snRNP assembly. Cell, 146, 384–395.

7. Wahl, M.C., Will, C.L. and Luhrmann, R. (2009) The spliceosome: design principles of a dynamic RNP machine. Cell, 136, 701–718.

8. Matera, A.G. and Wang, Z. (2014) A day in the life of the spliceosome. Nat. Rev. Mol. Cell Biol., 15, 108–121.

9. Prusty, A.B., Meduri, R., Prusty, B.K., Vanselow, J., Schlosser, A. and Fischer, U. (2017) Impaired spliceosomal UsnRNP assembly leads to Sm mRNA down-regulation and Sm protein degradation. J. Cell Biol., 216, 2391–2407.

10. Roithova, A., Feketova, Z., Vanacova, S. and Stanek, D. (2020) DIS3L2 and LSm proteins are involved in the surveillance of Sm ring-deficient snRNAs. Nucleic Acids Res., 48, 6184–6197.

11. Lu, Z., Guan, X., Schmidt, C.A. and Matera, A.G. (2014) RIP-seq analysis of eukaryotic Sm proteins identifies three major categories of Sm-containing ribonucleoproteins. Genome Biol, 15, R7.

12. Neuenkirchen, N., Englbrecht, C., Ohmer, J., Ziegenhals, T., Chari, A. and Fischer, U. (2015) Reconstitution of the human U snRNP assembly machinery reveals stepwise Sm protein organization. EMBO J., 34, 1925–1941.

13. Fischer, U., Liu, Q. and Dreyfuss, G. (1997) The SMN-SIP1 complex has an essential role in spliceosomal snRNP biogenesis. Cell, 90, 1023–1029.

14. Friesen, W.J. and Dreyfuss, G. (2000) Specific sequences of the Sm and Sm-like (Lsm) proteins mediate their interaction with the spinal muscular atrophy disease gene product (SMN). J. Biol. Chem., 275, 26370–26375.

15. Paushkin, S., Charroux, B., Abel, L., Perkinson, R.A., Pellizzoni, L. and Dreyfuss, G. (2000) The survival motor neuron protein of Schizosacharomyces pombe. Conservation of survival motor neuron interaction domains in divergent organisms. J. Biol. Chem., 275, 23841–23846.

16. Meister, G., Eggert, C., Buhler, D., Brahms, H., Kambach, C. and Fischer, U. (2001) Methylation of Sm proteins by a complex containing PRMT5 and the putative U snRNP assembly factor pICln. Curr. Biol., 11, 1990–1994.

17. Meister, G., Eggert, C. and Fischer, U. (2002) SMN-mediated assembly of RNPs: a complex story. Trends Cell Biol., 12, 472–478.

18. Paushkin, S., Gubitz, A.K., Massenet, S. and Dreyfuss, G. (2002) The SMN complex, an assemblyosome of ribonucleoproteins. Curr. Opin. Cell Biol., 14, 305–312.

19. Pellizzoni, L., Baccon, J., Rappsilber, J., Mann, M. and Dreyfuss, G. (2002) Purification of native survival of motor neurons complexes and identification of Gemin6 as a novel component. J. Biol. Chem., 277, 7540–7545.

20. Pellizzoni, L., Yong, J. and Dreyfuss, G. (2002) Essential role for the SMN complex in the specificity of snRNP assembly. Science, 298, 1775–1779.

21. Pillai, R.S., Grimmler, M., Meister, G., Will, C.L., Luhrmann, R., Fischer, U. and Schumperli, D. (2003) Unique Sm core structure of U7 snRNPs: assembly by a specialized SMN complex and the role of a new component, Lsm11, in histone RNA processing. Genes Dev., 17, 2321–2333.

22. Golembe, T.J., Yong, J. and Dreyfuss, G. (2005) Specific sequence features, recognized by the SMN complex, identify snRNAs and determine their fate as snRNPs. Mol. Cell. Biol., 25, 10989–11004.

23. Gonsalvez, G.B., Tian, L., Ospina, J.K., Boisvert, F.M., Lamond, A.I. and Matera, A.G. (2007) Two distinct arginine methyltransferases are required for biogenesis of Sm-class ribonucleoproteins. J. Cell Biol., 178, 733–740.

24. Kroiss, M., Schultz, J., Wiesner, J., Chari, A., Sickmann, A. and Fischer, U. (2008) Evolution of an RNP assembly system: a minimal SMN complex facilitates formation of UsnRNPs in Drosophila melanogaster. Proc Natl Acad Sci U S A, 105, 10045–10050.

25. Fischer, U., Englbrecht, C. and Chari, A. (2011) Biogenesis of spliceosomal small nuclear ribonucleoproteins. Wiley Interdiscip Rev RNA, 2, 718–731.

26. Grimm, C., Chari, A., Pelz, J.P., Kuper, J., Kisker, C., Diederichs, K., Stark, H., Schindelin, H. and Fischer, U. (2013) Structural basis of assembly chaperone-mediated snRNP formation. Mol. Cell, 49, 692–703.

27. Yong, J., Golembe, T.J., Battle, D.J., Pellizzoni, L. and Dreyfuss, G. (2004) snRNAs contain specific SMN-binding domains that are essential for snRNP assembly. Mol. Cell. Biol., 24, 2747–2756.

28. Pillai, R.S., Will, C.L., Luhrmann, R., Schumperli, D. and Muller, B. (2001) Purified U7 snRNPs lack the Sm proteins D1 and D2 but contain Lsm10, a new 14 kDa Sm D1-like protein. EMBO J., 20, 5470–5479.

29. Tisdale, S., Lotti, F., Saieva, L., Van Meerbeke, J.P., Crawford, T.O., Sumner, C.J., Mentis, G.Z. and Pellizzoni, L. (2013) SMN is essential for the biogenesis of U7 small nuclear ribonucleoprotein and 3’-end formation of histone mRNAs. Cell Rep, 5, 1187–1195.

30. Raker, V.A., Plessel, G. and Luhrmann, R. (1996) The snRNP core assembly pathway: identification of stable core protein heteromeric complexes and an snRNP subcore particle in vitro. EMBO J., 15, 2256–2269.

31. Weber, G., Trowitzsch, S., Kastner, B., Luhrmann, R. and Wahl, M.C. (2010) Functional organization of the Sm core in the crystal structure of human U1 snRNP. EMBO J., 29, 4172–4184.

32. Mattaj, I.W. (1986) Cap trimethylation of U snRNA is cytoplasmic and dependent on U snRNP protein binding. Cell, 46, 905–911.

33. Fischer, U. and Luhrmann, R. (1990) An essential signaling role for the m3G cap in the transport of U1 snRNP to the nucleus. Science, 249, 786–790.

34. Hamm, J., Darzynkiewicz, E., Tahara, S.M. and Mattaj, I.W. (1990) The trimethylguanosine cap structure of U1 snRNA is a component of a bipartite nuclear targeting signal. Cell, 62, 569–577.

35. Mouaikel, J., Bujnicki, J.M., Tazi, J. and Bordonne, R. (2003) Sequence-structure-function relationships of Tgs1, the yeast snRNA/snoRNA cap hypermethylase. Nucleic Acids Res., 31, 4899–4909.

36. Mouaikel, J., Narayanan, U., Verheggen, C., Matera, A.G., Bertrand, E., Tazi, J. and Bordonne, R. (2003) Interaction between the small-nuclear-RNA cap hypermethylase and the spinal muscular atrophy protein, survival of motor neuron. EMBO Rep, 4, 616–622.

37. Izaurralde, E., Lewis, J., Gamberi, C., Jarmolowski, A., McGuigan, C. and Mattaj, I.W. (1995) A cap-binding protein complex mediating U snRNA export. Nature, 376, 709–712.

38. Palacios, I., Hetzer, M., Adam, S.A. and Mattaj, I.W. (1997) Nuclear import of U snRNPs requires importin beta. EMBO J., 16, 6783–6792.

39. Huber, J., Cronshagen, U., Kadokura, M., Marshallsay, C., Wada, T., Sekine, M. and Luhrmann, R. (1998) Snurportin1, an m3G-cap-specific nuclear import receptor with a novel domain structure. EMBO J., 17, 4114–4126.

40. Massenet, S., Pellizzoni, L., Paushkin, S., Mattaj, I.W. and Dreyfuss, G. (2002) The SMN complex is associated with snRNPs throughout their cytoplasmic assembly pathway. Mol. Cell. Biol., 22, 6533–6541.

41. Narayanan, U., Ospina, J.K., Frey, M.R., Hebert, M.D. and Matera, A.G. (2002) SMN, the spinal muscular atrophy protein, forms a pre-import snRNP complex with snurportin1 and importin beta. Hum. Mol. Genet., 11, 1785–1795.

42. Narayanan, U., Achsel, T., Luhrmann, R. and Matera, A.G. (2004) Coupled in vitro import of U snRNPs and SMN, the spinal muscular atrophy protein. Mol. Cell, 16, 223–234.

43. Mahmoudi, S., Henriksson, S., Weibrecht, I., Smith, S., Soderberg, O., Stromblad, S., Wiman, K.G. and Farnebo, M. (2010) WRAP53 is essential for Cajal body formation and for targeting the survival of motor neuron complex to Cajal bodies. PLoS Biol., 8, e1000521.

44. Jady, B.E., Darzacq, X., Tucker, K.E., Matera, A.G., Bertrand, E. and Kiss, T. (2003) Modification of Sm small nuclear RNAs occurs in the nucleoplasmic Cajal body following import from the cytoplasm. EMBO J., 22, 1878–1888.

45. Richard, P., Darzacq, X., Bertrand, E., Jady, B.E., Verheggen, C. and Kiss, T. (2003) A common sequence motif determines the Cajal body-specific localization of box H/ACA scaRNAs. EMBO J., 22, 4283–4293.

46. Dominski, Z. and Marzluff, W.F. (2007) Formation of the 3’ end of histone mRNA: getting closer to the end. Gene, 396, 373–390.

47. Makarov, E.M., Owen, N., Bottrill, A. and Makarova, O.V. (2012) Functional mammalian spliceosomal complex E contains SMN complex proteins in addition to U1 and U2 snRNPs. Nucleic Acids Res., 40, 2639–2652.

48. Wan, L., Battle, D.J., Yong, J., Gubitz, A.K., Kolb, S.J., Wang, J. and Dreyfuss, G. (2005) The survival of motor neurons protein determines the capacity for snRNP assembly: biochemical deficiency in spinal muscular atrophy. Mol. Cell. Biol., 25, 5543–5551.

49. Panek, J., Roithova, A., Radivojevic, N., Sykora, M., Prusty, A.B., Huston, N., Wan, H., Pyle, A.M., Fischer, U. and Stanek, D. (2023) The SMN complex drives structural changes in human snRNAs to enable snRNP assembly. Nat Commun, 14, 6580.

50. Gabanella, F., Butchbach, M.E., Saieva, L., Carissimi, C., Burghes, A.H. and Pellizzoni, L. (2007) Ribonucleoprotein assembly defects correlate with spinal muscular atrophy severity and preferentially affect a subset of spliceosomal snRNPs. PLoS One, 2, e921.

51. Pellizzoni, L., Charroux, B., Rappsilber, J., Mann, M. and Dreyfuss, G. (2001) A functional interaction between the survival motor neuron complex and RNA polymerase II. J. Cell Biol., 152, 75–85.

52. Zhao, D.Y., Gish, G., Braunschweig, U., Li, Y., Ni, Z., Schmitges, F.W., Zhong, G., Liu, K., Li, W., Moffat, J. et al. (2016) SMN and symmetric arginine dimethylation of RNA polymerase II C-terminal domain control termination. Nature, 529, 48–53.

53. Pellizzoni, L., Kataoka, N., Charroux, B. and Dreyfuss, G. (1998) A novel function for SMN, the spinal muscular atrophy disease gene product, in pre-mRNA splicing. Cell, 95, 615–624.

54. Liu, Q., Fischer, U., Wang, F. and Dreyfuss, G. (1997) The spinal muscular atrophy disease gene product, SMN, and its associated protein SIP1 are in a complex with spliceosomal snRNP proteins. Cell, 90, 1013–1021.

55. Gubitz, A.K., Feng, W. and Dreyfuss, G. (2004) The SMN complex. Exp. Cell Res., 296, 51–56.

56. Battle, D.J., Lau, C.K., Wan, L., Deng, H., Lotti, F. and Dreyfuss, G. (2006) The Gemin5 protein of the SMN complex identifies snRNAs. Mol. Cell, 23, 273–279.

57. Kolb, S.J., Battle, D.J. and Dreyfuss, G. (2007) Molecular functions of the SMN complex. J Child Neurol, 22, 990–994.

58. Li, D.K., Tisdale, S., Lotti, F. and Pellizzoni, L. (2014) SMN control of RNP assembly: from post-transcriptional gene regulation to motor neuron disease. Semin. Cell Dev. Biol., 32, 22–29.

59. Jones, K.W., Gorzynski, K., Hales, C.M., Fischer, U., Badbanchi, F., Terns, R.M. and Terns, M.P. (2001) Direct interaction of the spinal muscular atrophy disease protein SMN with the small nucleolar RNA-associated protein fibrillarin. J. Biol. Chem., 276, 38645–38651.

60. Pellizzoni, L., Baccon, J., Charroux, B. and Dreyfuss, G. (2001) The survival of motor neurons (SMN) protein interacts with the snoRNP proteins fibrillarin and GAR1. Curr. Biol., 11, 1079–1088.

61. Poole, A.R. and Hebert, M.D. (2016) SMN and coilin negatively regulate dyskerin association with telomerase RNA. Biol Open, 5, 726–735.

62. Sanchez, G., Dury, A.Y., Murray, L.M., Biondi, O., Tadesse, H., El Fatimy, R., Kothary, R., Charbonnier, F., Khandjian, E.W. and Cote, J. (2013) A novel function for the survival motoneuron protein as a translational regulator. Hum. Mol. Genet., 22, 668–684.

63. Sanchez, G., Bondy-Chorney, E., Laframboise, J., Paris, G., Didillon, A., Jasmin, B.J. and Cote, J. (2016) A novel role for CARM1 in promoting nonsense-mediated mRNA decay: potential implications for spinal muscular atrophy. Nucleic Acids Res., 44, 2661–2676.

64. Bernabo, P., Tebaldi, T., Groen, E.J.N., Lane, F.M., Perenthaler, E., Mattedi, F., Newbery, H.J., Zhou, H., Zuccotti, P., Potrich, V. et al. (2017) In Vivo Translatome Profiling in Spinal Muscular Atrophy Reveals a Role for SMN Protein in Ribosome Biology. Cell Rep, 21, 953–965.

65. Lauria, F., Bernabo, P., Tebaldi, T., Groen, E.J.N., Perenthaler, E., Maniscalco, F., Rossi, A., Donzel, D., Clamer, M., Marchioretto, M. et al. (2020) SMN-primed ribosomes modulate the translation of transcripts related to spinal muscular atrophy. Nat. Cell Biol., 22, 1239–1251.

66. Piazzon, N., Schlotter, F., Lefebvre, S., Dodre, M., Mereau, A., Soret, J., Besse, A., Barkats, M., Bordonne, R., Branlant, C. et al. (2013) Implication of the SMN complex in the biogenesis and steady state level of the signal recognition particle. Nucleic Acids Res., 41, 1255–1272.

67. Tadesse, H., Deschenes-Furry, J., Boisvenue, S. and Cote, J. (2008) KH-type splicing regulatory protein interacts with survival motor neuron protein and is misregulated in spinal muscular atrophy. Hum. Mol. Genet., 17, 506–524.

68. Akten, B., Kye, M.J., Hao le, T., Wertz, M.H., Singh, S., Nie, D., Huang, J., Merianda, T.T., Twiss, J.L., Beattie, C.E. et al. (2011) Interaction of survival of motor neuron (SMN) and HuD proteins with mRNA cpg15 rescues motor neuron axonal deficits. Proc Natl Acad Sci U S A, 108, 10337–10342.

69. Fallini, C., Zhang, H., Su, Y., Silani, V., Singer, R.H., Rossoll, W. and Bassell, G.J. (2011) The survival of motor neuron (SMN) protein interacts with the mRNA-binding protein HuD and regulates localization of poly(A) mRNA in primary motor neuron axons. J. Neurosci., 31, 3914–3925.

70. Hubers, L., Valderrama-Carvajal, H., Laframboise, J., Timbers, J., Sanchez, G. and Cote, J. (2011) HuD interacts with survival motor neuron protein and can rescue spinal muscular atrophy-like neuronal defects. Hum. Mol. Genet., 20, 553–579.

71. Rage, F., Boulisfane, N., Rihan, K., Neel, H., Gostan, T., Bertrand, E., Bordonne, R. and Soret, J. (2013) Genome-wide identification of mRNAs associated with the protein SMN whose depletion decreases their axonal localization. RNA, 19, 1755–1766.

72. Todd, A.G., Lin, H., Ebert, A.D., Liu, Y. and Androphy, E.J. (2013) COPI transport complexes bind to specific RNAs in neuronal cells. Hum. Mol. Genet., 22, 729–736.

73. Dombert, B., Sivadasan, R., Simon, C.M., Jablonka, S. and Sendtner, M. (2014) Presynaptic localization of Smn and hnRNP R in axon terminals of embryonic and postnatal mouse motoneurons. PLoS One, 9, e110846.

74. Fallini, C., Rouanet, J.P., Donlin-Asp, P.G., Guo, P., Zhang, H., Singer, R.H., Rossoll, W. and Bassell, G.J. (2014) Dynamics of survival of motor neuron (SMN) protein interaction with the mRNA-binding protein IMP1 facilitates its trafficking into motor neuron axons. Dev Neurobiol, 74, 319–332.

75. Fallini, C., Donlin-Asp, P.G., Rouanet, J.P., Bassell, G.J. and Rossoll, W. (2016) Deficiency of the Survival of Motor Neuron Protein Impairs mRNA Localization and Local Translation in the Growth Cone of Motor Neurons. J. Neurosci., 36, 3811–3820.

76. Hao le, T., Duy, P.Q., An, M., Talbot, J., Iyer, C.C., Wolman, M. and Beattie, C.E. (2017) HuD and the Survival Motor Neuron Protein Interact in Motoneurons and Are Essential for Motoneuron Development, Function, and mRNA Regulation. J. Neurosci., 37, 11559–11571.

77. Tang, W., Kannan, R., Blanchette, M. and Baumann, P. (2012) Telomerase RNA biogenesis involves sequential binding by Sm and Lsm complexes. Nature, 484, 260–264.

78. Lorenz, R., Bernhart, S.H., Honer Zu Siederdissen, C., Tafer, H., Flamm, C., Stadler, P.F. and Hofacker, I.L. (2011) ViennaRNA Package 2.0. Algorithms Mol Biol, 6, 26.

79. Smedley, D., Haider, S., Ballester, B., Holland, R., London, D., Thorisson, G. and Kasprzyk, A. (2009) BioMart--biological queries made easy. BMC Genomics, 10, 22.

80. Morales, J., Pujar, S., Loveland, J.E., Astashyn, A., Bennett, R., Berry, A., Cox, E., Davidson, C., Ermolaeva, O., Farrell, C.M. et al. (2022) A joint NCBI and EMBL-EBI transcript set for clinical genomics and research. Nature, 604, 310–315.

81. Meyer, K., Ferraiuolo, L., Miranda, C.J., Likhite, S., McElroy, S., Renusch, S., Ditsworth, D., Lagier-Tourenne, C., Smith, R.A., Ravits, J. et al. (2014) Direct conversion of patient fibroblasts demonstrates non-cell autonomous toxicity of astrocytes to motor neurons in familial and sporadic ALS. Proc Natl Acad Sci U S A, 111, 829–832.

82. Blatnik, A.J., McGovern, V.L., Le, T.T., Iyer, C.C., Kaspar, B.K. and Burghes, A.H.M. (2020) Conditional deletion of SMN in cell culture identifies functional SMN alleles. Hum. Mol. Genet., 29, 3477–3492.

83. Dobin, A., Davis, C.A., Schlesinger, F., Drenkow, J., Zaleski, C., Jha, S., Batut, P., Chaisson, M. and Gingeras, T.R. (2013) STAR: ultrafast universal RNA-seq aligner. Bioinformatics, 29, 15–21.

84. Ewels, P., Magnusson, M., Lundin, S. and Kaller, M. (2016) MultiQC: summarize analysis results for multiple tools and samples in a single report. Bioinformatics, 32, 3047–3048.

85. Liao, Y., Smyth, G.K. and Shi, W. (2014) featureCounts: an efficient general purpose program for assigning sequence reads to genomic features. Bioinformatics, 30, 923–930.

86. Love, M.I., Huber, W. and Anders, S. (2014) Moderated estimation of fold change and dispersion for RNA-seq data with DESeq2. Genome Biol, 15, 550.

87. O’Leary, N.A., Wright, M.W., Brister, J.R., Ciufo, S., Haddad, D., McVeigh, R., Rajput, B., Robbertse, B., Smith-White, B., Ako-Adjei, D. et al. (2016) Reference sequence (RefSeq) database at NCBI: current status, taxonomic expansion, and functional annotation. Nucleic Acids Res., 44, D733–745.

88. Frankish, A., Diekhans, M., Ferreira, A.M., Johnson, R., Jungreis, I., Loveland, J., Mudge, J.M., Sisu, C., Wright, J., Armstrong, J. et al. (2019) GENCODE reference annotation for the human and mouse genomes. Nucleic Acids Res., 47, D766–D773.

89. Durinck, S., Spellman, P.T., Birney, E. and Huber, W. (2009) Mapping identifiers for the integration of genomic datasets with the R/Bioconductor package biomaRt. Nat Protoc, 4, 1184–1191.

90. Cook, K.B., Kazan, H., Zuberi, K., Morris, Q. and Hughes, T.R. (2011) RBPDB: a database of RNA-binding specificities. Nucleic Acids Res., 39, D301–308.

91. He, H., Liyanarachchi, S., Akagi, K., Nagy, R., Li, J., Dietrich, R.C., Li, W., Sebastian, N., Wen, B., Xin, B. et al. (2011) Mutations in U4atac snRNA, a component of the minor spliceosome, in the developmental disorder MOPD I. Science, 332, 238–240.

92. Leppek, K., Das, R. and Barna, M. (2018) Functional 5’ UTR mRNA structures in eukaryotic translation regulation and how to find them. Nat. Rev. Mol. Cell Biol., 19, 158–174.

93. Mayr, C. (2019) What Are 3’ UTRs Doing? Cold Spring Harb Perspect Biol, 11.

94. Briese, M., Haberman, N., Sibley, C.R., Faraway, R., Elser, A.S., Chakrabarti, A.M., Wang, Z., Konig, J., Perera, D., Wickramasinghe, V.O. et al. (2019) A systems view of spliceosomal assembly and branchpoints with iCLIP. Nat. Struct. Mol. Biol., 26, 930–940.

95. Zhang, Z., Pinto, A.M., Wan, L., Wang, W., Berg, M.G., Oliva, I., Singh, L.N., Dengler, C., Wei, Z. and Dreyfuss, G. (2013) Dysregulation of synaptogenesis genes antecedes motor neuron pathology in spinal muscular atrophy. Proc Natl Acad Sci U S A, 110, 19348–19353.

96. Maeda, M., Harris, A.W., Kingham, B.F., Lumpkin, C.J., Opdenaker, L.M., McCahan, S.M., Wang, W. and Butchbach, M.E. (2014) Transcriptome profiling of spinal muscular atrophy motor neurons derived from mouse embryonic stem cells. PLoS One, 9, e106818.

97. Doktor, T.K., Hua, Y., Andersen, H.S., Broner, S., Liu, Y.H., Wieckowska, A., Dembic, M., Bruun, G.H., Krainer, A.R. and Andresen, B.S. (2017) RNA-sequencing of a mouse-model of spinal muscular atrophy reveals tissue-wide changes in splicing of U12-dependent introns. Nucleic Acids Res., 45, 395–416.

98. Nichterwitz, S., Nijssen, J., Storvall, H., Schweingruber, C., Comley, L.H., Allodi, I., Lee, M.V., Deng, Q., Sandberg, R. and Hedlund, E. (2020) LCM-seq reveals unique transcriptional adaptation mechanisms of resistant neurons and identifies protective pathways in spinal muscular atrophy. Genome Res., 30, 1083–1096.

99. Hsieh-Li, H.M., Chang, J.G., Jong, Y.J., Wu, M.H., Wang, N.M., Tsai, C.H. and Li, H. (2000) A mouse model for spinal muscular atrophy. Nat. Genet., 24, 66–70.

100. Monani, U.R., Sendtner, M., Coovert, D.D., Parsons, D.W., Andreassi, C., Le, T.T., Jablonka, S., Schrank, B., Rossoll, W., Prior, T.W. et al. (2000) The human centromeric survival motor neuron gene (SMN2) rescues embryonic lethality in Smn(-/-) mice and results in a mouse with spinal muscular atrophy. Hum. Mol. Genet., 9, 333–339.

101. Le, T.T., Pham, L.T., Butchbach, M.E., Zhang, H.L., Monani, U.R., Coovert, D.D., Gavrilina, T.O., Xing, L., Bassell, G.J. and Burghes, A.H. (2005) SMNΔ7, the major product of the centromeric survival motor neuron (SMN2) gene, extends survival in mice with spinal muscular atrophy and associates with full-length SMN. Hum. Mol. Genet., 14, 845–857.

102. Nijssen, J., Aguila, J., Hoogstraaten, R., Kee, N. and Hedlund, E. (2018) Axon-Seq Decodes the Motor Axon Transcriptome and Its Modulation in Response to ALS. Stem Cell Reports, 11, 1565–1578.

103. Li, H., Custer, S.K., Gilson, T., Hao le, T., Beattie, C.E. and Androphy, E.J. (2015) alpha-COP binding to the survival motor neuron protein SMN is required for neuronal process outgrowth. Hum. Mol. Genet., 24, 7295–7307.

104. Custer, S.K., Astroski, J.W., Li, H.X. and Androphy, E.J. (2019) Interaction between alpha-COP and SMN ameliorates disease phenotype in a mouse model of spinal muscular atrophy. Biochem. Biophys. Res. Commun., 514, 530–537.

105. Pannone, B.K., Xue, D. and Wolin, S.L. (1998) A role for the yeast La protein in U6 snRNP assembly: evidence that the La protein is a molecular chaperone for RNA polymerase III transcripts. EMBO J., 17, 7442–7453.

106. Achsel, T., Brahms, H., Kastner, B., Bachi, A., Wilm, M. and Luhrmann, R. (1999) A doughnut-shaped heteromer of human Sm-like proteins binds to the 3’-end of U6 snRNA, thereby facilitating U4/U6 duplex formation in vitro. EMBO J., 18, 5789–5802.

107. Pannone, B.K., Kim, S.D., Noe, D.A. and Wolin, S.L. (2001) Multiple functional interactions between components of the Lsm2-Lsm8 complex, U6 snRNA, and the yeast La protein. Genetics, 158, 187–196.

108. Roundtree, I.A., Evans, M.E., Pan, T. and He, C. (2017) Dynamic RNA Modifications in Gene Expression Regulation. Cell, 169, 1187–1200.

109. Boo, S.H. and Kim, Y.K. (2020) The emerging role of RNA modifications in the regulation of mRNA stability. Exp Mol Med, 52, 400–408.

110. Workman, E., Saieva, L., Carrel, T.L., Crawford, T.O., Liu, D., Lutz, C., Beattie, C.E., Pellizzoni, L. and Burghes, A.H. (2009) A SMN missense mutation complements SMN2 restoring snRNPs and rescuing SMA mice. Hum. Mol. Genet., 18, 2215–2229.

111. Lotti, F., Imlach, W.L., Saieva, L., Beck, E.S., Hao le, T., Li, D.K., Jiao, W., Mentis, G.Z., Beattie, C.E., McCabe, B.D. et al. (2012) An SMN-dependent U12 splicing event essential for motor circuit function. Cell, 151, 440–454.

112. Li, D.K., Tisdale, S., Espinoza-Derout, J., Saieva, L., Lotti, F. and Pellizzoni, L. (2013) A cell system for phenotypic screening of modifiers of SMN2 gene expression and function. PLoS One, 8, e71965.

113. Iyer, C.C., Corlett, K.M., Massoni-Laporte, A., Duque, S.I., Madabusi, N., Tisdale, S., McGovern, V.L., Le, T.T., Zaworski, P.G., Arnold, W.D. et al. (2018) Mild SMN missense alleles are only functional in the presence of SMN2 in mammals. Hum. Mol. Genet., 27, 3404–3416.

114. McGovern, V.L., Kray, K.M., Arnold, W.D., Duque, S.I., Iyer, C.C., Massoni-Laporte, A., Workman, E., Patel, A., Battle, D.J. and Burghes, A.H.M. (2020) Intragenic complementation of amino and carboxy terminal SMN missense mutations can rescue Smn null mice. Hum. Mol. Genet., 29, 3493–3503.

115. Zhang, Z., Lotti, F., Dittmar, K., Younis, I., Wan, L., Kasim, M. and Dreyfuss, G. (2008) SMN deficiency causes tissue-specific perturbations in the repertoire of snRNAs and widespread defects in splicing. Cell, 133, 585–600.

116. Winkler, C., Eggert, C., Gradl, D., Meister, G., Giegerich, M., Wedlich, D., Laggerbauer, B. and Fischer, U. (2005) Reduced U snRNP assembly causes motor axon degeneration in an animal model for spinal muscular atrophy. Genes Dev., 19, 2320–2330.

117. Suswam, E.A., Nabors, L.B., Huang, Y., Yang, X. and King, P.H. (2005) IL-1beta induces stabilization of IL-8 mRNA in malignant breast cancer cells via the 3’ untranslated region: Involvement of divergent RNA-binding factors HuR, KSRP and TIAR. Int. J. Cancer, 113, 911–919.

118. Choi, E.Y. and Pintel, D. (2009) Splicing of the large intron present in the nonstructural gene of minute virus of mice is governed by TIA-1/TIAR binding downstream of the nonconsensus donor. J. Virol., 83, 6306–6311.

119. Yan, D., Perriman, R., Igel, H., Howe, K.J., Neville, M. and Ares, M., Jr. (1998) CUS2, a yeast homolog of human Tat-SF1, rescues function of misfolded U2 through an unusual RNA recognition motif. Mol. Cell. Biol., 18, 5000–5009.

120. Joseph, B., Orlian, M. and Furneaux, H. (1998) p21(waf1) mRNA contains a conserved element in its 3’-untranslated region that is bound by the Elav-like mRNA-stabilizing proteins. J. Biol. Chem., 273, 20511–20516.

121. Loflin, P. and Lever, J.E. (2001) HuR binds a cyclic nucleotide-dependent, stabilizing domain in the 3’ untranslated region of Na(+)/glucose cotransporter (SGLT1) mRNA. FEBS Lett., 509, 267–271.

122. Lee, M.H. and Schedl, T. (2004) Translation repression by GLD-1 protects its mRNA targets from nonsense-mediated mRNA decay in C. elegans. Genes Dev., 18, 1047–1059.

123. Tchen, C.R., Brook, M., Saklatvala, J. and Clark, A.R. (2004) The stability of tristetraprolin mRNA is regulated by mitogen-activated protein kinase p38 and by tristetraprolin itself. J. Biol. Chem., 279, 32393–32400.

124. Buratti, E., Stuani, C., De Prato, G. and Baralle, F.E. (2007) SR protein-mediated inhibition of CFTR exon 9 inclusion: molecular characterization of the intronic splicing silencer. Nucleic Acids Res., 35, 4359–4368.

125. Farley, B.M., Pagano, J.M. and Ryder, S.P. (2008) RNA target specificity of the embryonic cell fate determinant POS-1. RNA, 14, 2685–2697.

126. Cho, S.J., Zhang, J. and Chen, X. (2010) RNPC1 modulates the RNA-binding activity of, and cooperates with, HuR to regulate p21 mRNA stability. Nucleic Acids Res., 38, 2256–2267.

127. Wang, P.Y., Rao, J.N., Zou, T., Liu, L., Xiao, L., Yu, T.X., Turner, D.J., Gorospe, M. and Wang, J.Y. (2010) Post-transcriptional regulation of MEK-1 by polyamines through the RNA-binding protein HuR modulating intestinal epithelial apoptosis. Biochem. J., 426, 293–306.

128. Custer, S.K., Todd, A.G., Singh, N.N. and Androphy, E.J. (2013) Dilysine motifs in exon 2b of SMN protein mediate binding to the COPI vesicle protein alpha-COP and neurite outgrowth in a cell culture model of spinal muscular atrophy. Hum. Mol. Genet., 22, 4043–4052.

