## Supplemental Data for "Sm-site containing mRNAs can accept Sm-rings and are downregulated in Spinal Muscular Atrophy"

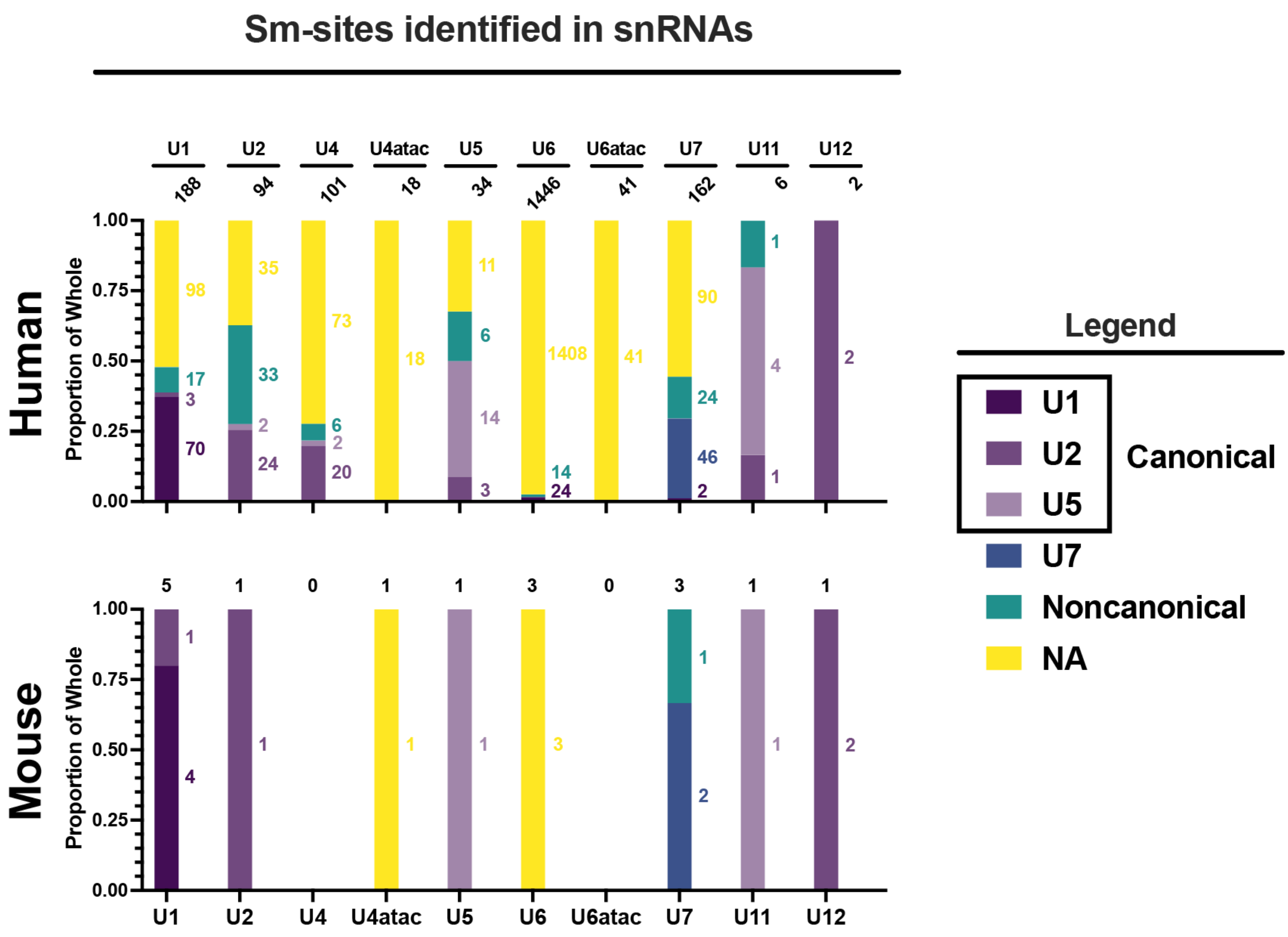


**Supplementary Figure 1: Breakdown of type of Sm-site detected in different annotated U snRNAs. (top)** for human genes, **(bottom)** for mouse genes. For all bar graphs, top numbers are the number of genes represented within the bar, side numbers are the values contributing to the bar for a given Sm-site, in color. Each bar corresponds to the annotation of snRNA—U1, U2, U4, U4atac, U5, U6, U6atac, U7, U11, and U12. Whithin each bare is a breakdown of each Sm-site type—U1, U2, U5, U7, Noncanonical or Absent. Of note, mouse snRNAs are very poorly annotated, with only a few variants, if any, defined for each type.

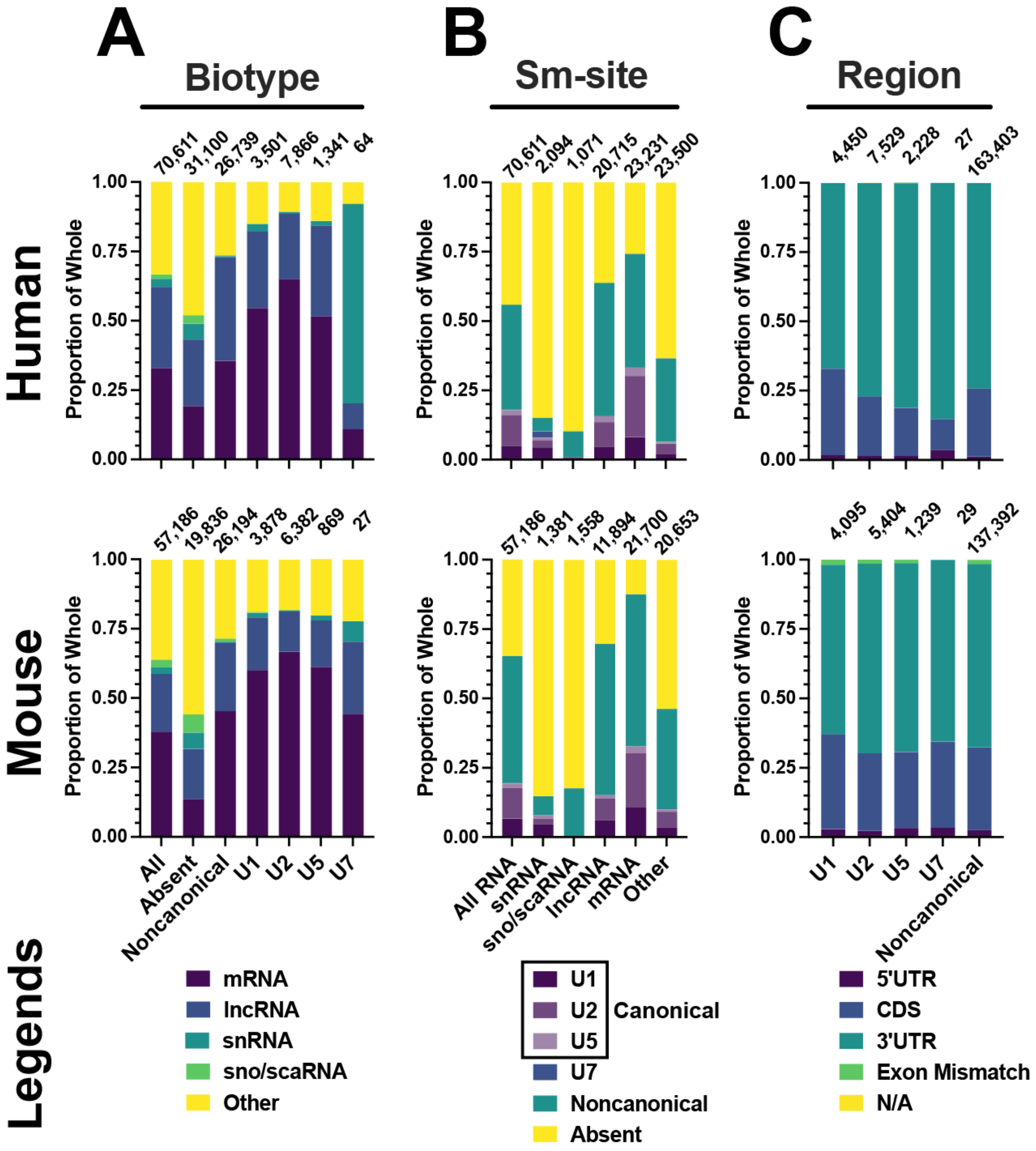


**Supplementary Figure 2: Breakdown of type of Sm-site detected in NCBI Refseq and Gencode human and mouse transcriptomes. (top)** for human genes, **(bottom)** for mouse genes. For all bar graphs, top numbers are the number of genes represented within the bar. Data presented in **A-C** correspond to a single, unique transcript ID of a single, unique gene. **(A)** Proportional bar graph of RNA biotypes. **All** is a breakdown of the annotated genome. **Absent** are RNAs not predicted to contain an Sm-site. **Noncanonical** are those RNAs only predicted to contain noncanonical Sm-site sequences. **U1**, **U2**, **U5**, and **U7** are RNAs predicted to have a U1, U2, U5, or U7 Sm-site, but may have additional Sm-site sequences. **(B)** Proportional bar graph giving a breakdown of types of Sm-sites predicted in each of the following biotypes: All, snRNA, sno/scaRNA, lncRNA, mRNA, and Other. **(C)** Proportional bar graph depicting the region of an mRNA where Sm-sites are predicted.

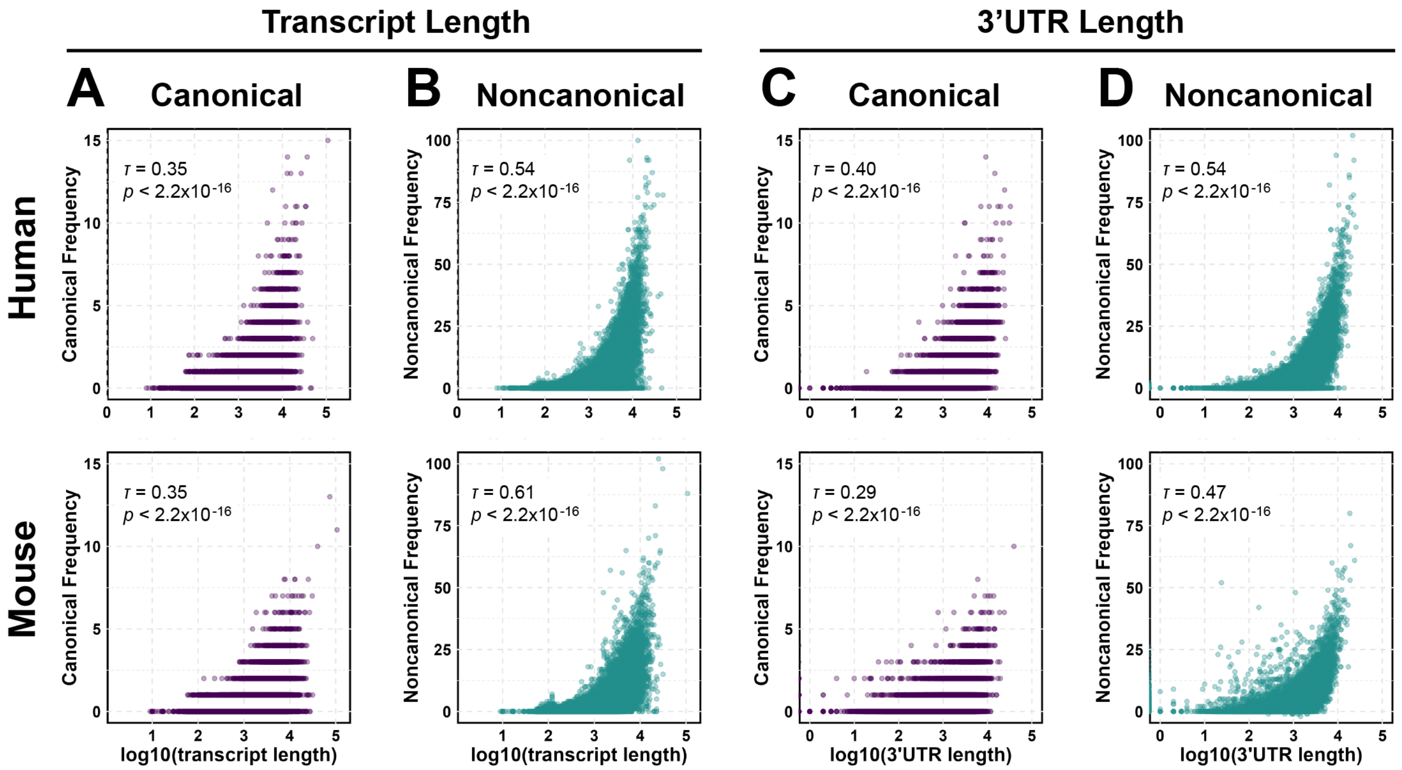


**Supplementary Figure 3. Frequency of Sm-sites is mildly correlated with transcript or 3’UTR length. Top:** human genes, **Bottom:** mouse genes. For all bar graphs, top numbers are the number of genes represented within the bar. Data presented in **A-D** represent a unique transcript ID of a single, unique gene ID. **(A)** Scatter plot of the frequency of canonical Sm-sites predicted in a transcript vs the length of the transcript. *R* is the coefficient of a Kendall rank-ordered test. **(B)** Scatter plot of the frequency of noncanonical Sm-sites predicted in a transcript vs the length of the transcript. $\tau$ is the coefficient of a Kendall rank-ordered test. **(C)** Scatter plot of the frequency of canonical Sm-sites predicted in the 3’UTR of an mRNA vs the length of the mRNA 3’UTR. *R* is the coefficient of a Kendall rank-ordered test. **(D)** Scatter plot of the frequency of noncanonical Sm-sites predicted in the 3’UTR of an mRNA vs the length of the mRNA 3’UTR. $\tau$ is the coefficient of a Kendall rank-ordered test.

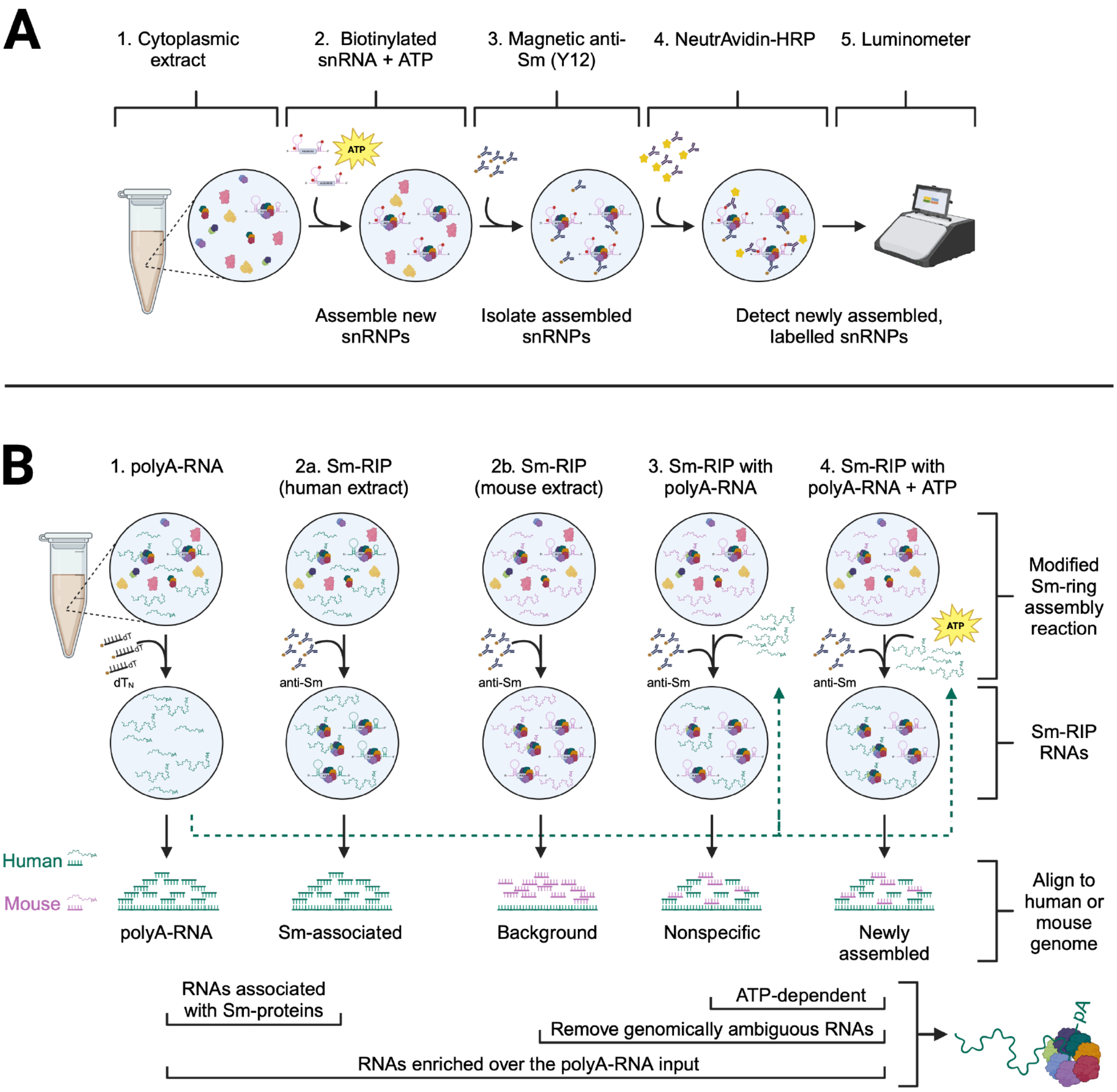


**Supplementary Figure 4: Modification of the standard Sm-ring assembly reaction to test Sm-protein ring assembly on polyA-RNAs. (A)** Schematic of the snRNP assembly reaction described by Wan *et al*. 1) a cytoplasmic cell extract is, 2) incubated with an *in vitro* transcribed, biotinylated human U4 snRNA and ATP. Newly assembled snRNPs are 3) enriched following an anti-Sm (Y12) immunoprecipitation and 4) detected using an NeutrAvidin-HRP antibody and 5) quantified using a luminometer. **(B)** Modified Sm-ring assembly reaction in which polyA-RNA is supplied in place of an *in vitro* transcribed snRNA. Green denotes an RNA or genome of human origin and pink denotes an RNA or genome of mouse origin. Four sample conditions were analyzed. 1) a polyA-RNA enriched library used as input for the Sm-ring assembly reactions and aligned to the genome of origin. 2) an anti-Sm (Y12) RNA immunoprecipitation (Sm-RIP) in cytoplasmic cell extract to identify those RNAs that associate with Sm-proteins under physiological conditions when aligned to the genome of origin (2a), and secondly to remove genomically ambiguous RNAs when aligned to the opposing species genome (2b). 3) an Sm-RIP in which polyA-RNA of one species is incubated with the cytoplasmic extract of the opposing species and aligned to the genome of polyA-RNA origin to identify RNAs that newly associate with Sm-proteins, but that are likely nonspecific. 4) an Sm-RIP in which polyA-RNA of one species is incubated with the cytoplasmic extract of the opposing species and ATP, which aligned to the genome of polyA-RNA origin will identify RNAs with newly assembled Sm-rings. Comparisons outlined underneath conditions were used to identify the candidate polyA-RNAs that most likely recieve an Sm-ring. The candidate should be show an ATP-dependent enrichment, it should be enriched over the polyA-RNA input library, and the RNA should associate with Sm-proteins under physiological conditions. Lastly, reads aligning to both human and mouse genomes (genomically ambiguous) can be removed, as these would be considered false-positives.

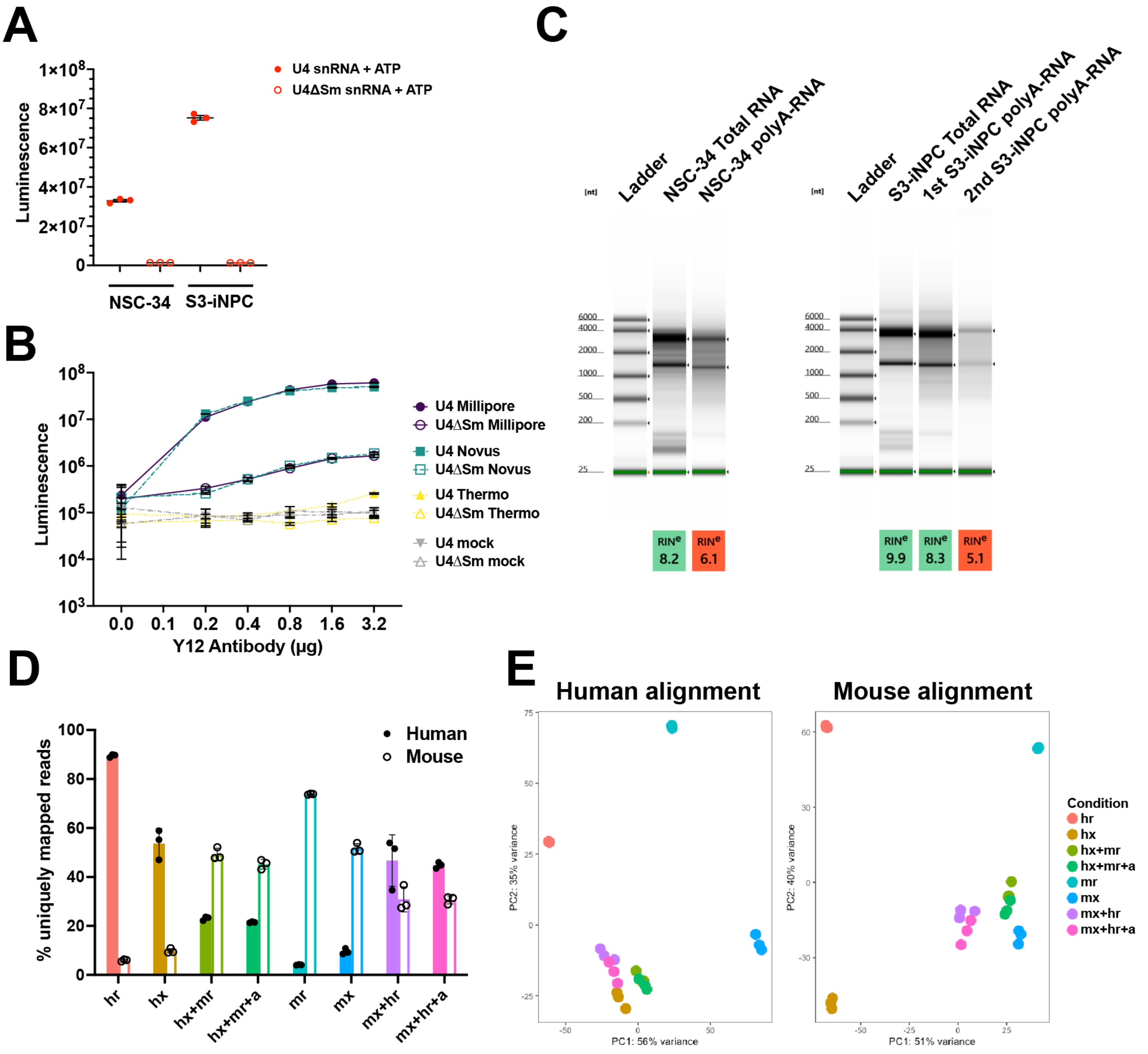


**Supplementary Figure 5: Controls and validation of the experimental outline. (A)** snRNP assembly capacity in cytoplasmic cell extracts used to perform Sm-assembly specific RNA immunoprecipitations. The amount of cytoplasmic cell extract was adjusted proportionally to yield equivalent snRNP assembly capacity. **(B)** snRNP assembly reactions performed to select manufacturer of anti-Sm Y12 antibody. A single lot of the Novus Y12 antibody was used for all subsequent experiments at a ratio of 1.6 µg antibody to 15 µL Dynabeads-Protein G. **(C)** RNA-high sensitivity TapeStation gel image comparing Total RNA to oligo-dT (polyA-RNA) enriched RNA used in Sm-assembly specific RNA immunoprecipitations. Isolation of S3-iNPC polyA-RNA required two rounds of oligo-dT purifiication. Reduced RIN was used to as a proxy for removal of non-polyA-RNAs. **(D)** Percent uniquely mapped, STAR aligned reads for each sequenced libraries. Alignment ot the human GRCh38 genome is depicted by solid shapes and mouse GRCm39 genome by the open shapes. The designation of ‘h’ or ‘m’ identifies the species of the following condition: ‘r’ polyA-RNA, ‘x’ cytoplasmic extract. ‘a’ indicates the addition of ATP. Therefore, mx+hr+a is a mouse cytoplasmic extract incubated with human polyA-RNA and ATP. **(E)** Principal component analysis for each sequencing library color coded by the sample condition. Naming is the same as in (D).

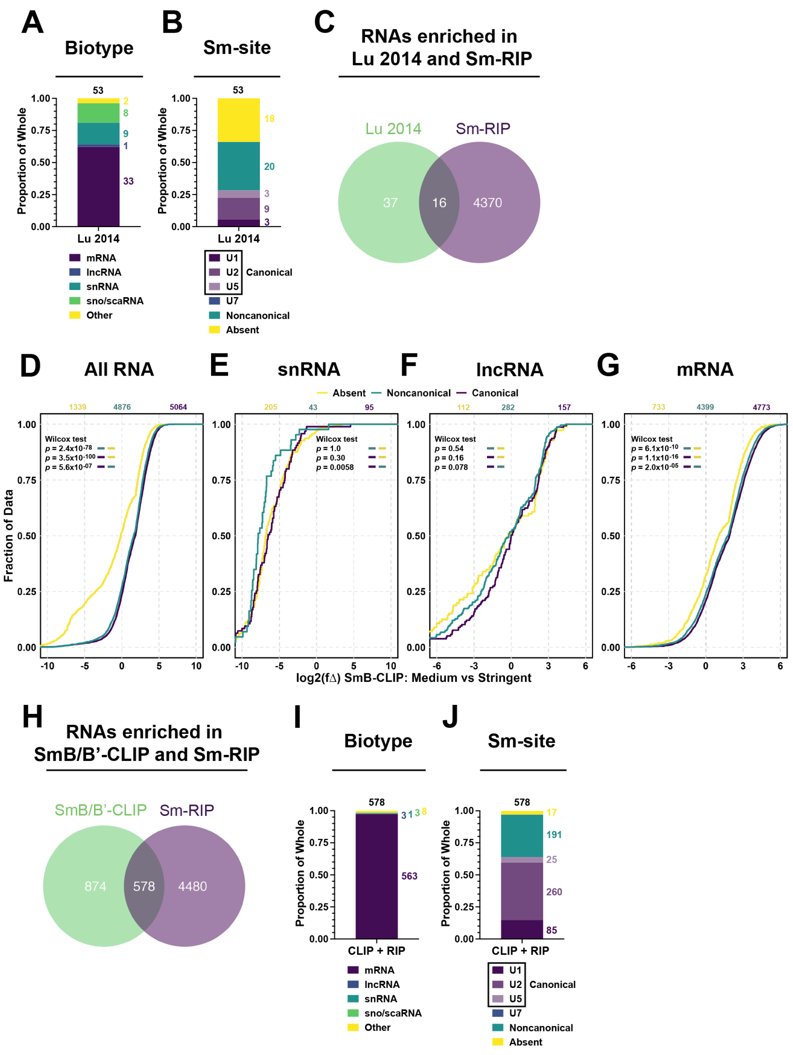


**Supplementary Figure 6: Sm-site containing mRNAs are enriched in previous Sm-RIP and anti-SmB/B'-CLIP-Seq data.** For **(ABIJ)**, values above bars indicate the number of genes contributing to the plots. Numbers in color to the sides of bars indicate the number of genes contributing to the specified group within the bar. **(A)** Proportional bar graph of the RNA biotypes for gene products enriched with Sm-proteins by Lu *et al* Genome Biology 2014. **(B)** Proportional bar graph giving a breakdown of types of Sm-sites predicted in RNAs enriched with Sm-proteins by Lu *et al* Genome Biology 2014. **(C)** Venn diagram depicting the overlapping gene products between different Sm-association conditions. **Lu 2014** are those RNAs found to be enriched with Sm-proteins within the Lu *et al* Genome Biology 2014 study. **Sm-enriched** are RNAs physiologically associated with Sm-proteins—log2(f∆) ≥ 0.6, *padj* < 0.05, between anti-Sm-RIP condition and the polyA-RNA sequencing conditions in Figure 2CD. **(D-G)** Cumulative distribution function plots comparing the Medium and Stringent wash conditions from the anti-SmB-CLIP-Seq data published by Briese *et al* 2020 [Briese Nat Struct Mol Biol 2020]. CDFs included plot **All RNA (D)**, only **snRNA (E)**, only **lncRNA (F)**, or only **mRNA (G)** delineating by Sm-site prediction—**Absent** (yellow), **Noncanonical** (green), **Canonical** (purple). Values above plots are the number of genes plotted for each group in color. *P-values* for Wilcox tests using the alternative that the left color is greater than the right color are provided in the upper-lefthand corner of the plots. **(H)** Venn diagram depicting the overlapping gene products between different Sm-association conditions. **SmB-CLIP** are those RNAs comprised of the highest quartile log2(f∆) between the Medium and Stringent wash conditions as described by Briese *et al* 2020. **Sm-enriched** are RNAs associated with Sm-proteins—log2(f∆) ≥ 0.6, *padj* < 0.05, between anti-Sm-RIP condition and the polyA-RNA sequencing conditions in Figure 2CD. **(I)** Proportional bar graph of the RNA biotypes for gene products shared between the SmB-CLIP and Sm-enriched conditions **(CLIP + RIP)**. **(J)** Proportional bar graph giving a breakdown of types of Sm-sites predicted in RNAs shared between the SmB-CLIP and Sm-enriched conditions **(CLIP + RIP)**.

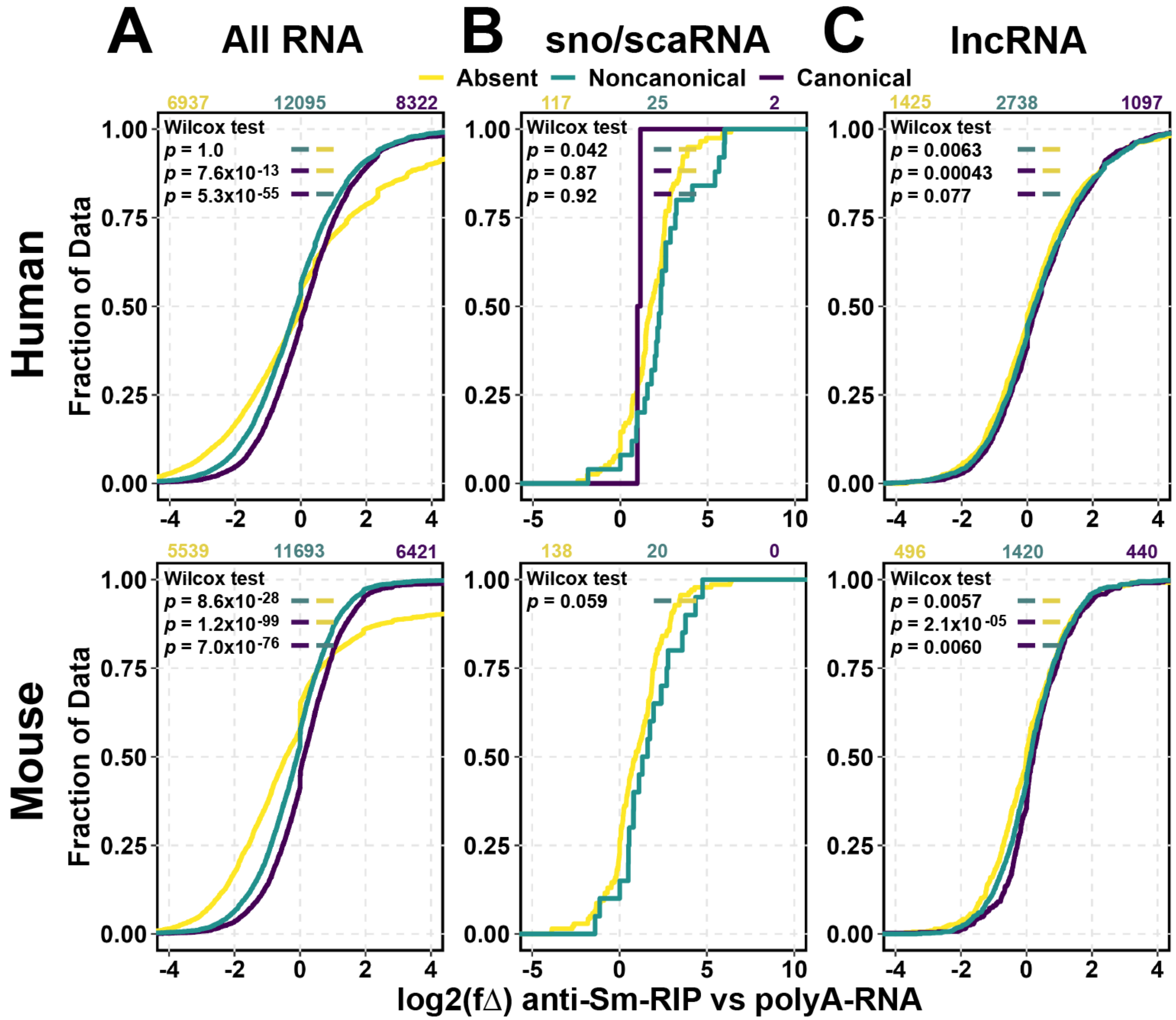


**Supplementary Figure 7. Sm-site containing RNAs are specifically enriched with Sm-proteins.** Cumulative distribution plots for given RNA biotypes, plotting the log2 fold change (log2(fΔ)) by increasing value. log2(fΔ) was calculated between the anti-Sm-RIP vs polyA-RNA conditions. U7 Sm-site-containing RNAs were removed from analsysis. **(A)** All RNA, **(B)** sno/scaRNA, **(C)** lncRNA. Colors designate whether an Sm-site is **Absent** (yellow), **Noncanonical** (green), or **Canonical** (purple). Values above CDFs indicate the number of genes plotted for each condition in color. *P-values* generated from Wilcox tests for the left color being greater than the right color are provided in the upper lefthand corner of the graphs.

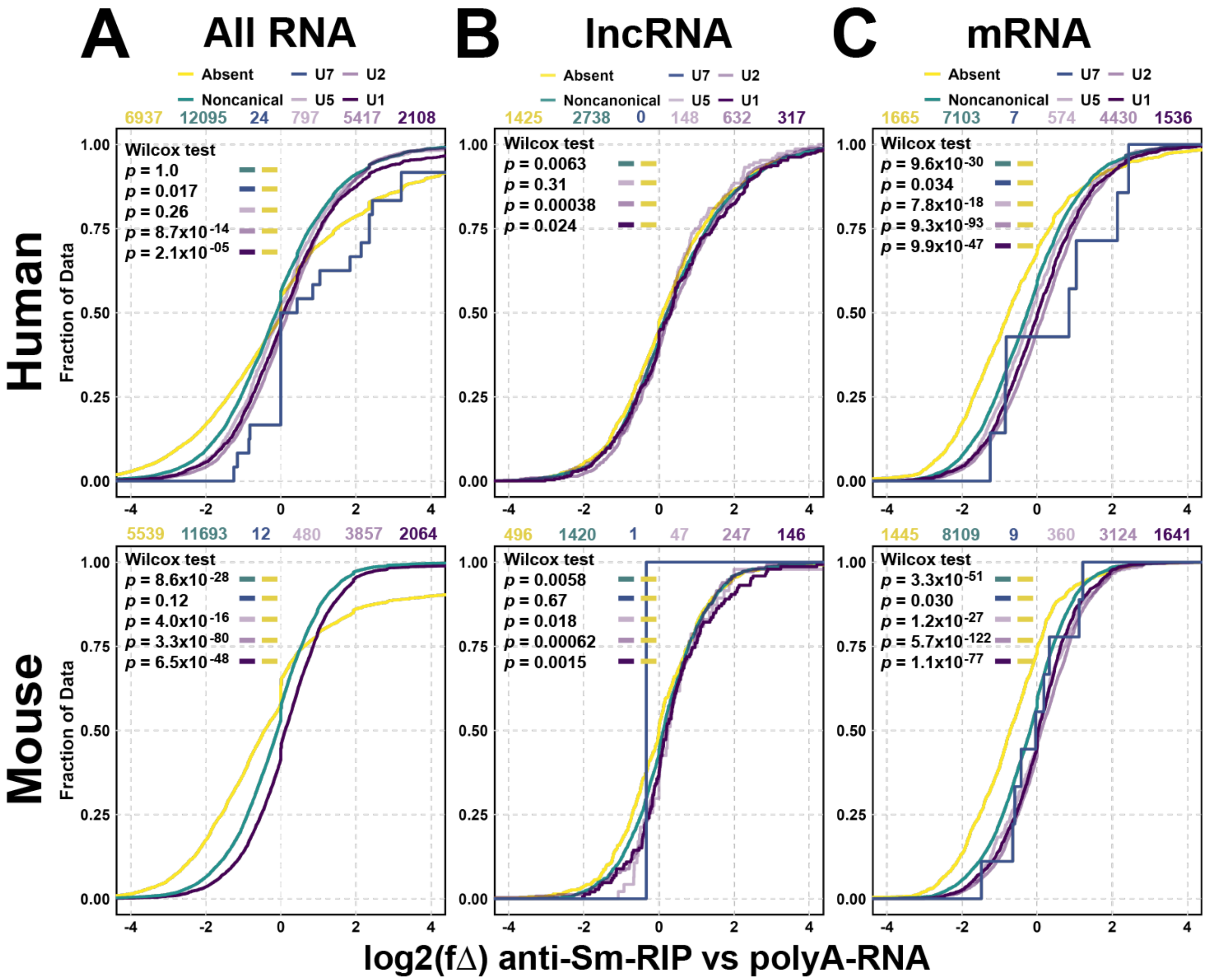


**Supplementary Figure 8: Type of Sm-site correlates with further enrichment with Sm-proteins.** Cumulative distribution plots for given RNA biotypes, plotting the log2 fold change (log2(fΔ)) by increasing value. log2(fΔ) was calculated between the anti-Sm-RIP vs polyA-RNA conditions. (A) All RNA, (B) lncRNA, (C) mRNA. Colors designate whether an Sm-site is Absent (yellow), Noncanonical (green), U7 (blue), or U1, U2, or U5 (purple, darkest to lightest). Values above CDFs indicate the number of genes plotted for each condition in color. *P-values* generated from Wilcox tests for the left color being greater than the right color are provided in the upper lefthand corner of the graphs.

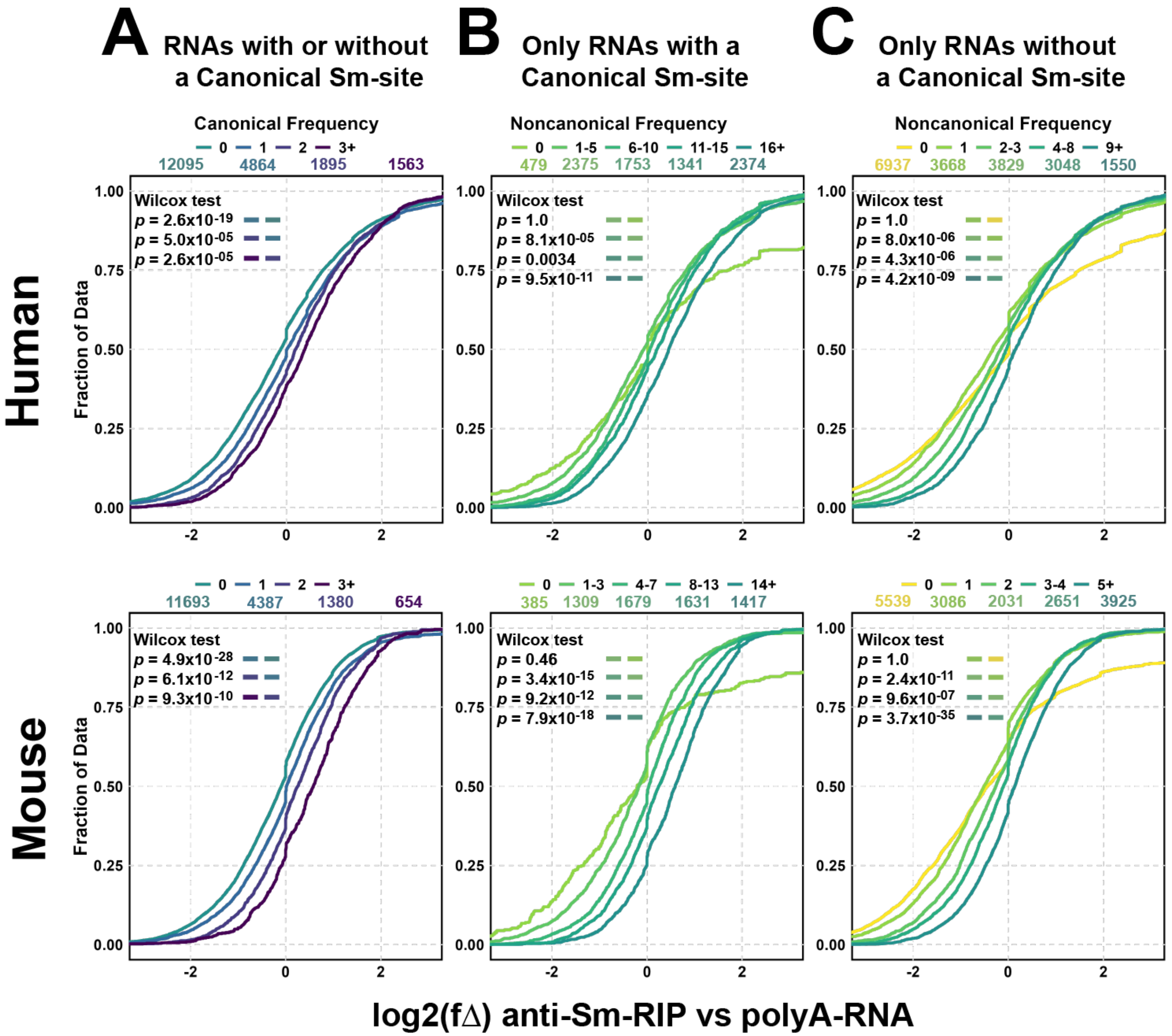


**Supplementary Figure 9: The frequency of Sm-sites in an RNA correlates with Sm-protein enrichment.** **(top)** for human analysis and **(bottom)** for mouse analysis. For each cumulative distribution plot, log2 fold change (log2(fΔ)) between the anti-Sm-RIP and polyA-RNA conditions are plotted in increasing order, delineated by the frequency of Sm-sites predicted for the product of the gene. Values above plots indicate the number of genes plotted for each group designated by color. Wilcox tests were performed using the alternative that the left color is greater than the right color. **(A)** Cumulative distribution plot delineating the frequency of canonical Sm-site prediction in an mRNA transcript by log2 fold change (log2(fΔ)) in anti-Sm-RIP vs the polyA-RNA transcriptome. mRNAs predicted to solely contain noncanonical Sm-sites are not plotted in this graph. **(B)** Cumulative distribution plot of only mRNAs predicted to contain at least one canonical Sm-site, delineating the frequency of noncanonical Sm-sites predicted within the mRNA transcript, against the log2 fold change in anti-Sm-RIP vs the polyA-RNA transcriptome. **(C)** Cumulative distribution plot of mRNAs predicted to solely contain noncanonical Sm-sites, delineating the frequency of noncanonical Sm-sites predicted within the mRNA transcript, against the log2 fold change in anti-Sm-RIP vs the polyA-RNA transcriptome.

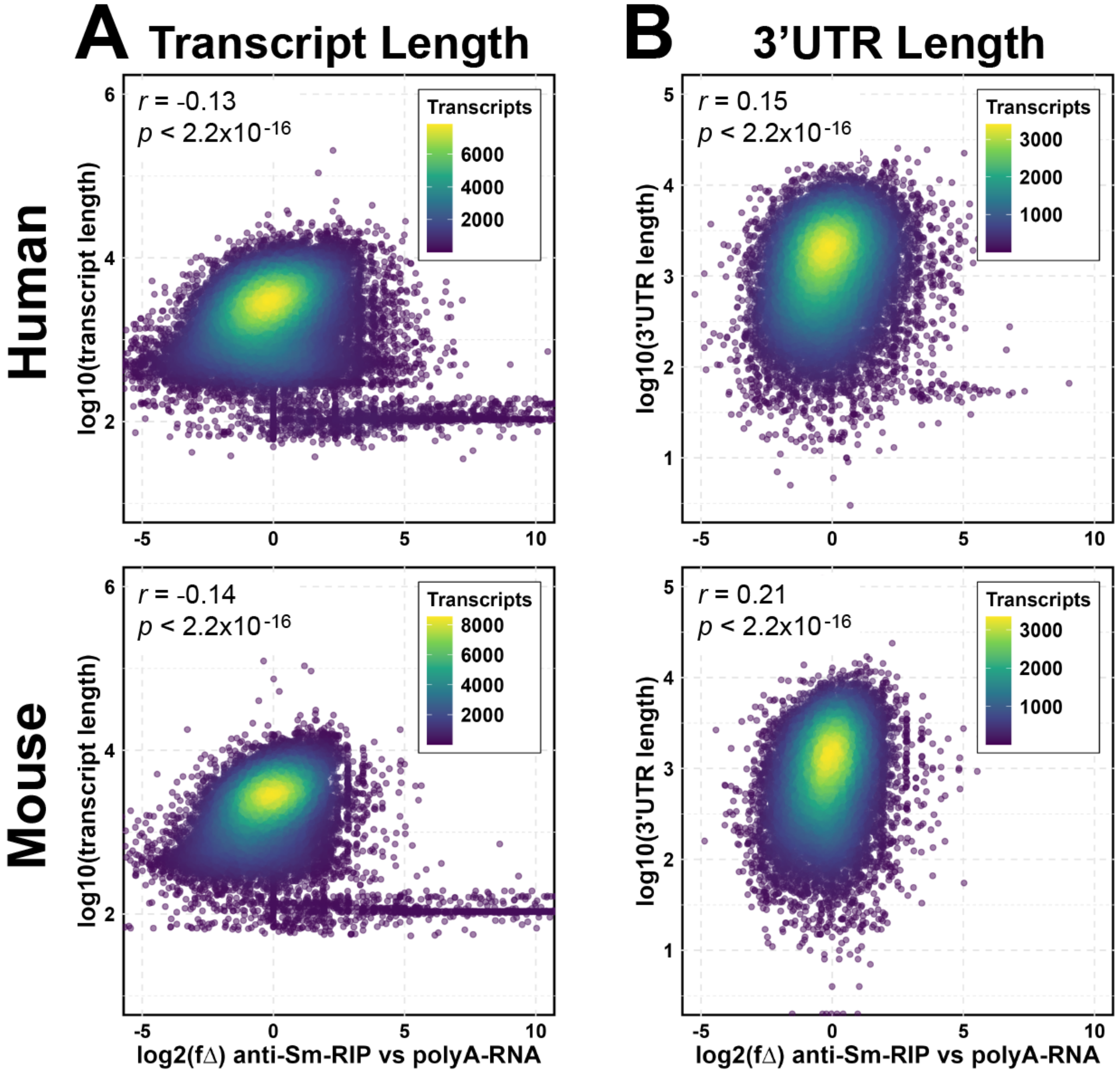


**Supplementary Figure 10: Transcript or 3'UTR length is not a predictor of anti-Sm RIP enrichment. (top)** for human analysis and **(bottom)** for mouse analysis. **(A)** log10 of transcript length plotted against log2 fold change between anti-Sm-RIP and the polyA-RNA transcriptome. *r* is a Pearson correlation coefficient. **(B)** log10 of 3'UTR length plotted against log2 fold change between anti-Sm-RIP and the polyA-RNA transcriptome. *r* is a Pearson correlation coefficient.

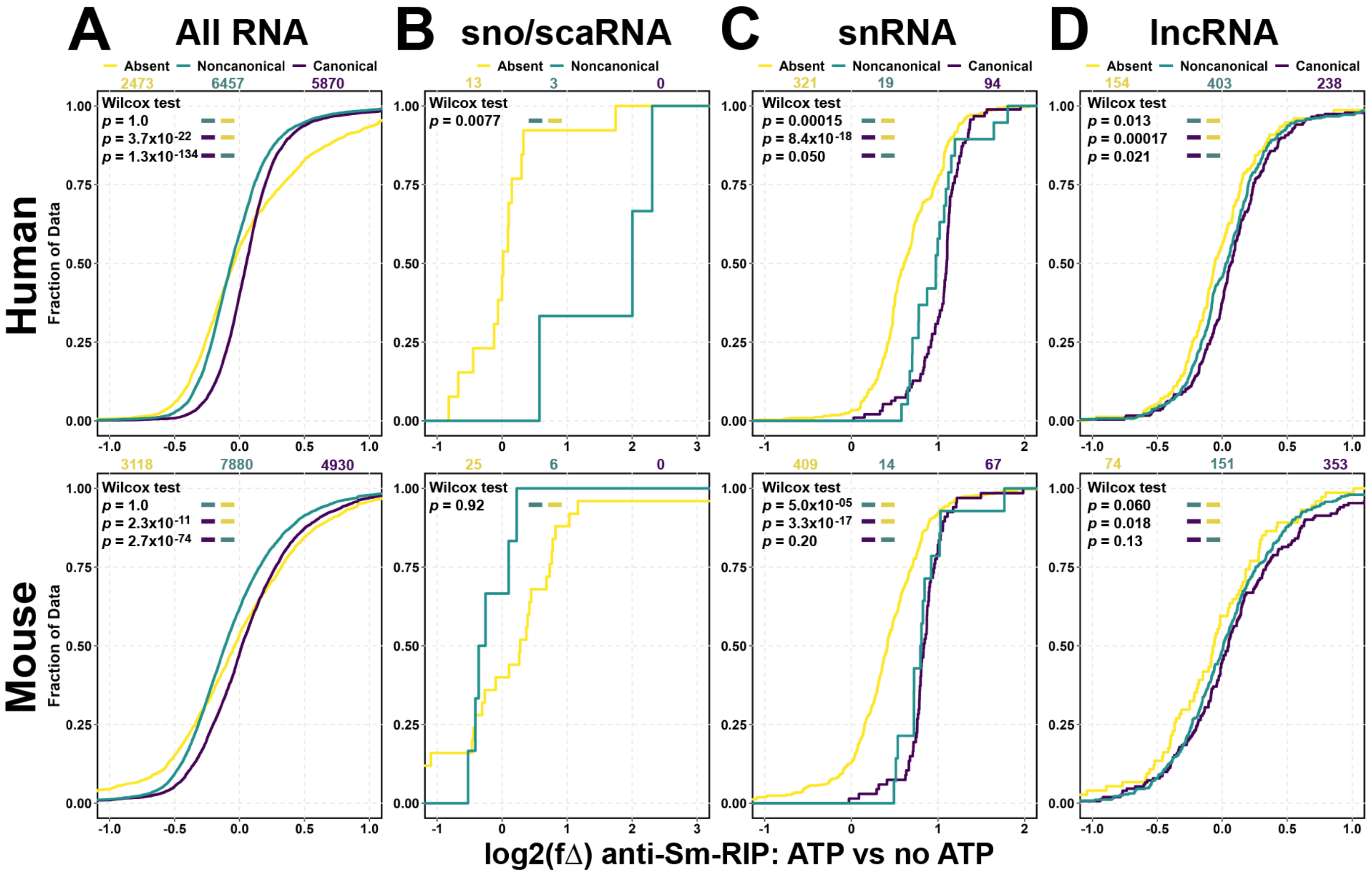


**Supplementary Figure 11: Noncoding RNAs containing Sm-sites are also enriched in anti-Sm-RIPs in an ATP-dependent manner.** For **(A-D)**, **(top)** for human analysis and **(bottom)** for mouse analysis. **(A-C)** Cumulative distribution plots of all RNA **(A)**, only sno/scaRNA **(B)**, snRNA **(C)**, or lncRNA **(D)** delineating by Sm-site prediction—**Absent** (yellow), **Noncanonical** (green), **Canonical** (purple). Values above plots are the number of genes plotted for each group in color. *P-values* for Wilcox tests using the alternative that the left color is greater than the right color are provided in the upper-lefthand corner of the plots.

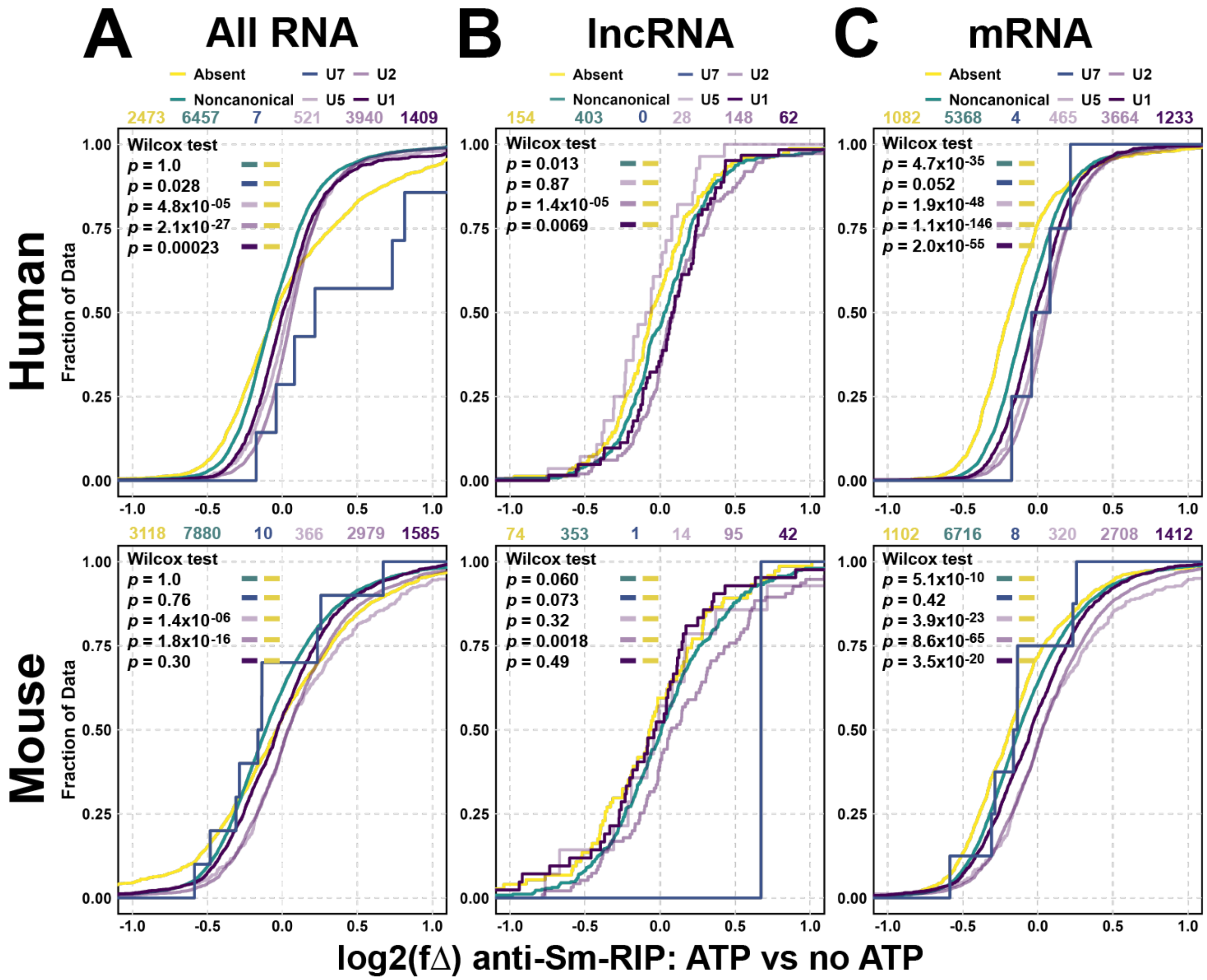


**Supplementary Figure 12: Type of Sm-site correlates with further enrichment following ATP addition and anti-Sm-RIP.** Cumulative distribution plots for given RNA biotypes, plotting the log2 fold change (log2(fΔ)) by increasing value. log2(fΔ) was calculated between the ATP+ and ATP- conditions. (A) All RNA, (B) lncRNA, (C) mRNA. Colors designate whether an Sm-site is **Absent** (yellow), **Noncanonical** (green), **U7** (blue), or **U1**, **U2**, or **U5** (purple, darkest to lightest). Values above CDFs indicate the number of genes plotted for each condition in color. *P-values* generated from Wilcox tests for the left color being greater than the right color are provided in the upper lefthand corner of the graphs.

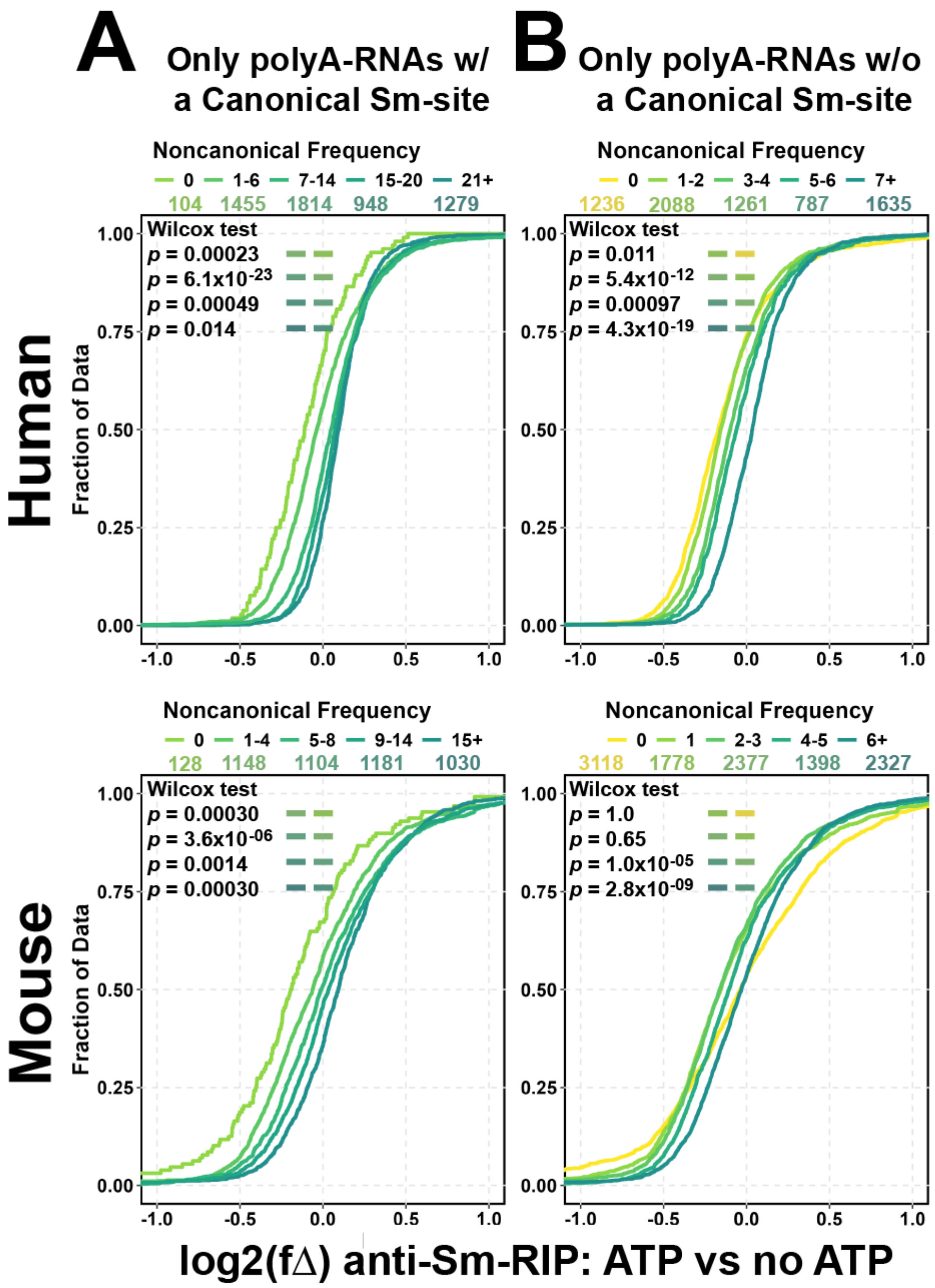


**Supplementary Figure 13. The frequency of Sm-sites in an RNA correlates with further Sm-protein enrichment upon addition of ATP into the assembly reactions. (top)** for human analysis and **(bottom)** for mouse analysis. For each cumulative distribution plot, log2 fold change (log2(fΔ)) between the anti-Sm-RIP supplemented with polyA-RNA and ATP vs without ATP conditions are plotted in increasing order, delineated by the frequency of Sm-sites predicted for the gene product. Values above plots indicate the number of genes plotted for each group designated by color. Wilcox tests were performed using the alternative that the left color is greater than the right color. **(A)** Cumulative distribution plot of only mRNAs predicted to contain at least one canonical Sm-site, delineating the frequency of noncanonical Sm-sites predicted within the mRNA transcript, against the log2 fold change in anti-Sm-RIP supplemented with polyA-RNA and ATP vs anti-Sm-RIPs supplemented only with polyA-RNA. **(B)** Cumulative distribution plot of mRNAs predicted to solely contain noncanonical Sm-sites, delineating the frequency of noncanonical Sm-sites predicted within the mRNA transcript, against the log2 fold change in anti-Sm-RIP supplemented with polyA-RNA and ATP vs anti-Sm-RIPs supplemented only with polyA-RNA.

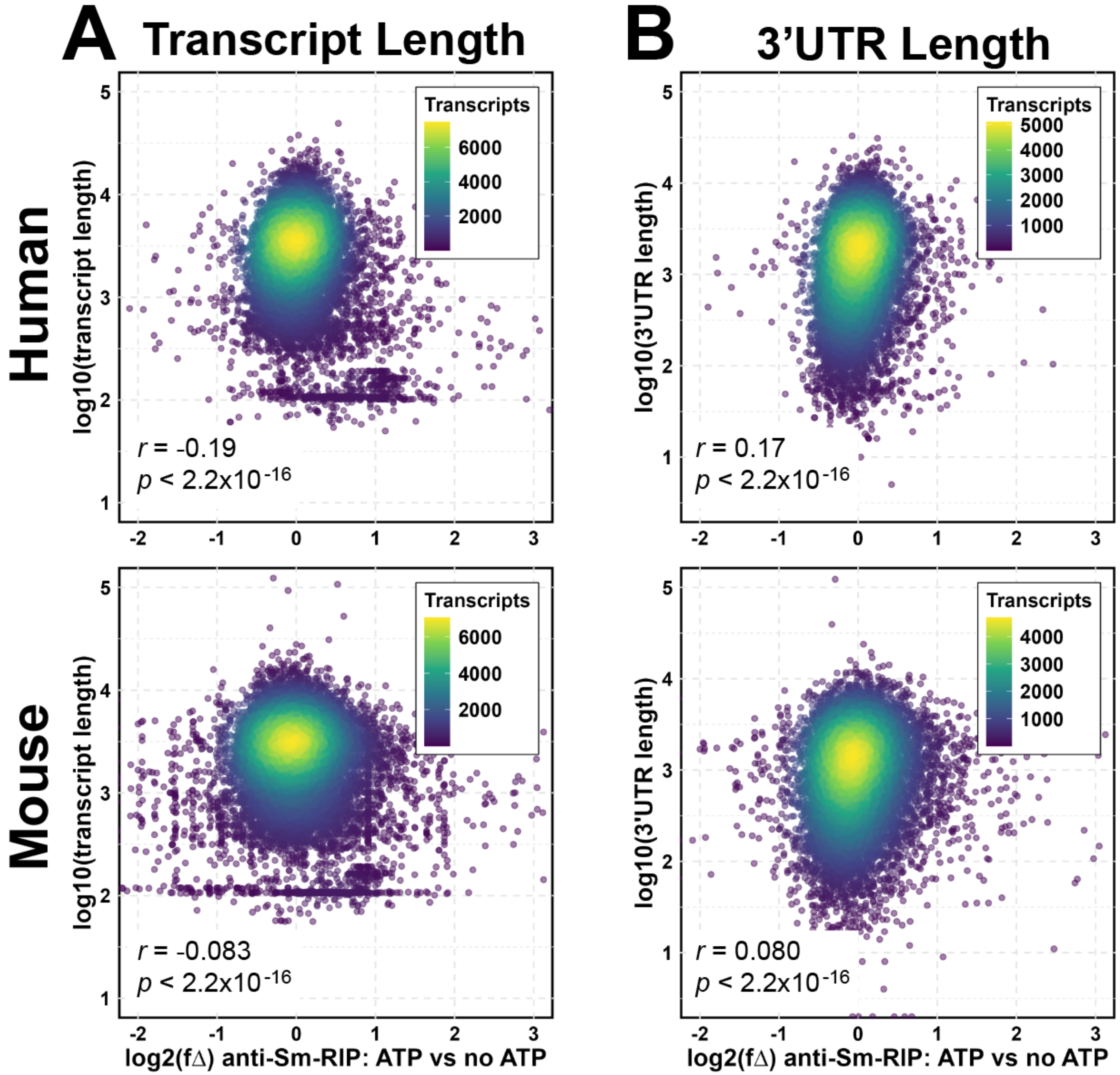


**Supplementary Figure 14: Transcript or 3'UTR length is not a predictor of ATP-dependent enrichment in anti-Sm-RIP experiments. (top)** for human analysis and **(bottom)** for mouse analysis. **(A)** log10 of transcript length plotted against log2 fold change between anti-Sm-RIP supplemented with polyA-RNA and ATP vs anti-Sm-RIPs supplemented only with polyA-RNA. *r* is a Pearson correlation coefficient. **(B)** log10 of 3'UTR length plotted against log2 fold change between anti-Sm-RIP supplemented with polyA-RNA and ATP vs anti-Sm-RIPs supplemented only with polyA-RNA. *r* is a Pearson correlation coefficient.

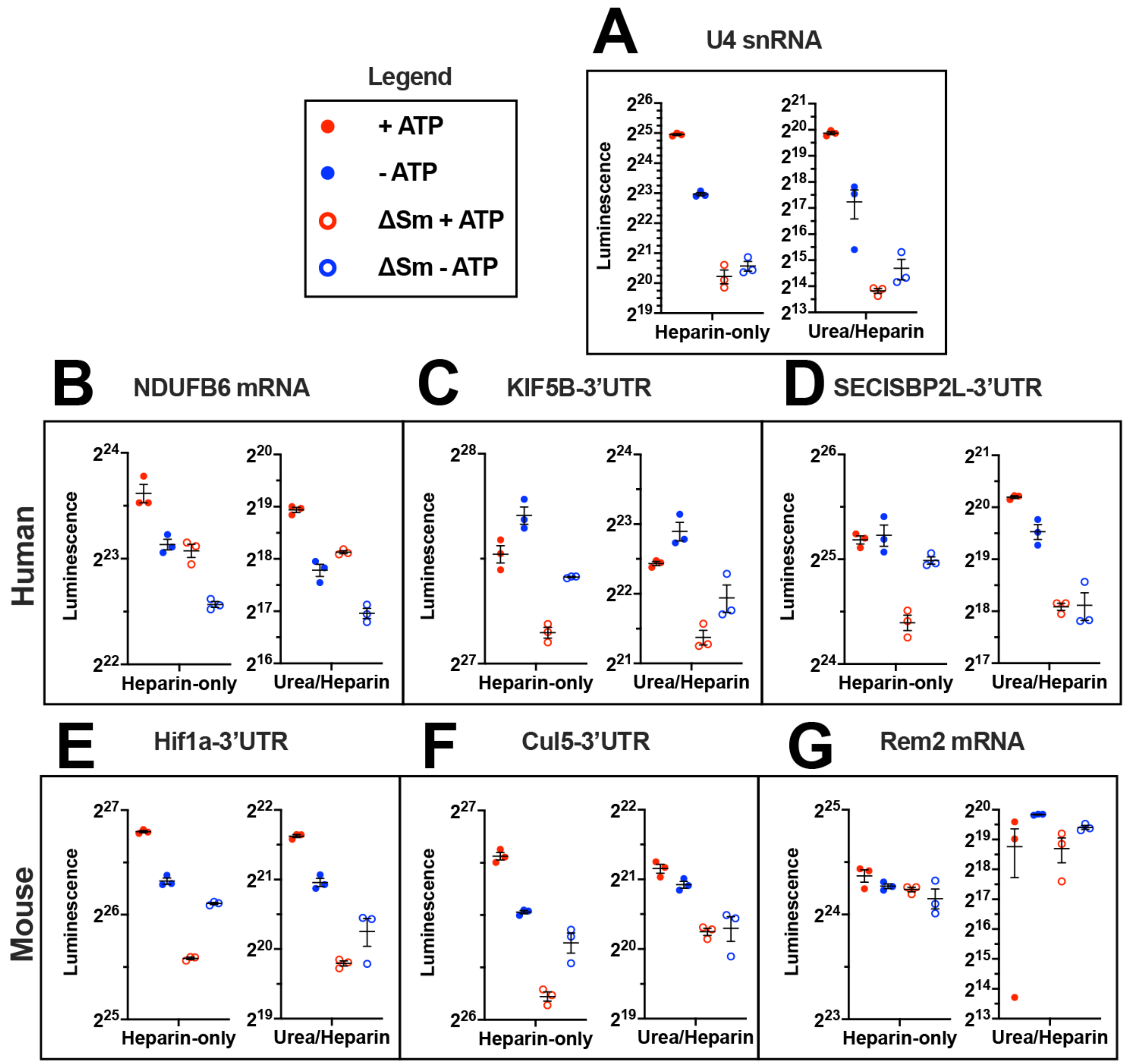


**Supplementary Figure 15: Raw luminescence values for ATP and Sm-site dependent assembly of Sm-protein rings on mRNAs as shown in Figure 4D-J**. **(A-G)** Luminescence results from detection of *in vitro* transcribed, biotin-labelled human U4 snRNA **(A)**, mRNAs (NDUFB6 **(B)** and Rem2 **(G)**), and mRNA 3'UTRs (KIF5B **(C)**, SECISBP2L **(D)**, Hif1a **(E)**, and Cul5 **(F)**) enriched following anti-Sm-RIP. 4 conditions were performed for each RNA: (solid red dot) cytoplasmic cell extract supplemented with wild-type RNA and ATP, (solid blue dot) cytoplasmic cell extract supplemented with wild-type RNA but not with ATP, (open red circle) cytoplasmic cell extract supplemented with ATP and RNA mutated to remove the Sm-site sequence, and (open blue circle) cytoplasmic cell extract supplemented with RNA mutated to remove the Sm-site sequence but not ATP. All shown RNAs have a canonical Sm-site except Rem2. Left graphs are the results obtained by performing the immunoprecipitation in 2 mg/mL heparin, RSB-500 + 0.1% NP-40 and washed 8 times with RSB-500 + 0.1% NP-40. Right graphs are the results obtained following 15 min treatment with 2M urea and 5 mg/mL heparin, followed by immunoprecipitation in 2 mg/mL heparin, RSB-500 + 0.1% NP-40 and washed 8 times with RSB-500 + 0.1% NP-40. Raw luminescence values are comparable between Heparin-only and Urea/Heparin conditions were performed on separate plates. Raw luminescence is comparable for the different RNAs within these groups as they were performed at the same time, using the same reagents, and same plate.

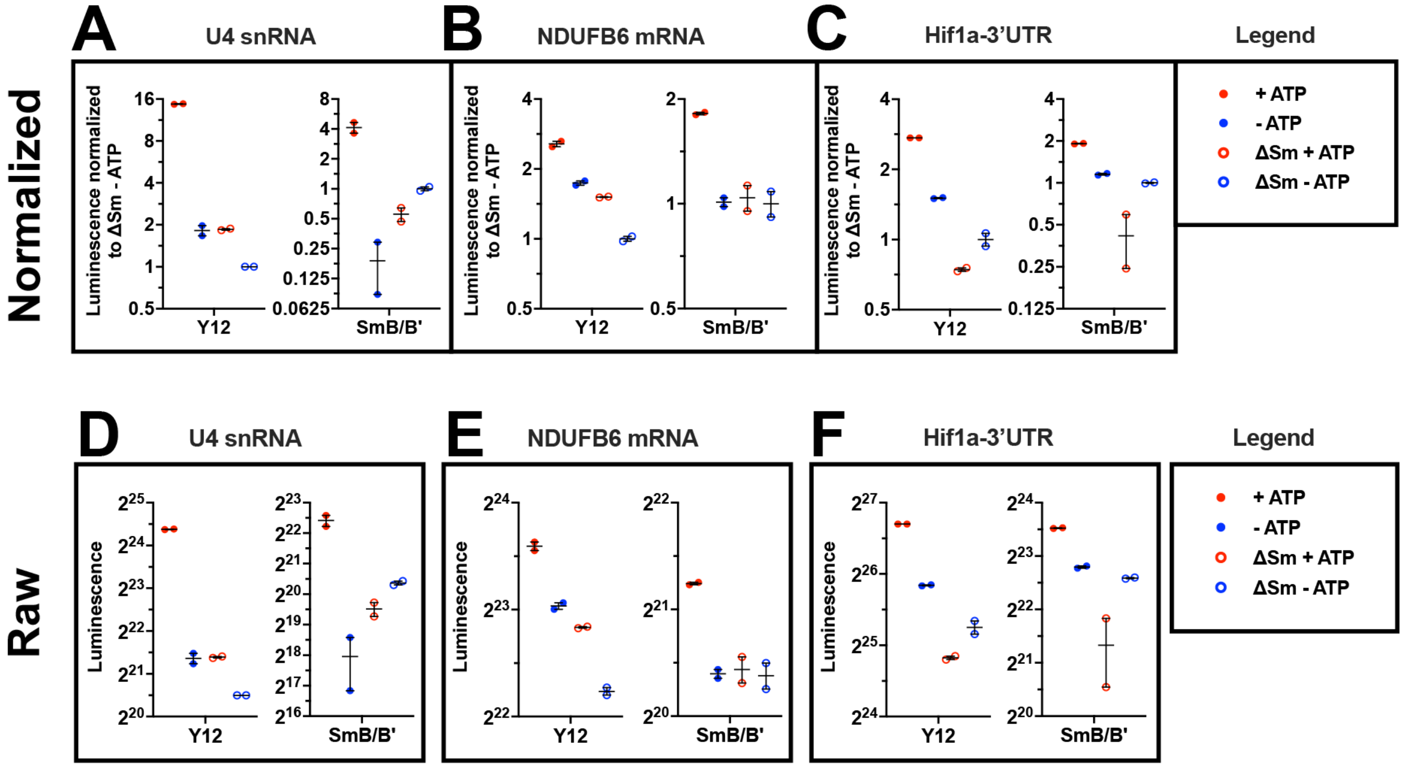


**Supplementary Figure 16: Sm-ring assembly on mRNAs is reproducible with an antibody specific for SmB/B'.** Luminescence results from detection of *in vitro* transcribed, biotin-labelled human U4 snRNA **(AD)**, NDUFB6 mRNA **(BE)** and Hif1a 3'UTR **(CF)** enriched following anti-Sm-RIP. 4 conditions were performed for each RNA: (solid red dot) cytoplasmic cell extract supplemented with wild-type RNA and ATP, (solid blue dot) cytoplasmic cell extract supplemented with wild-type RNA but not with ATP, (open red circle) cytoplasmic cell extract supplemented with ATP and RNA mutated to remove the Sm-site sequence, and (open blue circle) cytoplasmic cell extract supplemented with RNA mutated to remove the Sm-site sequence but not ATP. All shown RNAs have a canonical Sm-site and all reactions were incubated with 2M urea and 5 mg/mL heparin for 15 min, followed by immunoprecipitation in 2 mg/mL heparin, RSB-500 + 0.1% NP-40 and washed 8 times with RSB-500 + 0.1% NP-40. Left graphs are the results obtained by performing the immunoprecipitation with the infamous Y12 antibody, and right was performed using an SmB/B' specific antibody. Raw luminescence values are given on the bottom and luminescence normalized to the ΔSm-ATP condition are given on the top set of graphs.

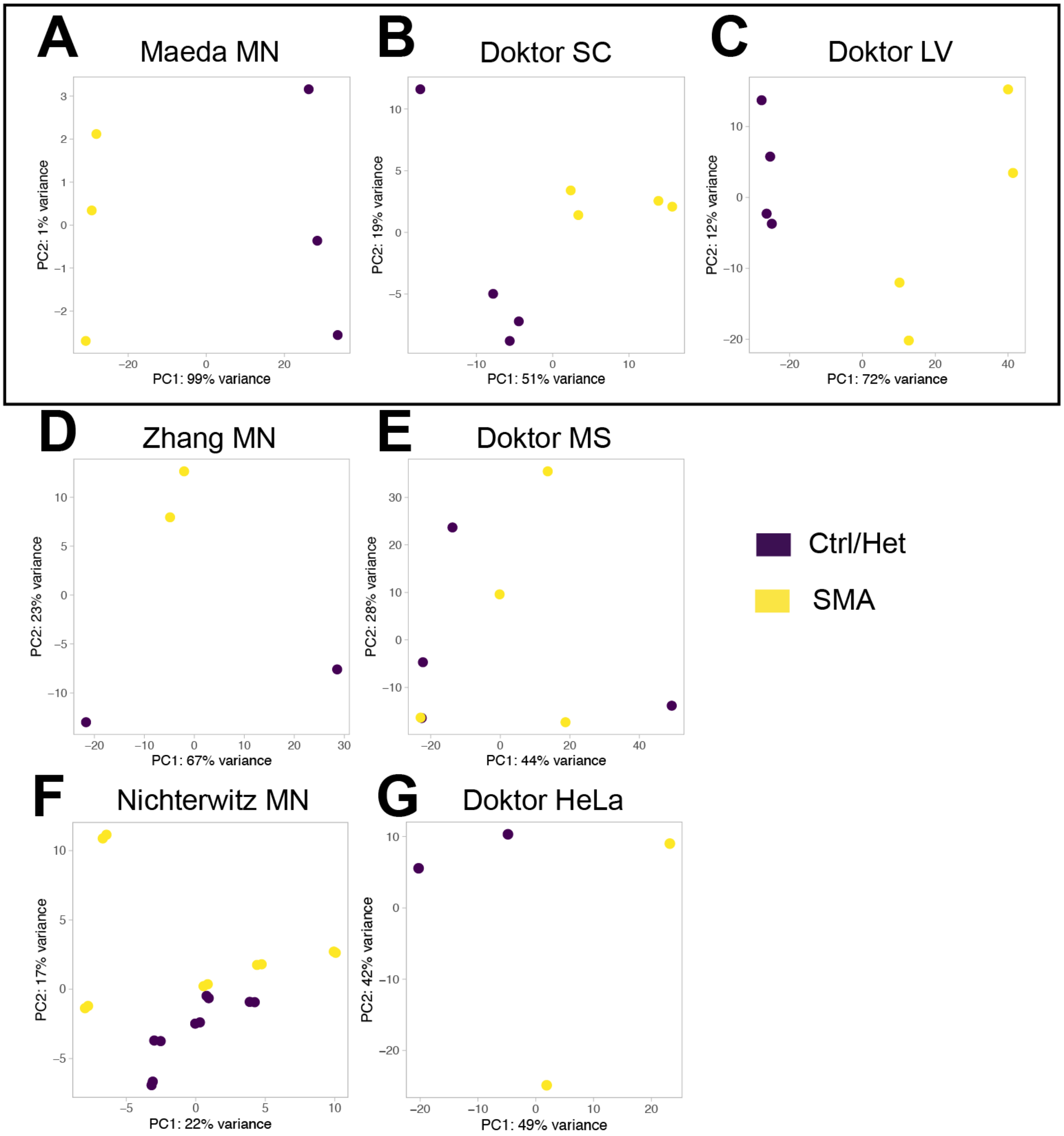


**Supplementary Figure 17: Principal component analyses for external sequencing data used in this study.** Boxed PCA plots **(A-C)** indicate those datasets used for analyses described in Figure 5. Un-boxed PCA plots **(D-G)** were excluded from analysis as the principal variance between samples is not due to the sample genotype and therefore any variations cannot be expected to be due to a decrease in SMN. **(A)** comparison of mESC differentiated motor neurons derived from SMA ($Smn^{+/-}$;$SMN2^{+/+}$) and normal ($Smn^{+/+}$;$SMN2^{+/+}$) as published by Maeda *et al PLoS ONE* 2014 (96). **(B)** comparison of spinal cord lysates collected from post-natal day 5 Taiwanese SMA ($Smn^{-/-}$;$SMN2^{+/+}$) and Taiwanese Het ($Smn^{+/-}$;$SMN2^{+/+}$) mice as published by Doktor *et al NAR* 2017 (97). **(C)** comparison of liver lysates collected from post-natal day 5 Taiwanese SMA ($Smn^{-/-}$;$SMN2^{+/+}$) and Taiwanese Het ($Smn^{+/-}$;$SMN2^{+/+}$) mice as published by Doktor *et al NAR* 2017 (97). **(D)** comparison of post-natal-day 1 Δ7SMA ($Smn^{-/-}$;$SMN2^{+/+}$;SMNΔ$7^{+/+}$) and Δ7Het ($Smn^{+/-}$;$SMN2^{+/+}$;SMNΔ$7^{+/+}$) laser-microdissected mouse motor neurons as published by Zhang *et al PNAS* 2014 (115). **(E)** comparison of muscle lysates collected from post-natal day 5 Taiwanese SMA ($Smn^{-/-}$;$SMN2^{+/+}$) and Taiwanese Het ($Smn^{+/-}$;$SMN2^{+/+}$) mice as published by Doktor *et al NAR* 2017 (97). **(F)** comparison of post-natal-day 5 Δ7SMA ($Smn^{-/-}$;$SMN2^{+/+}$;SMNΔ$7^{+/+}$) and Δ7Het ($Smn^{+/-}$;$SMN2^{+/+}$;SMNΔ$7^{+/+}$) laser-microdissected mouse motor neurons as published by Nichterwitz *et al* (98). **(G)** comparison of siRNA-mediated knockdown of SMN in HeLa cell extracts as published by Doktor *et al NAR* 2017 (97).
