## Supplementary material for "Sm-site containing mRNAs can accept Sm-rings and are downregulated in Spinal Muscular Atrophy": Sequences for mRNA candidates used to test Sm-ring assembly

Sequences used to clone candidates for Sm-ring assembly.

Format:
Gene name, gene_id, cDNA

Region of Sm-site and number of Sm-sites identified, with type (NC for noncanonical, C for canonical).

Wild-type sequences are given first, followed by sequences with Sm-sites removed (ΔSm).

Within sequence:
**Start codon**
**noncanonical smsite**
**Canonical smsite**
**Stop codon**
polyA-site

NDUFB6-203 ENST00000379847.8 cDNA

CDS 2 x NC
3’UTR 3 x NC 1 x C

TAATACGACTCACTATAGGTTCCCGCAAGGTCGCTTTGCAGAGCGGGAGCGCGCTTAAGTAACTAGTCCG
TAGTTCGAGGGTGCGCCGTGTCCTTTTGCGTTGGTACCAGCGGCGAC**ATG**ACGGGGTACACTCCGGATGA
GAAACTGCGGCTGCAGCAGCTGCGAGAGCTGAGAAGGCGATGGCTGAAGGACCAGGAGCTGAGCCCTCGG
GAGCCGGTGCTGCCCCCACAGAAGATGGGGCCTATGGAGAAATTCTGGAATAAATTTTTGGAGAATAAAT
CCCCTTGGAGGAAAATGGTCCATGGGGTATACAAAAAGAGT**ATCTTTG**TTTTCACTCATGTACTTGTACC
TGTCTGGATTATTCATTATTACATGAAGTATCATGTTTCTGAAAAACCATATGGCATAGTTGAAAAGAAG
TCCAGA**ATATTCC**CTGGTGATACAATTCTGGAGACTGGAGAAGTAATTCCACCAATGAAAGAATTTCCTG
ATCAACATCAT**TAA**AGATTATGTAAAAAGTTAAAAGGCTTATGAGCCTAAGTTTGTTCCTATATTACC**AT**
**ATTTA**CTGAATTTTCTGGAAAAGTAACTTTAATAAAGTTTAATCTCAGAAATTGTCAT**ATCTGTT**TTCAA
GCATTGTACAATTTGAGACTGAGTAATTTAACAATAAGTAAAAAGTGGACATGCTAAACAAATATGAGAG
ACTACCTACTTTTTCTGGTCATTCTTGACTTGGAAAACGGTATGGAAAAGTATTTAGTTACATGTTTGTT
TGTTTTTTTCTTACACAGTACTTACACTAATTTGGTATCAGGGTATGCAACAGTGAAATATCACAATAAA
CAAATGTAAGAACAGCAATTCCATGCACTTTTGTTTTAAGGAA**ATCTTTC**Cggccaggcgcagtggctca
tgcctgtaatcccagcactttgggaggccgaggcgggcagatcacgaagtcaggagatcgagaccatcat
ggctaacacagtgaaaccccgtccctactaaaaatacaaaaaaattagccgggcgtggtggcgggctcct
gtagtcccagctactgcggaggctgaggcacgagattggtgtgaacccagaaggcggagcttgcagtaag
ccgagatagtgccactgagcctgggcgacagagcaagactccgtctcaaaaaaaaaaaaaGCTTTCCGAA
TC**ATTTTTG**AAGAATTTAGAAACTTGATTGAAAAGCTTATTCCAACTATGATCTGACACTCAAGACTGTC
AGATTTAGGTTGCTGTTAATTTTGTCATGAGAATGTAAAATACTAAAATCTCTAAGTGAAAAATTTGCAT
ACTAGTGCTTGTTATAAAGGATAATGCAAAAATAAACTTGGGAACTTTGCACGTG

NDUFB6-ΔSm:
TAATACGACTCACTATAGGTTCCCGCAAGGTCGCTTTGCAGAGCGGGAGCGCGCTTAAGTAACTAGTCCG
TAGTTCGAGGGTGCGCCGTGTCCTTTTGCGTTGGTACCAGCGGCGAC**ATG**ACGGGGTACACTCCGGATGA
GAAACTGCGGCTGCAGCAGCTGCGAGAGCTGAGAAGGCGATGGCTGAAGGACCAGGAGCTGAGCCCTCGG
GAGCCGGTGCTGCCCCCACAGAAGATGGGGCCTATGGAGAAATTCTGGAATAAATTTTTGGAGAATAAAT
CCCCTTGGAGGAAAATGGTCCATGGGGTATACAAAAAGAGT**ACCCCCG**TTTTCACTCATGTACTTGTACC
TGTCTGGATTATTCATTATTACATGAAGTATCATGTTTCTGAAAAACCATATGGCATAGTTGAAAAGAAG
TCCAGA**ACCCCCC**CTGGTGATACAATTCTGGAGACTGGAGAAGTAATTCCACCAATGAAAGAATTTCCTG
ATCAACATCAT**TAA**AGATTATGTAAAAAGTTAAAAGGCTTATGAGCCTAAGTTTGTTCCTATATTACC**AC**
**CCCCA**CTGAATTTTCTGGAAAAGTAACTTTAATAAAGTTTAATCTCAGAAATTGTCAT**ACCCCCT**TTCAA
GCATTGTACAATTTGAGACTGAGTAATTTAACAATAAGTAAAAAGTGGACATGCTAAACAAATATGAGAG
ACTACCTACTTTTTCTGGTCATTCTTGACTTGGAAAACGGTATGGAAAAGTATTTAGTTACATGTTTGTT
TGTTTTTTTCTTACACAGTACTTACACTAATTTGGTATCAGGGTATGCAACAGTGAAATATCACAATAAA
CAAATGTAAGAACAGCAATTCCATGCACTTTTGTTTTAAGGAA**ACCCCCC**Cggccaggcgcagtggctca
tgcctgtaatcccagcactttgggaggccgaggcgggcagatcacgaagtcaggagatcgagaccatcat
ggctaacacagtgaaaccccgtccctactaaaaatacaaaaaaattagccgggcgtggtggcgggctcct
gtagtcccagctactgcggaggctgaggcacgagattggtgtgaacccagaaggcggagcttgcagtaag
ccgagatagtgccactgagcctgggcgacagagcaagactccgtctcaaaaaaaaaaaaaGCTTTCCGAA
TC**ACCCCCG**AAGAATTTAGAAACTTGATTGAAAAGCTTATTCCAACTATGATCTGACACTCAAGACTGTC
AGATTTAGGTTGCTGTTAATTTTGTCATGAGAATGTAAAATACTAAAATCTCTAAGTGAAAAATTTGCAT
ACTAGTGCTTGTTATAAAGGATAATGCAAAAATAAACTTGGGAACTTTGCACGTG

KIF5B-201 ENST00000302418.5 cDNA

CDS 3 x NC
3’UTR 9 x NC 2 x C

GTCGCCCAACGGCGGCCTCAGGAGTGATCGGGCAGCAGTCGGCCGGCCAGCGGACGGCAGAGCGGGCGGA
CGGGTAGGCCCGGCCTGCTCTTCGCGAGGAGGAAGAAGGTGGCCACTCTCCCGGTCCCCAGAACCTCCCC
AGCCCCCGCAGTCCGCCCAGACCGTAAAGGGGGACGCTGAGGAgccgcggacgctctccccggtgccgcc
gccgctgccgccgccatggctgccATGATGGATCGGAAGTGAGCATTAGGGTTAACGGCTGCCGGCGCCG
GCTCTTCAAGTCCCGGCTCCCCGGCCGCCTCCACCCGGGGAAGCGCAGCGCGGCGCAGCTGACTGCTGCC
TCTCACGGCCCTCGCGACCACAAGCCCTCAGGTCCGGCGCGTTCCCTGCAAGACTGAGCGGCGGGGAGTG
GCTCCCGGCCGCCGGCCCCGGCTGCGAGAAAG**ATG**GCGGACCTGGCCGAGTGCAACATCAAAGTG**ATGTG**
**TC**GCTTCAGACCTCTCAACGAGTCTGAAGTGAACCGCGGCGACAAGTACATCGCCAAGTTTCAGGGAGAA
GACACGGTCGTGATCGCGTCCAAGCCTTATGCATTTGATCGGGTGTTCCAGTCAAGCACATCTCAAGAGC
AAGTGTATAATGACTGTGCAAAGAAGATTGTTAAAGATGTACTTGAAGGATATAATGGAACAATATTTGC
ATATGGACAAACATCCTCTGGGAAGACACACACAATGGAGGGTAAACTTCATGATCCAGAAGGCATGGGA
ATTATTCCAAGAATAGTGCAAGATATTTTTAATTATATTTACTCCATGGATGAAAATTTGGAATTTCATA
TTAAGGTTTC**ATATTTT**GAAATATATTTGGATAAGATAAGGGACCTGTTAGATGTTTCAAAGACCAACCT
TTCAGTTCATGAAGACAAAAACCGAGTTCCCTATGTAAAGGGGTGCACAGAGCGTTTTGTATGTAGTCCA
GATGAAGTTATGGATACCATAGATGAAGGAAAATCCAACAGACATGTAGCAGTTACAAATATGAATGAAC
ATAGCTCTAGGAGTCACAGTATATTTCTTATTAATGTCAAACAAGAGAACACACAAACGGAACAAAAGCT
GAGTGGAAAACTTTATCTGGTTGATTTAGCTGGTAGTGAAAAGGTTAGTAAAACTGGAGCTGAAGGTGCT
GTGCTGGATGAAGCTAAAAACATCAACAAGTCACTTTCTGCTCTTGGAAATGTTATTTCTGCTTTGGCTG
AGGGTAGTACATATGTTCCATATCGAGATAGTAAAATGACAAGAATCCTTCAAGATTCATTAGGTGGCAA
CTGTAGAACCACTATTGTAATTTGCTGCTCTCCATCATCATACAATGAGTCTGAAACAAAATCTACACTC
TTATTTGGCCAAAGGGCCAAAACAATTAAGAACACAGTTTGTGTCAATGTGGAGTTAACTGCAGAACAGT
GGaaaaagaagtatgaaaaagaaaaagaaaaaaaTAAGATCCTGCGGAACACTATTCAGTGGCTTGAAAA
TGAGCTCAACAGATGGCGTAATGGGGAGACGGTGCCTATTGATGAACAGTTTGACAAAGAGAAAGCCAAC
TTGGAAGCTTTCACAGTGGATAAAGATATTACTCTTACCAATGATAAACCAGCAACCGCAATTGGAGTTA
TAGGAAATTTTACTGATGCTGAAAGAAGAAAGTGTGAAGAAGAAATTGCTAAATTATACAAACAGCTTGA
TGACAAGGATGAAGAAATTAACCAGCAAAGTCAACTGGTAGAGAAACTGAAGACGCAAATGTTGGATCAG
GAGGAGCTTTTGGCATCTACCAGAAGGGATCAAGACAATATGCAAGCTGAGCTGAATCGCCTTCAAGCAG
AAAATGATGCCTCTAAAGAAGAAGTGAAAGAAGTTTTACAGGCCCTAGAAGAACTTGCTGTCAATTATGA
TCAGAAGTCTCAGGAAGTTGAAGACAAAACTAAGGAATATGAATTGCTTAGTGATGAATTGAATCAGAAA
TCGGCAACTTTAGCGAGTATAGATGCTGAGCTTCAGAAACTTAAGGAAATGACCAACCACCAGAAAAAAC
GAGCAGCTGAGATGATGGCATCTTTACTAAAAGACCTTGCAGAAATAGGAATTGCTGTGGGAAATAATGA
TGTAAAGCAGCCTGAGGGAACTGGCATGATAGATGAAGAGTTCACTGTTGCAAGACTCTACATTAGCAAA
ATGAAGTCAGAAGTAAAAACCATGGTGAAACGTTGCAAGCAGTTAGAAAGCACACAAACTGAGAGCAACA
AAAAAATGGAAGAAAATGAAAAGGAGTTAGCAGCATGTCAGCTTCGTATCTCTCAACATGAAGCCAAAAT
CAAGTCATTGACTGAATACCTTCAAAATGTGGAACAAAAGAAAAGACAGTTGGAGGAATCTGTCGATGCC
CTCAGTGAAGAACTAGTCCAGCTTCGAGCACAAGAGAAAGTCCATGAAATGGAAAAGGAGCACTTAAATA
AGGTTCAGACTGCAAATGAAGTTAAGCAAGCTGTTGAACAGCAGATCCAGAGCCATAGAGAAACTCATCA
AAAACAGATCAGTAGTTTGAGAGATGAAGTAGAAGCAAAAGCAAAACTTATTACTGATCTTCAAGACCAA
AACCAGAAAATGATGTTAGAGCAGGAACGTCTAAGAGTAGAACATGAGAAGTTGAAAGCCACAGATCAGG
AAAAGAGCAGAAAACTACATGAACTTACGGTTATGCAAGATAGACGAGAACAAGCAAGACAAGACTTGAA
GGGTTTGGAAGAGACAGTGGCAAAAGAACTTCAGACTTTACACAACCTGCGCAAACTCTTTGTTCAGGAC
CTGGCTACAAGAGTTAAAAAGAGTGCTGAGATTGATTCTGATGACACCGGAGGCAGCGCTGCTCAGAAGC
AAAAAATCTCCTTTCTTGAAAATAATCTTGAACAGCTCACTAAAGTGCACAAACAGTTGGTACGTGATAA
TGCAGATCTCCGCTGTGAACTTCCTAAGTTGGAAAAGCGACTTCGAGCTACAGCTGAGAGAGTGAAAGCT

TTGGAATCAGCACTGAAAGAAGCTAAAGAAAATGCATCTCGTGATCGCAAACGCTATCAGCAAGAAGTAG
ATCGCATAAAGGAAGCAGTCAGGTCAAAGAATATGGCCAGAAGAGGGCATTCTGCACAGATTGCTAAACC
TATTCGTCCCGGGCAACATCCAGCAGCTTCTCCAACTCACCCAAGTGCAATTCGTGGAGGAGGTGC**ATTT**
**GTT**CAGAACAGCCAGCCAGTGGCAGTGCGAGGTGGAGGAGGCAAACAAGTG**TAA**TCGTTTATACATACCC
ACAGGTGTTAAAAAGTAATCGAAGTACGAAGAGGACATGGTATCAAGCAGTCATTCAATGACTATAACCT
CTACTCCCTTGGGATTGTAGAATTATAACTTTTaaaaaaaTGTATAAATTATACCTGGCCTGTACAGCT
GTTTCCTACCTACTCTTCTTGTAAACTCTGCTGCTTCCCAACACAACTAGAGTGCAATTTTGGCATCTTA
GGAGGGAAAAAGGACAGTTTACAACTGTGGCCCTATTTATTACACAGTTTGTCTATCGTGTCTTAAATTT
AGTCTTTACTGTGCCAAGCTAACTGTACCTTATAGGACTGTACtttttgt**atttttttg**tgtatgttt**att**
**ttttA**ATCTCAGTTTAAATTACCTAGCTGCTACTGCTTCTTGTTTTTCTTTTCCTATTAAAACGTCTTCC
tttttttttCTTAAGAGAAAATGGAACATTTAGGTTAAATGTCTTTAAATTTTACCACTTAACAACACTA
CATGCCCATAAAATATATCCAGTCAGTACTGTATTTTAAAATCCCTTGAAATGATGATATCAGGGTTAAA
ATTACTTGTATTGTTTCTGAAGTTTGCTCCTGAAAACTACTGTTTGAGCACTGAAACGTTACAAATGCCT
AATAGGCATTTGAGACTGAGCAAGGCTACTTGTTATCTCATGAAATGCCTGTTGCCGAGTTATTTTGAAT
AGAAATATTTTAAAGTATCAAAAGCAGATCTTAGTTTAAGGGAGTTTGGAAAAGGAATTATATTTCTCTT
TTTCCTGATTCTGTACTCAACAAGTCTTGATGGAATTAAAATACTCTGCTTTATTCTGGTGAGCCTGCTA
GCTAATATAAGTATTGGACAGGTAATAATTTGTCATCTTTAATATTAGTAAAATGAATTAAGATATTATA
GGATTAAACATAATTTTATACGGTTAGTACTTTATTGGCCGACCTAAATTTATAGCGTGTGGAAATTGAG
AAAAATGAAGAAACAGGACAGATATATGATGAATTAAAAATATATATAGGTCAATTTTGGTCTGAAATCC
CTGAGGTGTTTTTAACCTGCTACACTA**ATTTGTA**CACTAATTTATTTCTTTAGTCTAGAAATAGTAAATT
GTTTGCAAGTCACTAATAATCATTAGATAAATTATTTTCTTGGCCATAGCCGATAATTTTGTAATCAGTA
CTAAGTGTATACGT**ATTTTTG**CCACTTTTTCCTCAGATGATTAAAGTAAGTCAACAGCTTATTTTAGGAA
ACTGTAAAAGTAATAGGGAAAGAGATTTCACTATTTGCTTCATCAGTGGTAGGGGGGCGGTGACTGCAAC
TGTGTTAGCAGAAATTCACAGAGAATGGGGATTTAAGGTTAGCAGAGAAACTTGGAAAGTTCTGTGTTAG
GATCTTGCTGGCAGAATTAACTTTTTGCAAAAGTTTTATACACAGATATTTGTATTAAATTTGGAGCCAT
AGTCAGAAGACTCAGATCATAATTGGCTTATTTTTCTATTTCCGTAACTATTGTAATTTCCACTTTTGta
ataattttgatttaaaatataaatttatttattt**attttttt**taataGTCAAAA**ATCTTTG**CTGTTGTAGT
CTGCAACCTCTAAAATGATTGTGTTGCTTTTAGGATTGATCAGAAGAAACACTCCAAAAATTGAGATGAA
ATGTTGGTGCAGCCAGTTATAAGTAATATAGTTAACAAGCAAAAAAAGTGCTGCCACCTTTTATGATGAT
TTTCTAAATGGAGAAACATTTGGCTGCATCCACATAGACCTTT**ATGTTTT**GTTTTCAGTTGAAAACTTGC
CTCCTTTGGCAACATTCGTAAATGAAGCAGAAtttttttttctctttttttCCAA**ATATGTT**AGTTTTGTT
CTTGTAAGATGTATCATGGGTATTGGTGCTGTGTAATGAACAACGAATTTTAATTAGCATGTGGTTCAGA
ATATACAATGTTAGGTTTTTAAAAAGTATCTTGATGGTTCTTTTCTATTTATAATTTCAGACTTTCATAA
AGTGTACCAAGAATTTCATAAATTTGTTTTCAGTGAACTGCTTTTTGCTATGGTAGGTCATTAAACACAG
CACTTACTCTTAAAAATGAAAATTTCTGATCATCTAGGATATTGACACATTTCAATTTGCAGTGTCTTTT
TGACTGGATATATTAACGTTCCTCTGAATGGCATTGATAGATGGTTCAGAAGAGAAACTCAATGAAATAA
AGAGA**ATATTTA**TTCATGGCGATTAATTAAATTATTTGCCTAACTTAAGAAAACTACTGTGCGTAACTCT
CAGTTTGTGCTTAACTCCATTTGACATGAGGTGACAGAAGAGAGTCTGAGTCTACCTGTGGA**ATATGTT**G
GTTTATTTTCAGTGCTTGAAGATACATTCACAAATACTTGGTTTGGGAAGACACCGTTTAATTTTAAGTT
AACTTGCATGTTGTAAATGCGTTTT**ATGTTTA**AATAAAGAGGAAAATTTTTTGAAA

KIF5B-3’UTR:
TAATACGACTCACTATAGGTTTATACATACCCACAGGTGTTAAAAAGTAATCGAAGTACGAAGAGGACAT
GGTATCAAGCAGTCATTCAATGACTATAACCTCTACTCCCTTGGGATTGTAGAATTATAACTTTTaaaaa
aaaTGTATAAATTATACCTGGCCTGTACAGCTGTTTCCTACCTACTCTTCTTGTAAACTCTGCTGCTTCC
CAACACAACTAGAGTGCAATTTTGGCATCTTAGGAGGGAAAAAGGACAGTTTACAACTGTGGCCCTATTT
ATTACACAGTTTGTCTATCGTGTCTTAAATTTAGTCTTTACTGTGCCAAGCTAACTGTACCTTATAGGAC
TGTACtttttgt**atttttttg**tgtatgttt**attttttA**ATCTCAGTTTAAATTACCTAGCTGCTACTGCTT
CTTGTTTTTCTTTTCCTATTAAAACGTCTTCCtttttttttCTTAAGAGAAAATGGAACATTTAGGTTAA

ATGTCTTTAAATTTTACCACTTAACAACACTACATGCCCATAAAATATATCCAGTCAGTACTGTATTTTA
AAATCCCTTGAAATGATGATATCAGGGTTAAAATTACTTGTATTGTTTCTGAAGTTTGCTCCTGAAAACT
ACTGTTTGAGCACTGAAACGTTACAAATGCCTAATAGGCATTTGAGACTGAGCAAGGCTACTTGTTATCT
CATGAAATGCCTGTTGCCGAGTTATTTTGAATAGAAATATTTTAAAGTATCAAAAGCAGATCTTAGTTTA
AGGGAGTTTGGAAAAGGAATTATATTTCTCTTTTTCCTGATTCTGTACTCAACAAGTCTTGATGGAATTA
AAATACTCTGCTTTATTCTGGTGAGCCTGCTAGCTAATATAAGTATTGGACAGGTAATAATTTGTCATCT
TTAATATTAGTAAAATGAATTAAGATATTATAGGATTAAACATAATTTTATACGGTTAGTACTTTATTGG
CCGACCTAAATTTATAGCGTGTGGAAATTGAGAAAAATGAAGAAACAGGACAGATATATGATGAATTAAA
AATATATATAGGTCAATTTTGGTCTGAAATCCCTGAGGTGTTTTTAACCTGCTACACTA**ATTTGTA**CACT
AATTTATTTCTTTAGTCTAGAAATAGTAAATTGTTTGCAAGTCACTAATAATCATTAGATAAATTATTTT
CTTGGCCATAGCCGATAATTTTGTAATCAGTACTAAGTGTATACGT**ATTTTTG**CCACTTTTTCCTCAGAT
GATTAAAGTAAGTCAACAGCTTATTTTAGGAAACTGTAAAAGTAATAGGGAAAGAGATTTCACTATTTGC
TTCATCAGTGGTAGGGGGGCGGTGACTGCAACTGTGTTAGCAGAAATTCACAGAGAATGGGGATTTAAGG
TTAGCAGAGAAACTTGGAAAGTTCTGTGTTAGGATCTTGCTGGCAGAATTAACTTTTTGCAAAAGTTTTA
TACACAGATATTTGTATTAAATTTGGAGCCATAGTCAGAAGACTCAGATCATAATTGGCTTATTTTTCTA
TTTCCGTAACTATTGTAATTTCCACTTTTGtaataatttttgatttaaaatataaatttatttattt**attt**
**ttt**taataGTCAAAA**ATCTTTG**CTGTTGTAGTCTGCAACCTCTAAAATGATTGTGTTGCTTTTAGGATTG
ATCAGAAGAAACACTCCAAAAATTGAGATGAAATGTTGGTGCAGCCAGTTATAAGTAATATAGTTAACAA
GCAAAAAAAGTGCTGCCACCTTTTATGATGATTTTCTAAATGGAGAAACATTTGGCTGCATCCACATAGA
CCTTT**ATGTTTT**GTTTTCAGTTGAAAACTTGCCTCCTTTGGCAACATTCGTAAATGAAGCAGAAtttttt
tttctctttttttCCAA**ATATGTT**AGTTTTGTTCTTGTAAGATGTATCATGGGTATTGGTGCTGTGTAATG
AACAACGAATTTTAATTAGCATGTGGTTCAGAATATACAATGTTAGGTTTTTAAAAAGTATCTTGATGGT
TCTTTTCTATTTATAATTTCAGACTTTCATAAAGTGTACCAAGAATTTCATAAATTTGTTTTCAGTGAAC
TGCTTTTTGCTATGGTAGGTCATTAAACACAGCACTTACTCTTAAAAATGAAAATTTCTGATCATCTAGG
ATATTGACACATTTCAATTTGCAGTGTCTTTTTGACTGGATATATTAACGTTCCTCTGAATGGCATTGAT
AGATGGTTCAGAAGAGAAACTCAATGACACGTG

KIF5B-3’UTR-ΔSm:
TAATACGACTCACTATAGGTTTATACATACCCACAGGTGTTAAAAAGTAATCGAAGTACGAAGAGGACAT
GGTATCAAGCAGTCATTCAATGACTATAACCTCTACTCCCTTGGGATTGTAGAATTATAACTTTTaaaaa
aaaTGTATAAATTATACCTGGCCTGTACAGCTGTTTCCTACCTACTCTTCTTGTAAACTCTGCTGCTTCC
CAACACAACTAGAGTGCAATTTTGGCATCTTAGGAGGGAAAAAGGACAGTTTACAACTGTGGCCCTATTT
ATTACACAGTTTGTCTATCGTGTCTTAAATTTAGTCTTTACTGTGCCAAGCTAACTGTACCTTATAGGAC
TGTACtttttgt**aCtCtCtg**tgtatgttt**aCtCtCtA**ATCTCAGTTTAAATTACCTAGCTGCTACTGCTT
CTTGTTTTTCTTTTCCTATTAAAACGTCTTCCtttttttttCTTAAGAGAAAATGGAACATTTAGGTTAA
ATGTCTTTAAATTTTACCACTTAACAACACTACATGCCCATAAAATATATCCAGTCAGTACTGTATTTTA
AAATCCCTTGAAATGATGATATCAGGGTTAAAATTACTTGTATTGTTTCTGAAGTTTGCTCCTGAAAACT
ACTGTTTGAGCACTGAAACGTTACAAATGCCTAATAGGCATTTGAGACTGAGCAAGGCTACTTGTTATCT
CATGAAATGCCTGTTGCCGAGTTATTTTGAATAGAAATATTTTAAAGTATCAAAAGCAGATCTTAGTTTA
AGGGAGTTTGGAAAAGGAATTATATTTCTCTTTTTCCTGATTCTGTACTCAACAAGTCTTGATGGAATTA
AAATACTCTGCTTTATTCTGGTGAGCCTGCTAGCTAATATAAGTATTGGACAGGTAATAATTTGTCATCT
TTAATATTAGTAAAATGAATTAAGATATTATAGGATTAAACATAATTTTATACGGTTAGTACTTTATTGG
CCGACCTAAATTTATAGCGTGTGGAAATTGAGAAAAATGAAGAAACAGGACAGATATATGATGAATTAAA
AATATATATAGGTCAATTTTGGTCTGAAATCCCTGAGGTGTTTTTAACCTGCTACACTA**ACTCGCA**CACT
AATTTATTTCTTTAGTCTAGAAATAGTAAATTGTTTGCAAGTCACTAATAATCATTAGATAAATTATTTT
CTTGGCCATAGCCGATAATTTTGTAATCAGTACTAAGTGTATACGT**ACTCTCG**CCACTTTTTCCTCAGAT
GATTAAAGTAAGTCAACAGCTTATTTTAGGAAACTGTAAAAGTAATAGGGAAAGAGATTTCACTATTTGC
TTCATCAGTGGTAGGGGGGCGGTGACTGCAACTGTGTTAGCAGAAATTCACAGAGAATGGGGATTTAAGG
TTAGCAGAGAAACTTGGAAAGTTCTGTGTTAGGATCTTGCTGGCAGAATTAACTTTTTGCAAAAGTTTTA

TACACAGATATTTGTATTAAATTTGGAGCCATAGTCAGAAGACTCAGATCATAATTGGCTTATTTTTCTA
TTTCCGTAACTATTGTAATTTCCACTTTTGtaataattttgatttaaaatataaatttatttattt**aCtC**
**tCt**taataGTCAAAA**ACCCTCG**CTGTTGTAGTCTGCAACCTCTAAAATGATTGTGTTGCTTTTAGGATTG
ATCAGAAGAAACACTCCAAAAATTGAGATGAAATGTTGGTGCAGCCAGTTATAAGTAATATAGTTAACAA
GCAAAAAAAGTGCTGCCACCTTTTATGATGATTTTCTAAATGGAGAAACATTTGGCTGCATCCACATAGA
CCTTT**ACGCTCT**GTTTTCAGTTGAAAACTTGCCTCCTTTGGCAACATTCGTAAATGAAGCAGAAtttttt
tttctctttttttCCAA**ACACGCT**AGTTTTGTTCTTGTAAGATGTATCATGGGTATTGGTGCTGTGTAATG
AACAACGAATTTTAATTAGCATGTGGTTCAGAATATACAATGTTAGGTTTTTAAAAAGTATCTTGATGGT
TCTTTTCTATTTATAATTTCAGACTTTCATAAAGTGTACCAAGAATTTCATAAATTTGTTTTCAGTGAAC
TGCTTTTTGCTATGGTAGGTCATTAAACACAGCACTTACTCTTAAAAATGAAAATTTCTGATCATCTAGG
ATATTGACACATTTCAATTTGCAGTGTCTTTTTGACTGGATATATTAACGTTCCTCTGAATGGCATTGAT
AGATGGTTCAGAAGAGAAACTCAATGACACGTG

SECISBP2L-209 ENST00000559471.6 cDNA

CDS 5 x NC
3’UTR 12 x NC 3 x C

AGTGGCGTAGCCGAATCGGTGTCGCGGCCAGCCAGATAGGGGCGGAGGTCCGGAACCCAGTCTGGACCCG
AGCGGGGGGCCATGGAGAAAGCGGCCCGAGGCGCTGTTTACACCGACTAGCGCGGGCCCGTTGCGGCTGC
AGGCACC**ATG**GACCGAGCCCCCACGGAGCAGAATGTCAAGCTGTCAGCTGAGGTGGAGCCATTTATTCCC
CAGAAGAAGAGTCCTGATACATTTATGATCCCTATGGCTCTCCCAAATGATAATGGAAGTGTTTCTGGTG
TGGAACCAACTCCAATTCCCAGCTACCTGATTACTTGTTACCC**ATTTGTG**CAGGAAAACCAGTCCAATAG
ACAGTTTCCTTTATATAACAATGATATACGATGGCAACAACCCAATCCAAACCCTACTGGACCATACTTT
GCCTATCCCATTATATCTGCTCAGCCGCCTGTTTCTACAGAGTATACATATTATCAGCTGATGCCAGCAC
CATGTGCCCAGGTTATGGGTTTCTATCATCCTTTTCCTACACCTTACTCCAACACCTTTCAGGCTGCAAA
TACTGTAAATGCTATCACCACAGAATGCACTGAGCGTCCAAGTCAGCTTGGACAGGTCTTCCCATTGTCC
AGCCATCGAAGCAGAAACAGTAACAGAGGATCAGTGGTCCCAAAACAACAGCTTTTACAACAGCACATAA
AAAGCAAAAGGCCGCTGGTGAAAAATGTAGCTACTCAGAAAGAAACAAATGCAGCAGGTCCTGATAGTCG
ATCAAAAATTGTGCTTCTGGTAGATGCTTCACAGCAAACTGATTTCCCATCAGATATCGCTAACAAGTCT
CTCTCAGAGACCACTGCAACAATGCTCTGGAAGTCCAAGGGCAGGAGAAGAAGAGCATCCCACCCTACTG
CTGAATCTTCTAGTGAGCAGGGGGCTAGTGAAGCCGACATTGACAGTGATAGTGGTTACTGCAGTCCCAA
ACACAGCAACAACCAGCCTGCAGCAGGGGCTTTGAGAAATCCTGATTCTGGGACCATGAATCATGTGGAA
TCATCT**ATGTGTG**CAGGTGGTGTAAATTGGTCCAATGTAACTTGCCAGGCAACTCAGAAAAAACCTTGGA
TGGAAAAAAATCAGACATTTTCTAGAGGTGGAAGGCAAACTGAACAAAGAAATAATTCACAGGTTGGATT
CAGATGCCGAGGACACAGTACTTCCTCAGAAAGAAGACAGAATTTGCAAAAGAGACCAGATAATAAGCAT
TTAAGCTCTAGTCAATCCCATAGAAGCGATCCAAATTCTGAGTCTTTATATTTTGAGGATGAAGATGGGT
TTCAAGAACTAAATGAGAATGGAAATGCTAAGGATGAGAATATTCAACAAAAACTTTCTTCTAAAGTATT
GGATGATTTACCTGAAAACTCACCAATCAATATAGTTCAGACTCCAATTCCTATTACCACCTCAGTTCCC
AAACGTGCAAAAAGTCAGAAGAAGAAAGCTTTAGCAGCAGCCCTTGCCACAGCTCAAGAGTATTCAGAAA
TAAGTATGGAGCaaaaaaaaTTACAGGAAGCTTTATCAAAAGCAGCTGGAAAAAAGAATAAAACACCTGT
GCAGCTAGATTTAGGGGACATGTTAGCTGCTCTGGAAAAACAACAGCAAGCAATGAAAGCACGGCAAATT
ACTAACACCAGACCTCTGTCATATACAGTGGTTACTGCAGCTTCTTTTCACACTAAAGACTCTACTAATA
GAAAACCTTTAACCAAAAGTCAGCCCTGTTTGACATCCTTTAATTCTGTGGACATTGCTTCTTCTaaagc
aaaaaaaggaaaagagaaggaaattgcaaaactaaaacGACCCACAGCACTTAAAAAGGTTATTTTAAAA
GAAAGAGAGGAAAAGAAGGGGCGCTTAACTGTGGACCACA**ATCTTTT**GGGATCCGAGGAACCAACAGAAA
TGCACTTAGATTTTATTGATGACTTGCCACAGGAGATTGTTTCCCAGGAAGATACTGGACTAAGCATGCC
CAGTGATACTTCACTCTCTCCAGCAAGTCAGAACTCTCCATACTGTATGACACCTGTGTCACAAGGCTCT
CCTGCTAGTTCTGGAATAGGCAGTCCAATGGCATCTTCAACAATAACCAAAATCCACAGCAAAAGATTTA
GAGAGTATTGTAATCAGGTTCTTTGTAAAGAGATTGATGA**ATGTGTG**ACTCTTCTTCTCCAAGAGCTTGT
CAGTTTCCAGGAACGCATCTACCAAAAAGATCCTGTAAGAGCAAAAGCAAGGAGACGACTCGTTATGGGT
CTAAGAGAAGTTACCAAACATATGAAGTTAAACAAGATCAAGTGTGTTATAATTTCTCCAAACTGTGAAA
AAATCCAGTCAAAAGGTGGTCTGGATGAGGCTCTCTATAATGTTATAGCCATGGCACGGGAACAAGAAAT
TCCTTTTGTGTTTGCCCTTGGAAGGAAAGCTCTAGGACGCTGTGTGAACAAGCTGGTTCCTGTTAGCGTA
GTGGGAATCTTCAACTACTTTGGTGCTGAGAGCCTGTTTAATAAATTAGTAGAACTCACTGAGGAGGCCA
GGAAAGCATATAAAGATATGGTTGCAGCAATGGAACAGGAGCAGGCTGAGGAAGCCTTAAAGAATGTGAA
GAAGGTACCACACCACATGGGACATTCTCGGAATCCCTCTGCAGCAAGTGCCATTTCTTTCTGCAGTGTT
ATTTCTGAACCGATCTCTGAAGTAAATGAAAAGGAATATGAAACAAATTGGAGAAACATGGTGGAAACTT
CAGATGGACTGGAAGCATCAGAAAATGAGAAAGAGGTATCCTGTAAGCACAGCACTTCTGAAAAACCCAG
TAAACTTCCATTTGACACACCCCCAATTGGTAAGCAGCCATCATTAGTGGCTACAGGCAGTACTACCTCA
GCTACAAGTGCTGGGAAATCCACAGCAAGTGATAAAGAGGAAGTGAAGCCAGATGACCTGGAATGGGCCT
CACAGCAGAGTACAGAGACTGGCTCTTTGGATGGCAGTTGCCGAGATCTTTTGAATTCCTCCATCACCAG

CACCACCAGCACTCTTGTACCTGGCATGCTTgaagaagaagaagatgaagatgaggaggaggaggaagaT
TATACTCATGAACCCAT**ATCTGTA**GAAGTGCAGCTCAATAGTAGAATTGAGTCTTGGGTCTCAGAGACCC
AGAGAACTATGGAAACCCTTCAGCTTGGAAAAACCCTTAATGGTTCTGAGGAAGACAATGTAGAGCAAAG
TGGAGAAGAGGAAGCAGAGGCGCCTGAGGTGCTGGAGCCAGGGATGGACAGTGAGGCATGGACTGCTGAC
CAGCAGGCCAGTCCTGGGCAGCAGAAGTCCAGCAACTGCAGCTCGCTCAACAAAGAGCACTCTGATTCTA
ATTACACAACGCAAACTACG**TAA**CTCAGGAAATGTCGGCTCTCTATCTCCAGCTGTGGAAGGGTTGCAGC
CATTACCTTTTATGCTTCATCTCAACATTTTGCACTGTCCAGTATTTAATATACGTATTTAATTCCCAAC
AAAT**ATTTTTG**TAGCTTTTACTTGTTATG**ATCTGTA**GCTTAGCTTTTAATTAGTATCTAAGTGTCTTTCT
AAGAACTGTGTGGAAAATTCAG**ATCTGTT**TCAGCTTATTTTGTAATCAAAAACAGTGATAAAAAGAAGAC
CAGATCTTAAAGAAAATAAATTTCAAATGCTTACTTAAAAGACATTTTGAAAGTTAAAGAACAAGGTTCT
AAGGATAGAAGCAGTTATCAGTGTTTGCTTCAGGACTCCACCTCCTCTACTCTAATTTGACCAAAAAATT
GTTTGGGCTTCTTTAAAAAAGAACTGGGGGTGGAGTCAGAAAATTAAATGAAAGGCTGAGGGTAACTAAG
TCCACCAGTGTTGTATGTTAAAAAATCAATGCAACTTTTATGTGGTCCACAA**ATGTTTA**GTCAGAAGTCA
CTGATTATTGTAATTAATTAGTGTTGGGATGGGCTAAAACAGAGCCTTCAAAACTTCGGCTAGCAGTGGA
GCCACCATCTTAGATTATAGCTAGCTAGCCTC**ATTTGTG**GAAAGTGATAGATGCTGTCTATAATAGTGAA
CAGTCACCCATGATAGGACCTCCAGGTTCTGTCTCATATTTGCTTCTTACTTACCTCAGGAATGCTCTTG
TACATAGACTTATTTACAAAAAGCTAGGCACATGTTGACAGGTGAATAACTGTAACCGATTGTATGACTG
CTGCACTTACATGTAAACTCTTCAGAAACAGAGTCTTATACTGGTGTGTTCTCTTGCATGCTTCTGGTTC
AGGACTCTTGATTTGAGATATGGATTTGATTGAGTATCCAAACTTGTCCTGAGTGCAAAACTGTTTCACC
TTTTAAAAAATACCTATTTTGCACCTAGCCTTGAGCACCTTCCACATAGCAATGACCATAGTTACTGTCA
GGAGGTCAAGGAAAGGAACTTTGCACAACTTGTGACATGTATCCTGATAATCAAGGCTTAGAGGAGGAAG
TTTTAGAAGATAAGAGAAAGTTGTTCTAATTGTGCTGAAACTATTAGATGATTTAGAGTATACAGATATG
TAGGTATTAATTCTCTATTCACTATTATTTATCTCTGCCCTTCTCTAGGAGTTTGTATACCTGCTTAGGA
GACAATAAATGAGCTAA**ATGTTTT**ATTTGCTAGTCAGTCACCACCTGGACTTCAGTGACTTTACAAGTTT
ATGTAATGGTGGAAGAATGACAAACTATGTAATTTTTTTGTCTTCCATccaactccccaccacccccaac
tgtcccccccaccccccTCACACACATGCACACATCCGTACGTGTGTGTGTTTTCCACTTACAAGCTTCC
ATAAGCAGGCACAAAACTGAGAAGGAAGGGGTATTATCCCTGCCCTGATTATCTGGGGCAGGGCTTTGCC
TCACAGAGGCAGGAGAGAAGAATTGGGCAGATTCTTTACTGAACTCATTGGGACTACTGTGCTAGTTTTG
**ATGTTTT**ATAATGCTGGCATTTAATTACTGGAGAGATTGGATTCTTGGTTGATGATTTAGT**ATTTGTG**AA
TTGTGAAAGTTCAGGAGCTGTGTAGAAAATGTTAGTCAATCAACTTTATTATTGTGCTAAAAGGGGACAT
TCTTATACTGTCCTGTCTAAACTGTTCTCCAGTATAGACTTCCTAGGCACTAAATATCCA**ATATTTA**AAG
GAACACAGCAGGTAAGGAATGAAGCCTCTGAAATAGTACTCATGGATTTATACATGGCAGATCTTACTGT
CTCTACACATTTGGAAGTGTTCGTTGGTTTAAAGAAATGATAGAGGTTTTGAACTACTGACAGTCTTAAA
AGTGAATTTAAAAACTGTTCATACTTTTTATGGTGTAAATTTCCTTTGCTCGATGTCAGTGATTCAGATA
ACTCTTGACCTTGAGATGATGGCTTTTCACAGGTTTCTT**ATATTTT**ATATCTCTTCTGAACATGAATTGT
CATTTTA**GATTTTTGA**CATTTGTATCAAAAGAGAAGTTGAGGAAATCTTCAGAACACTGGTAACTTTTAG
TTTTGCTATAGACTTCAGAAGTGTTTATTT**ATATGTT**CGGTAAATGCTCTCGCATATGCAGTACCTCTTC
TGCCAGCAAATCCAAGGGACCATAGCCTTTTTATGAGACAGGTCACCTCTAGAGGACAACCCAAGAATTA
TTAAAGGAAATGTTACCATTTTGAGAGCATGCTTAAATAAATATTAATAATGTCTTTATAACTTGTTTCC
TTTAAATTTTGGAATATTGAATTACAGGCTTTGGAGGAGTTGTGAAAATTAGGAAAGTTTTTATAT**ATTT**
**TTTGA**AGTGGGCATGGTTGGCTCTTTGAAGACCTATAAAGAGATCCAGTGGGAAGAGTAAGGGTTGGTTC
ATCATCACAAGAAATAAAAAACATAGTG**ATTTTTT**CTCTTAATGTGTAGAGGTGGTTTTACTGGCAATAA
TTAATAATAGATTTCTATTTCAGTATGTAAGCATATTAACTAAAATATGAATTACACTTCCAAAGTTAGA
TTTCTGCTTCAGTAGGTTTGTTTGCTGTGAAGATTACTTCTCAAAAGACAGATGTTCATATTAGCTTAAT
TTTCGGTTTAAATATGTTTGTAAATGATGTAAT**ATATTTC**TTTTGACTAAATGTGGAAAAGTAATGTGTG
TTATACATTGAGAAGTTTTTACTGGCTTTGACTGGAGGTTGTTTTTGCAGAGATGGTATTTTATATGATT
CCAGTATTTGGAAAAGAATTAGTCAAAAGGAATTCACATAGTTTAAATACTGAGAAATTAATATCCAAAT
ATGTACTTGTCTGATTTCTAAATAAGCTGGGGGAGGAGGGAGGGGTGGGAATTGAAATGTGCAAATGAGT
AGTGAATGCTACACTCATTTTCAACTCTTTAACATGAAACTGTTCAATCTTAACACATTGTTACTTTAAT

**ATATGTA**TAAAGAAGTATTACTGTTTGTAAAGCTGCTGTTTGCTTaaaaaaaaaaaaCACCCTTGTCATG
TATTTTCTGTATGTTGGGCCAACAGGTTAGAACATCAACTCatttaaaa**attttta**tctttttttgattt
aaaaaaattcTGTGAAATAATTTATTTACAGACATCTTCCTCCTCCCTCATCCCTTCCAACCTTTACATA
CATCACAGAATCAACCAAACTGTTTGCCTAATCTGAAATCTGAATCCTAATGAGAAAAATTTAAATTTTG
TTGGCACATCACACCTTGAAAGT**ATTTGta**ttattttataatttaatttctaaatataCCACATAAGTTT
ATAATTTAATGTCTTAATTGTAATGCTCTAATAAAAAACTAGCAAAATTAGTGTGAGTTATAACATGAAG
GGATTTTCATCTTTTGCTGTATGAAGGATAATTGTTATATCACATTTGGGGGGTAATAACAGCTTTTTTG
CACTATGTAAATACTAGTGGGGATTCTTCTGTACTAATAAAATGATTATTGAAATGAAA

SECISBP2L-571-3’UTR:
TAATACGACTCACTATAGGTGAATTTAAAAACTGTTCATACTTTTTATGGTGTAAATTTCCTTTGCTCGA
TGTCAGTGATTCAGATAACTCTTGACCTTGAGATGATGGCTTTTCACAGGTTTCTT**ATATTTT**ATATCTC
TTCTGAACATGAATTGTCATTTTA**GATTTTTGA**CATTTGTATCAAAAGAGAAGTTGAGGAAATCTTCAGA
ACACTGGTAACTTTTAGTTTTGCTATAGACTTCAGAAGTGTTTATTT**ATATGTT**CGGTAAATGCTCTCGC
ATATGCAGTACCTCTTCTGCCAGCAAATCCAAGGGACCATAGCCTTTTTATGAGACAGGTCACCTCTAGA
GGACAACCCAAGAATTATTAAAGGAAATGTTACCATTTTGAGAGCATGCTTAAATAAATATTAATAATGT
CTTTATAACTTGTTTCCTTTAAATTTTGGAATATTGAATTACAGGCTTTGGAGGAGTTGTGAAAATTAGG
AAAGTTTTTATAT**ATTTTTTGA**AGTGGGCATGGTTGGCTCTTTGAAGACCTATAAAGAGATCCAGTGGGA
AGAGTAAGGGTTGGTTCATCATCACAAGACACGTG

SECISBP2L-571-3’UTR-ΔSm:
TAATACGACTCACTATAGAGTGAATTTAAAAACTGTTCATACTTTTTATGGTGTAAATTTCCTTTGCTCG
ATGTCAGTGATTCAGATAACTCTTGACCTTGAGATGATGGCTTTTCACAGGTTTCTT**ACACTCT**ATATCT
CTTCTGAACATGAATTGTCATTTTA**GACTCTCGA**CATTTGTATCAAAAGAGAAGTTGAGGAAATCTTCAG
AACACTGGTAACTTTTAGTTTTGCTATAGACTTCAGAAGTGTTTATTT**ACACGCT**CGGTAAATGCTCTCG
CATATGCAGTACCTCTTCTGCCAGCAAATCCAAGGGACCATAGCCTTTTTATGAGACAGGTCACCTCTAG
AGGACAACCCAAGAATTATTAAAGGAAATGTTACCATTTTGAGAGCATGCTTAAATAAATATTAATAATG
TCTTTATAACTTGTTTCCTTTAAATTTTGGAATATTGAATTACAGGCTTTGGAGGAGTTGTGAAAATTAG
GAAAGTTTTTATAT**ACTCTCTGA**AGTGGGCATGGTTGGCTCTTTGAAGACCTATAAAGAGATCCAGTGGG
AAGAGTAAGGGTTGGTTCATCATCACAAGACACGTG

Cul5-209 ENSMUST00000166367.8 cDNA

CDS 7 x NC
3’UTR 17 x NC 1 x C

ACGCCCCCGCCCGCCGCGTCACGTGACGCCGCCACGGACCCTGAGGTGCGGGCCCTAAGCCGAGATAAAG
TCGTTGCCGGCGGCCCAAGCGGGTGCAAGCCCAGCGGCAGAAGCGAAGGCGAGCTGGGGAGGCCCGAGGA
AGTGGCTACTACCTCTTCCCGGTCTGGTCGGCTCCCGGTCCCTTCCCACCATTCGCCGCTCGCGTCTCCT
CAAGCGTTTGCATGCGCTCTCTCGCGTGGGCAGGCCGGGGTGACCATGTAGCTGGAAAGCCCGAGGAAGC
ACGGCTGCCCGGGACGAGCTCGGCGCTGACGGCACGCCGCTCCGGCGTCCCCCGCATCCCCCGCCGCGGC
CTGCGGGGTCCTGCTGGGAACCCCGGCCTCTCGAGGAGGCCTGGCCCCGAGCGCCGCCAAGTCTCGCCCC
GTCTCGCGAGAGTCCAAGTTGAGAAC**ATG**GCGACGTCTA**ATCTGTT**AAAGAATAAAGGTTCTCTCCAGTT
TGAAGACAAGTGGGACTTCATGCATCCAATTGTTTTGAAGCTTTTACGCCAGGA**ATCTGTA**ACAAAACAG
CAGTGGTTTGATCTATTTTCGGATGTACATGCTGTCTGTCTCTGGGATGATAAAGGCTCATCAAAAATTC
ATCAGGCTTTAAAAGAAGATATTCTTGAGTTTATTAAGCAAGCACAGGCTCGTGTACTGAGCCATCAAGA
TGACACAGCTTTGCTGAAGGCATATATTGTTGAATGGCGGAAATTCTTCACACAGTGTG**ATATTTT**ACCA
AAACCTTTTTGTCAATTAGAGGTGACTCTATTGGGTAAACAAAGCAGCAATAAAAAATCAAATATGGAAG
ACAGTATTGTTCGAAAGCTCATGCTTGATACGTGGAATGAGTCGATTTTTTCAAATATAAAGAACAGACT
CCAGGACAGTGCAATGAAGCTGGTGCATGCTGAGAGATTAGGGGAAGCTTTTGATTCCCAGCTGGTCATC
GGGGTGCGAGAGTCCTATGTTAATCTTTGCTCCAACCCCGAGGACAAGCTTCAGATCTATAGGGATAATT
TTGAGAAGGCATACTTGGACTCAACAGAGAGGTTTTATAGAACACAGGCACCCTC**ATATTTA**CAGCAAAA
TGGTGTGCAGAATTACATGAAATATCTCATGGAATGCTGTGTAAATGCGCTGGTGACCTCCTTTAAAGAG
ACTATTTTAGCAGAATGCCAAGGCATGATCAAGCGAAATGAAACTGAAAAGTTACATTTG**ATGTTTT**CCT
TGATGGACAAAGTTCCTAATGGGATAGAGCCGATGTTGAAGGACTTGGAGGAGCATATTATAAGTGCGGG
CCTAGCAGACATGGTGGCCGCAGCTGAAACCATCACTACTGACTCTGAGAAGTATGTGGAGCAATTACTT
ACACTGTTTAATAGATTCAGTAAACTGGTCAAAGAAGCTTTTCAGGATGATCCTCGTTTCCTTACTGCAA
GAGATAAGGCATATAAAGCAGTTGTTAATGATGCTACT**ATATTTA**AACTTGAATTGCCTTTGAAGCAAAA
AGGAGTGGGGTTGAAAACTCAGCCTGAGTCAAAATGCCCCGAGTTGCTTGCCAATTACTGTGACATGTTG
TTAAGGAAAACGCCATTAAGCAAAAAACTAACATCTGAGGAGATTGAAGCAAAGCTTAAAGAAGTGCTCT
TGGTACTTAAATATGTACAAAACAAAG**ATGTTTT**TATGAGGTATCACAAAGCTCATCTTACCCGACGTCT
CATATTGGACATCTCTGCTGATAGTGAGATTGAAGAAAACATGGTAGAGTGGCTAAGAGAAGTTGGTATG
CCAGCAGATTATGTGAACAAGCTTGCTAGAATGTTTCAGGACATAAAAGTATCTGAAGACTTGAACCAAG
CTTTTAAGGAAATGCACAAAAATAATAAGTTGGCATTACCAGCTGATTCCGTTAATATAAAGATTTTGAA
TGCTGGTGCTTGGTCTAGAAGCTCCGAGAAAGTCTTTGTCTCACTTCCTACTGAACTGGAGGATTTGATA
CCTGAAGTAGAAGAATTTTACaaaaaaaaTCACAGTGGTAGAAAATTACACTGGCACCATCTCATGTCAA
ATGGAATTATAACATTTAAAAATGAAGTAGGTCAGTATGATTTGGAAGTAACCACGTTTCAGTTGGCTGT
GTTGTTTGCATGGAACCAAAGGCCTAGAGAGAAAATCAGCTTTGAAAATCTAAAACTTGCAACGGAACTC
CCAGATGCTGAACTTAGAAGGACTTTATGGTCTTTAGTAGCTTTTCCCAAGCTCAAACGGCAAGTTTTGT
TGTATGACCCTCAAGTCAACTCACCCAAAGATTTTACAGAAGGCACCCTCTTCTCAGTGAACCAGGACTT
CAGTCTCATAAAAAATGCAAAAGTACAGAAAAGGGGGAAAATCAATTTGATTGGACGCTTGCAGCTCACT
ACAGAACGAATGAGAGAAGAAGAAAATGAAGGGATAGTCCAACTAAGAATATTAAGAACCCAGGAAGCCA
TCATACAAATAATGAAAATGAGAAAGAAAATTAGCAATGCCCAGCTGCAGACTGAATTAGTAGAAATTCT
GAAAAACATGTTCCTGCCTCAGAAGAAGATGATAAAGGAGCAGATGGAGTGGCTGATTGAACACAGGTAC
ATCCGGAGGGACGAGGCCGACATCAACACCTTCATCTACATGGCC**TAG**CCGGGCCGCTGCTGCCGCACAC
ACGCCCCTGAAGGCCTGGGCAGAGGCTGTCCCAGCCCCAGCTGGAGGAAGCTTTATTTGGACTTTGATTA
CATAAATATTAAACTCTGCCTTACCTTACAAAACGACTCTATTTTGCCAGTCACATTAGTTAGCATGATG
GCATTCCCTTCATGTTGCACACTCTTTAACAGCATGCTGTTTTGTGGAGAAAATTGCATTCATGAAGAGC
CCATTGTGAACTTTCAAAGTCAATTCA**ATTTTTA**CCCACCTAGAGAAATAACATGTCGGAAGGGTGAGGG
TGGGGTTCTTTTTGCTTCTTTTATCCCCTTTTCTTCTTCAAAGAAATATACTTGCACAAGGAAGGATTTT

CAGATATTCATGCACTGAAAAATGCTGGGGAtttttgtttgtttgtttgggttttttaaattctttttttt
ttttttttttttATGAAATTAAAGCTAGACGTAGCACTAAGCTACTTCTGATGCAAAGAATGAATCGTCA
ACACTGTGCTGGGCAAGGGCAGGCACACTTGAGAAGACACGTGTGCCAGTGTCAAGTTGTTTGTCAAACA
CTGCTTTCACTTTAGTGACTGGATCTTAATGGTAACCTGAAATGGTATTAAATATTTCTACCTTATAAAT
CCTGATTTTCAATGAGCAGGAGAGTTCTGTTTAGTTATTTGTTGTTAAATGATAAAGATTTGGG**ATTTTT**
**C**TTTTAAACCTTGTACGGCTGGAGAGCTTGTTTTGAAAACATGAAGTTTATAATGAATGTTGCTTCAGTT
AAAAAATGTGTGGGTATACGTACATTATTGATTGCATAATCAGAATCTTGAAGCAAAAATTAAGTGTGTT
ACTAATACACTCAATTCTCATACGTCTA**ATATTTA**TTATACTGTACCTGAATTTGTTGAAAACAATGCAG
AAATATTCTGATGCAGTGCAGTGAGAGAATTGCTTTCTTAACCTGTAGCATAGAATTATTTGGTGATTGA
AAGTGTTGTGATTAGGGATTTACCACCTCAGTTTTCTGGAAAGACATTCTCTATTTAAAAGTAATAATTA
CCTGCCACTTAGCAGGAAGAAAGGTGAGATTACAACCTGCAACCTGAAAAGCACTCAAAGTTGTTAGTCT
TAACATGAGCAGTAACCTTGCCAA**ATATGTA**CAT**ATATTTA**T**ATATTTT**AAAAATAAAC**ATTTTTA**AAAT
GTTCAGAGGGCAACACCTACCTTGCTCCTCTCAGTGATCCAGAGCAGTGACTCAGGACTTTACAGAGCAG
TGGAAACTTAGAGATTGAGGACTCAAGCAACTGAAGAGAATTATTCCCCAGCAGGAAAAATGGCTCCCTG
TGGATTTTACTCATTGTGGTGCACGTTTCGAGTTTTCTTGCAAATTTTAGAACTTTTGATGTTAAGTAGT
ATTTTGAAGTATTGGTTCTGAAGCTAGCAGAAGATGGAGTGTTGAGAAGCATACAGTATTATCTATCCTA
TTACCACAGAGGGCTGTGGATCTGCCCTGCTGCTCAGCAGCTGGGGTCTCCTTGCTGGAAAGAAGTAAAG
T**ATATGTT**AGATGTTTAAGGCTTTGATAATTATAGTCATATATCTGTTGTGAAACTCATAGGAAGTTGGA
AGTGC**ATGTGTG**CTTGCTGTGCCCTGCAGCTCTGAGGTGATGGCTAAAGCAGGTGTGCAGCGCAAACTCA
GTCCCTGCTGCTCTACCACGTGGAGGATGGAACATCCGGAATTTAGAGGGATAGTTTTATCATGACTGTT
CAATAGCTTGGTGTTGGCAAATTTTCCTGTCCCAATACCAGAATGCCAAAGGAGGAAACAATCAAGGGAA
AAATTTTAGGCCATTTAAAATGACAAAAAAGATTTGTCATTTTAAACCATCATTTTCTTTAGAACATCTT
TGAAATTTCTAATCGCTTTGGTCATTTAAGAATGAGAGCTGTGATGTTTTGATTTATAAAGGCAAACTTA
ATGCAAAGTGGGTAACACTAAATCCATAGCAAAGttttttattaaatttttataa**attttttt**aatagtta
attatgtcctaattgtattttGGGGAATGAGCAGCAGTAAATACCACAGACTTAACAGTTTCATCCTTTT
TTAAAAATATATATATAAGAGTGTAACTACTTTTTGGTGTTTATCAAAACTATTTAGTTGACAAGGTGCC
TATACCATTGTTGGGATTTTCTTTAA**ATGTTTT**ATGAATAGATTTGATAATTTATCTGAAGATCCAGAAC
CACTGACAGCGCCCCTCTCTCTCCCTGCAGACTTCATAGTCTTGCTATCTGAGCAGGGCCAGACATGCAC
AGGGAGGGAAGGAGAGGCCAGCTCCCACTCACACTGGGAGGCATTGTTAAGTGAACGGTACCGCAGGCTC
AGATTTGAGCACTTCTGTCTACCTAAGGTGGCGCC**ATCTGTA**AATGCTAACCTGTCACCAACTCACGGCC
AGAGTAAGGAGACAGGGAACAGATGCATGCAGTGGAAAGGGGAGACTCCTGGACGCGCAGTTAGACAAAT
GTGAGGCCGTCATCAGTCTAATAATTTATAAGGATCCTCAATCAAATTCTGAAGTCTCCGACTTTCACAG
TTCTTGGAAGGGATCCCTGAGTTTACCAGCGTGCTCACTGAAAGCTCTCACTCTTTGTGCAGAAATATTA
AAGTCTCCGAAAAGGTGAACTGTTAATCATTCTT**Attttttt**tAACTCTTCATTAAATTGAATATGATAT
CATTAGCTCTGCTCCAAGGGCAAATTTTCAAGTTTAATCTGGGTGAA**ATATTTG**CTAGTTTACAGAAAGA
TTTGCTATCATATCAATAGCTGGCTCTTCTGTTTTTGTGTGAATGACTGGGATGCTGACACAAGTTGTCC
CAAGGTCACAGTTATGAGAGAAACACTGTTGGAGAGCGTTCCTGTCATCTGCACT**ATGTGTG**GTCTGGAG
TTCCTAAGGTGTAGCCTCTCATCGTGACCTGTACAGTTTTGAATGTGCACCACTACATACCCGGATGGCA
CTGTACAGTTTCCCACGGTAGCAGTCTGTATGCAGTAGGCTGAA**Atatttt**gatgaacgctt**aatttttg**
**g**atttt**attttttt**aagttgtataatttattttCTTGCAAAATAAAAGTGTAATATAAAACATTTCATCTA
TCCAGAAAATCTTGATGTTCTACCATAAAAATTTTTGCAACAGTAAAAAATTTTGGCAagcc

Cul5-3’UTR:
TAATACGACTCACTATAGGTTCAGAGGGCAACACCTACCTTGCTCCTCTCAGTGATCCAGAGCAGTGACT
CAGGACTTTACAGAGCAGTGGAAACTTAGAGATTGAGGACTCAAGCAACTGAAGAGAATTATTCCCCAGC
AGGAAAAATGGCTCCCTGTGGATTTTACTCATTGTGGTGCACGTTTCGAGTTTTCTTGCAAATTTTAGAA
CTTTTGATGTTAAGTAGTATTTTGAAGTATTGGTTCTGAAGCTAGCAGAAGATGGAGTGTTGAGAAGCAT
ACAGTATTATCTATCCTATTACCACAGAGGGCTGTGGATCTGCCCTGCTGCTCAGCAGCTGGGGTCTCCT
TGCTGGAAAGAAGTAAAGT**ATATGTT**AGATGTTTAAGGCTTTGATAATTATAGTCATATATCTGTTGTGA

AACTCATAGGAAGTTGGAAGTGC**ATGTGTG**CTTGCTGTGCCCTGCAGCTCTGAGGTGATGGCTAAAGCAG
GTGTGCAGCGCAAACTCAGTCCCTGCTGCTCTACCACGTGGAGGATGGAACATCCGGAATTTAGAGGGAT
AGTTTTATCATGACTGTTCAATAGCTTGGTGTTGGCAAATTTTCCTGTCCCAATACCAGAATGCCAAAGG
AGGAAACAATCAAGGGAAAAATTTTAGGCCATTTAAAATGACAAAAAAGATTTGTCATTTTAAACCATCA
TTTTCTTTAGAACATCTTTGAAATTTCTAATCGCTTTGGTCATTTAAGAATGAGAGCTGTGATGTTTTGA
TTTATAAAGGCAAACTTAATGCAAAGTGGGTAACACTAAATCCATAGCAAAGtttttattaaattttta
taa**atttttt**aatagttaattatgtcctaattgtattttGGGGAATGAGCAGCAGTAAATACCACAGACT
TAACAGTTTCATCCTTTTTTAAAAATATATATATAAGAGTGTAACTACTTTTTGGTGTTTATCAAAACTA
TTTAGTTGACAAGGTGCCTATACCATTGTTGGGATTTTCTTTAA**ATGTTTT**ATGAATAGATTTGATAATT
TATCTGAAGATCCAGAACCACTGACAGCGCCCCTCTCTCTCCCTGCAGACTTCATAGTCTTGCTATCTGA
GCAGGGCCAGACATGCACAGGGAGGGAAGGAGAGGCCAGCTCCCACTCACACTGGGAGGCATTGTTAAGT
GAACGGTACCGCAGGCTCAGATTTGAGCACTTCTGTCTACCTAAGGTGGCGCC**ATCTGTA**AATGCTAACC
TGTCACCAACTCACGGCCAGAGTAAGGAGACAGGGAACAGATGCATGCAGTGGAAAGGGGAGACTCCTGG
ACGCGCAGTTAGACAAATGTGAGGCCGTCATCAGTCTAATAATTTATAAGGATCCTCAATCAAATTCTGA
AGTCTCCGACTTTCACAGTTCTTGGAAGGGATCCCTGAGTTTACCAGCGTGCTCACTGAAAGCTCTCACT
CTTTGTGCAGAAATATTAAAGTCTCCGAAAAGGTGAACTGTTAATCATTCTT**Atttttt**ttAACTCTTCA
TTAAATTGAATATGATATCATTAGCTCTGCTCCAAGGGCAAATTTTCAAGTTTAATCTGGGTGAA**ATATT**
**TG**CTAGTTTACAGAAAGATTTGCTATCATATCAATAGCTGGCTCTTCTGTTTTTGTGTGAATGACTGGGA
TGCTGACACAAGTTGTCCCAAGGTCACAGTTATGAGAGAAACACTGTTGGAGAGCGTTCCTGTCATCTGC
ACT**ATGTGTG**GTCTGGAGTTCCTAAGGTGTAGCCTCTCATCGTGACCTGTACAGTTTTGAATGTGCACCA
CTACATACCCGGATGGCACTGTACAGTTTCCCACGGTAGCAGTCTGTATGCAGTAGGCTGAA**Atatttt**g
atgaacgctt**aattttttgg**atttt**atttttt**aagttgtataatttattttCTTGCAACACGTG

Cul5-3’UTR-ΔSm:
TAATACGACTCACTATAGGTTCAGAGGGCAACACCTACCTTGCTCCTCTCAGTGATCCAGAGCAGTGACT
CAGGACTTTACAGAGCAGTGGAAACTTAGAGATTGAGGACTCAAGCAACTGAAGAGAATTATTCCCCAGC
AGGAAAAATGGCTCCCTGTGGATTTTACTCATTGTGGTGCACGTTTCGAGTTTTCTTGCAAATTTTAGAA
CTTTTGATGTTAAGTAGTATTTTGAAGTATTGGTTCTGAAGCTAGCAGAAGATGGAGTGTTGAGAAGCAT
ACAGTATTATCTATCCTATTACCACAGAGGGCTGTGGATCTGCCCTGCTGCTCAGCAGCTGGGGTCTCCT
TGCTGGAAAGAAGTAAAGT**ACACGCT**AGATGTTTAAGGCTTTGATAATTATAGTCATATATCTGTTGTGA
AACTCATAGGAAGTTGGAAGTGC**ACGCGCG**CTTGCTGTGCCCTGCAGCTCTGAGGTGATGGCTAAAGCAG
GTGTGCAGCGCAAACTCAGTCCCTGCTGCTCTACCACGTGGAGGATGGAACATCCGGAATTTAGAGGGAT
AGTTTTATCATGACTGTTCAATAGCTTGGTGTTGGCAAATTTTCCTGTCCCAATACCAGAATGCCAAAGG
AGGAAACAATCAAGGGAAAAATTTTAGGCCATTTAAAATGACAAAAAAGATTTGTCATTTTAAACCATCA
TTTTCTTTAGAACATCTTTGAAATTTCTAATCGCTTTGGTCATTTAAGAATGAGAGCTGTGATGTTTTGA
TTTATAAAGGCAAACTTAATGCAAAGTGGGTAACACTAAATCCATAGCAAAGtttttattaaattttta
taa**aCtCtCt**aatagttaattatgtcctaattgtattttGGGGAATGAGCAGCAGTAAATACCACAGACT
TAACAGTTTCATCCTTTTTTAAAAATATATATATAAGAGTGTAACTACTTTTTGGTGTTTATCAAAACTA
TTTAGTTGACAAGGTGCCTATACCATTGTTGGGATTTTCTTTAA**ACGCTCT**ATGAATAGATTTGATAATT
TATCTGAAGATCCAGAACCACTGACAGCGCCCCTCTCTCTCCCTGCAGACTTCATAGTCTTGCTATCTGA
GCAGGGCCAGACATGCACAGGGAGGGAAGGAGAGGCCAGCTCCCACTCACACTGGGAGGCATTGTTAAGT
GAACGGTACCGCAGGCTCAGATTTGAGCACTTCTGTCTACCTAAGGTGGCGCC**ACCCGCA**AATGCTAACC
TGTCACCAACTCACGGCCAGAGTAAGGAGACAGGGAACAGATGCATGCAGTGGAAAGGGGAGACTCCTGG
ACGCGCAGTTAGACAAATGTGAGGCCGTCATCAGTCTAATAATTTATAAGGATCCTCAATCAAATTCTGA
AGTCTCCGACTTTCACAGTTCTTGGAAGGGATCCCTGAGTTTACCAGCGTGCTCACTGAAAGCTCTCACT
CTTTGTGCAGAAATATTAAAGTCTCCGAAAAGGTGAACTGTTAATCATTCTT**ACtCtCt**ttAACTCTTCA
TTAAATTGAATATGATATCATTAGCTCTGCTCCAAGGGCAAATTTTCAAGTTTAATCTGGGTGAA**ACACT**
**CG**CTAGTTTACAGAAAGATTTGCTATCATATCAATAGCTGGCTCTTCTGTTTTTGTGTGAATGACTGGGA
TGCTGACACAAGTTGTCCCAAGGTCACAGTTATGAGAGAAACACTGTTGGAGAGCGTTCCTGTCATCTGC

ACT**ACGCGCG**GTCTGGAGTTCCTAAGGTGTAGCCTCTCATCGTGACCTGTACAGTTTTGAATGTGCACCA
CTACATACCCGGATGGCACTGTACAGTTTCCCACGGTAGCAGTCTGTATGCAGTAGGCTGAA**ACaCtCt**g
atgaacgctt**aaCtCtCgg**atttt**aCtCtCt**aagttgtataatttattttCTTGCAACACGTG

Hif1a-201 ENSMUST00000021530.8 cDNA

CDS 4 x NC
3’UTR 10 x NC 3 x C

GAgcgggcgcgcgcacccctcggcttttccctcccctcgccgcgcgcccgagcgcgcctccgcccttgcc
cgccccctgccgctgcttcagcgcctCAGTGCACAGAGCCTCCTCGGCTGAGGGACGCGAGGACTGTCC
TCGCCGCCGTCGCGGGCAGTGTCTAGCCAGGCCTTGACAAGCTAGCCGGAGGAGCGCCTAGGAACCCGAG
CCGGAGCTCAGCGAGCGCAGCCTGCAGCTCCCGCCTCGCCGTCCCGGGGGGCGTCCCGCCTCCCACCCCG
CCTCTGGACTTGTCTCTTTCTCCGCGCGCGCGGACAGAGCCGGCGTTTAGGCCCGAGCGAGCCCGGGGGC
CGCCGGCCGGGAAGACAACGCGGGCACCGATTCGCC**ATG**GAGGGCGCCGGCGGCGAGAACGAGAAGAAAA
AGATGAGTTCTGAACGTCGAAAAGAAAAGTCTAGAGATGCAGCAAGATCTCGGCGAAGCAAAGAGTCTGA
AGTTTTTTATGAGCTTGCTCATCAGTTGCCACTTCCCCACAATGTGAGCTCACATCTTGATAAAGCTTCT
GTTATGAGGCTCACCATCAGTTATTTACGTGTGAGAAAACTTCTGGATGCCGGTGGTCTAGACAGTGAAG
ATGAGATGAAGGCACAGATGGACTGTTTTTATCTGAAAGCCCTAGATGGCTTTGTGATGGTGCTAACAGA
TGACGGCGACATGGTTTACATTTCTGATAACGTGAACAAATACATGGGGTTAACTCAGTTTGAACTAACT
GGACACAGTGTGTTTGATTTTACTCATCCATGTGACCATGAGGAAATGAGAGAAATGCTTACACACAGAA
ATGGCCCAGTGAGAAAAGGGAAAGAACTAAACACACAGCGGAGCTTTTTTCTCAGAATGAAGTGCACCCT
AACAAGCCGGGGGAGGACGATGAACATCAAGTCAGCAACGTGGAAGGTGCTTCACTGCACGGGCCATATT
CATGTCTATGATACCAACAGTAACCAACCTCAGTGTGGGTACAAGAAACCACCCATGACGTGCTTGGTGC
TGATTTGTGAACCCATTCCTCATCCGTCAAATATTGAAATTCCTTTAGATAGCAAGACATTTCTCAGTCG
ACACAGCCTCGATATGAAATTTTCTTACTGTGATGAAAGAATTACTGAGTTGATGGGTTATGAGCCGGAA
GAACTTTTGGGCCGCTCAATTTATGAATATTATCATGCTTTGGATTCTGATCATCTGACCAAAACTCACC
ATG**ATATGTT**TACTAAAGGACAAGTCACCACAGGACAGTACAGGATGCTTGCCAAAAGAGGTGG**ATATGT**
**C**TGGGTTGAAACTCAAGCAACTGTCATATATAATACGAAGAACTCCCAGCCACAGTGCATTGTGTGTGTG
AATTATGTTGTAAGTGGTATTATTCAGCACGACTTGATTTTCTCCCTTCAACAAACAGA**ATCTGTG**CTCA
AACCAGTTGAATCTTCAGATATGAAGATGACTCAGCTGTTCACCAAAGTTGAATCAGAGGATACAAGCTG
CCTTTTTGATAAGCTTAAGAAGGAGCCTGATGCTCTCACTCTGCTGGCTCCAGCTGCCGGCGACACCATC
ATCTCTCTGGATTTTGGCAGCGATGACACAGAAACTGAAGATCAACAACTTGAAGATGTTCCATTATATA
ATGATGTA**ATGTTTC**CCTCTTCTAATGAAAAATTAAATATAAACCTGGCAATGTCTCCTTTACCTTCATC
GGAAACTCCAAAGCCACTTCGAAGTAGTGCTGATCCTGCACTGAATCAAGAGGTTGCATTAAAATTAGAA
TCAAGTCCAGAGTCACTGGGACTTTCTTTTACCATGCCCCAGATTCAAGATCAGCCAGCAAGTCCTTCTG
ATGGAAGCACTAGACAAAGTTCACCTGAGAGACTTCTTCAGGAAAACGTAAACACTCCTAACTTTTCCCA
GCCTAACAGTCCCAGTGAATATTGCTTTGATGTGGATAGCGATATGGTCAATGTATTCAAGTTGGAACTG
GTGGAAAAACTGTTTGCTGAAGACACAGAGGCAAAGAATCCATTTTCAACTCAGGACACTGATTTAGATT
TGGAGATGCTGGCTCCCTATATCCCAATGGATGATGATTTCCAGTTACGTTCCTTTGATCAGTTGTCACC
ATTAGAGAGCAATTCTCCAAGCCCTCCAAGTATGAGCACAGTTACTGGGTTCCAGCAGACCCAGTTACAG
AAACCTACCATCACTGCCACTGCCACCACAACTGCCACCACTGATGAATCAAAAACAGAGACGAAGGACA
ATAAAGAAGATATTAAAATACTGATTGCATCTCCATCTTCTACCCAAGTACCTCAAGAAACGACCACTGC
TAAGGCATCAGCATACAGTGGCACTCACAGTCGGACAGCCTCACCAGACAGAGCAGGAAAGAGAGTCATA
GAACAGACAGACAAAGCTCATCCAAGGAGCCTTAACCTGTCTGCCACTTTGAATCAAAGAAATACTGTTC
CTGAGGAAGAATTAAACCCAAAGACAATAGCTTCGCAGAATGCTCAGAGGAAGCGAAAAATGGAACATGA
TGGCTCCCTTTTTCAAGCAGCAGGAATTGGAACATTATTGCAGCAACCAGGTGACTGTGCACCTACTATG
TCACTTTCCTGGAAACGAGTGAAAGGATTCATATCTAGTGAACAGAATGGAACGGAGCAAAAGACTATTA
TTTTAATACCCTCCGATTTAGCATGCAGACTGCTGGGGCAGTCAATGGATGAGAGTGGATTACCACAGCT
GACCAGTTACGATTGTGAAGTTAATGCTCCCATACAAGGCAGCAGAAACCTACTGCAGGGTGAAGAATTA
CTCAGAGCTTTGGATCAAGTTAAC**TGA**GCGTTTCCTAATCTCATTCCttttgattgttaatgtttttgtt
cagttgttgttgtttgttgggtttttgtttctgttggtt**atttttgG**ACACTGGTGGCTCAGCAGTCTAT
TT**ATATTTT**CTATATCTAATTTTAGAAGCCTGGCTACAATACTGCACAAACTCAGATAGTTTAGTTTTCA

TCCCCTTTCTACTTAATTTTCATTAATGCTCTTTTTA**ATATGTT**CTTTTAATGCCAGATCACAGCACATT
CACAGCTCCTCAGCATTTCACCATTGCATTGCTGTAGTGTCATTTAAAATGCACCtttttatttattt**at**
**ttttGG**TGAGGGAGTTTGTCCCTTATTGAATT**ATTTTTA**ATGAAATGCCAATATAATTTTTTAAGAAAGC
AGTAAATTCTCATCATGATCATAGGCAGTTGAAAACTTTTTACTC**Atttttt**tCATGTTTTACATGAAAA
TAATGCTTTGTCAGCAGTACATGGTAGCCACAATTGCACAATATATTTTCTTTAAAAAACCAGCAGTTAC
TCATGCAATATATTCTGCATTTATAAAACTAGTTTTTAAGA**AAttttttttGG**CCTATGGAATTGTTAAG
CCTGGATCATGAAGCTGTTGATCTTATAATGATTCTTAAACTGTATGGTTTCTTTATATGGGTAAAGCCA
TTTACATGATATAAAGAAATATGCTTATATCTGGAAGGTATGTGGCATTTATTTGGATAAAATTCTCAAT
TCAGAGAAGTTATCTGGTGTTTCTTGACTTTACCAACTCAAAACAGTCCCTCTGTAGTTGTGGAAGCTTA
TGCTAATATTGTGTAATTGATTATGAAACATAAATGTTCTGCCCACCCTGTTGGTATAAAGACATTTTGA
GCATACTGTAAACAAACAAACAAAAAATCATGCTTTGTTAGTAAAATTGCCTAGTATGTT**GATTTGTTG**A
AAATAT**GATGTTTGG**TTTTATGCACTTTGTCGCTATTAACATCCTTTTTTCATATAGATTTCAATAAGTG
AGTAATTTTAGAAGCATTATTTTAGGAATATAGAGTTGTCATAGTAAACATCTTGTTTTTTCTATGTATA
CTGTATA**AATTTTTCG**TTCCCTTGCTCTTTGTGGTTGGGTCTAACACTAACTGTACTGTTTTGTTATATC
AAATAAACATCTTCTGTGGACCAGGCCCCCTGGGTCAGCGCTTACGTTTAAATAACATTTGTCTCAACAT
TTCTAGCTCATAAAACGATTTCTCAAAAATTTAAGTTCTTTATAAAAATTAGATTGTACATTTCTACATT
CATTTTATTGCCATTTTCTAATGTATG**ATGTGTC**CCTAAATGTCATGTTAAATAATGACATCATAATATT
GCATTGTAAAGAG**AAtttttttttAG**AATTTTGCCATTATAAATGTATGAGTCTATTAAAATATAAAGTA
CAAACCTTCAGTATTTGCAGTATGAATGGAGTAAGTGAAACAGTTCATGAAACATGATCATACTGTTTTG
AGGGCTCAGGCTCCTGCGTGCATGTCTAATCTGTTCCCATTAGCAGGTGAAGGAAGCTAGGGCTGAAACA
AGAGTTTTCCGCGCTCTCAGGGAGCTATGTGGCATGTCAGAATCTTAGGTCTCAGAACATACCTTGTTTG
GTTTTGATATTGGTTTGGTTTGATTCTGGTACATGGCACATTAATATGCAGATACATTATATAGATAATC
ATATATTACCTG**ATGTTTC**TTTACTTTGCCAGCTTTAAAAAAGTATCTTATGCAATTGTGAATTTTAGAA
ACTTCCAAATAAACACCACAAACCTTCCAGCTTA

Hif1a-3’UTR:
TAATACGACTCACTATAGGCGTTTCCTAATCTCATTCCttttgattgttaatgtttttgttcagttgttg
ttgtttgttgggtttttgtttctgttggtt**atttttgG**ACACTGGTGGCTCAGCAGTCTATTT**ATATTTT**
CTATATCTAATTTTAGAAGCCTGGCTACAATACTGCACAAACTCAGATAGTTTAGTTTTCATCCCCTTTC
TACTTAATTTTCATTAATGCTCTTTTTA**ATATGTT**CTTTTAATGCCAGATCACAGCACATTCACAGCTCC
TCAGCATTTCACCATTGCATTGCTGTAGTGTCATTTAAAATGCACCtttttatttattt**atttttGG**TGA
GGGAGTTTGTCCCTTATTGAATT**ATTTTTA**ATGAAATGCCAATATAATTTTTTAAGAAAGCAGTAAATTC
TCATCATGATCATAGGCAGTTGAAAACTTTTTACTC**Atttttt**tCATGTTTTACATGAAAATAATGCTTT
GTCAGCAGTACATGGTAGCCACAATTGCACAATATATTTTCTTTAAAAAACCAGCAGTTACTCATGCAAT
ATATTCTGCATTTATAAAACTAGTTTTTAAGA**AAttttttttGG**CCTATGGAATTGTTAAGCCTGGATCA
TGAAGCTGTTGATCTTATAATGATTCTTAAACTGTATGGTTTCTTTATATGGGTAAAGCCATTTACATGA
TATAAAGAAATATGCTTATATCTGGAAGGTATGTGGCATTTATTTGGATAAAATTCTCAATTCAGAGAAG
TTATCTGGTGTTTCTTGACTTTACCAACTCAAAACAGTCCCTCTGTAGTTGTGGAAGCTTATGCTAATAT
TGTGTAATTGATTATGAAACATAAATGTTCTGCCCACCCTGTTGGTATAAAGACATTTTGAGCATACTGT
AAACAAACAAACAAAAAATCATGCTTTGTTAGTAAAATTGCCTAGTATGTT**GATTTGTTG**AAAATAT**GAT**
**GTTTGG**TTTTATGCACTTTGTCGCTATTAACATCCTTTTTTCATATAGATTTCAATAAGTGAGTAATTTT
AGAAGCATTATTTTAGGAATATAGAGTTGTCATAGTAAACATCTTGTTTTTTCTATGTATACTGTATA**AA**
**TTTTTCG**TTCCCTTGCTCTTTGTGGTTGGGTCTAACACTAACTGTACTGTTTTGTTATATCACACGTG

Hif1a-3’UTR-ΔSm:
TAATACGACTCACTATAGGCGTTTCCTAATCTCATTCCttttgattgttaatgtttttgttcagttgttg
ttgtttgttgggtttttgtttctgttggtt**aCtCtCgG**ACACTGGTGGCTCAGCAGTCTATTT**ACACTCT**
CTATATCTAATTTTAGAAGCCTGGCTACAATACTGCACAAACTCAGATAGTTTAGTTTTCATCCCCTTTC
TACTTAATTTTCATTAATGCTCTTTTTA**ACACGCT**CTTTTAATGCCAGATCACAGCACATTCACAGCTCC

TCAGCATTTCACCATTGCATTGCTGTAGTGTCATTTAAAATGCACCtttttatttattt**aCtCtCGG**TGA
GGGAGTTTGTCCCTTATTGAATT**ACTCTCA**ATGAAATGCCAATATAATTTTTTAAGAAAGCAGTAAATTC
TCATCATGATCATAGGCAGTTGAAAACTTTTTACTC**ACtCtCt**tCATGTTTTACATGAAAATAATGCTTT
GTCAGCAGTACATGGTAGCCACAATTGCACAATATATTTTCTTTAAAAAACCAGCAGTTACTCATGCAAT
ATATTCTGCATTTATAAAACTAGTTTTTAAGA**AACtCtCtCtGG**CCTATGGAATTGTTAAGCCTGGATCA
TGAAGCTGTTGATCTTATAATGATTCTTAAACTGTATGGTTTCTTTATATGGGTAAAGCCATTTACATGA
TATAAAGAAATATGCTTATATCTGGAAGGTATGTGGCATTTATTTGGATAAAATTCTCAATTCAGAGAAG
TTATCTGGTGTTTCTTGACTTTACCAACTCAAAACAGTCCCTCTGTAGTTGTGGAAGCTTATGCTAATAT
TGTGTAATTGATTATGAAACATAAATGTTCTGCCCACCCTGTTGGTATAAAGACATTTTGAGCATACTGT
AAACAAACAAACAAAAAATCATGCTTTGTTAGTAAAATTGCCTAGTATGTT**GACTCGCTG**AAAATAT**GAC**
**GCTCGG**TTTTATGCACTTTGTCGCTATTAACATCCTTTTTTCATATAGATTTCAATAAGTGAGTAATTTT
AGAAGCATTATTTTAGGAATATAGAGTTGTCATAGTAAACATCTTGTTTTTTCTATGTATACTGTATA**AA**
**CTCTCCG**TTCCCTTGCTCTTTGTGGTTGGGTCTAACACTAACTGTACTGTTTTGTTATATCACACGTG

Rem2-202 ENSMUST00000164766.8 cDNA

CDS 0 x NC
3’UTR 1 x NC

TAATACGACTCACTATAGATTAGCATATGTGACCTCATCAGGAGGCGGGACAATTTCCCCAGGTGTCTGG
AGCGGGGAGGGGTGGGGAATATGATgggggggggCAGTATTTAAAGGGAAAAGCTGACAGTGCTGCTGAG
TGAGGAACCGGTGCTCTGAGCCGCTGGGCTGCACTCGCACATGCACGCCG**ATG**CACACGGACCTTGACAC
CGACATGGACATGGACACAGAAACCGTAGCACTTTGTTCTTCCAGCAGCCGCCAGGCCTCCCCACTGGGG
ACACCCACACCAGAAGCAGATACTACACTTCTGAAACAGAAGCCAGAGAAACTGTTAGCAGAGTTGGACC
TGAGCGGGCCTCCTCCTGCTCCTGGGGTCCCCAGACGAAGAGGAAGCATGCCCGTGCCCTACAAACACCA
GCTGCGGCGGGCCCAAGCTGTAGATGAACTTGACTGGCCACCCCAGGCCTCCCCCTCTGGCTCCTCTGAC
TCCTTGGGCTCAGGGGAGGCAGCCCTTACCCAAAAAGATGGCGTCTTTAAGGTCATGCTCGTGGGGGAGA
GTGGCGTGGGCAAGAGCACTCTAGCGGGCACTTTTGGAGGTCTCCAGGGAGACCATGCTCACGAGATGGA
GAACTCAGAGGACACCTATGAGAGACGGATCATGGTGGACAAAGAAGAAGTGACTTTAATTGTTTATGAC
ATCTGGGAACAGGGAGATGCAGGAGGATGGCTGCAGGATCACTGCCTTCAGACGGGGGATGCCTTTCTCA
TCGTCTTCTCAGTGACAGATCGACGAAGCTTCTCTAAAGTTCCAGAAACCCTTCTTCGGCTCCGGGCTGG
GAGGCCCCACCATGACCTACCTGTCATCCTTGTTGGAAATAAGAGTGACCTGGCCCGCTCCCGGGAGGTA
TCACTGGAGGAGGGTCGCCATCTGGCTGGGACGCTGAGCTGCAAGCACATCGAGACGTCGGCCGCTCTCC
ACCACAACACTCGTGAGCTCTTCGAGGGTGCTGTGCGTCAGATCAGGCTGCGGCGGGGCCGGGGTCATGC
CGGGGGCCAGCGACCCGAACCTAGCAGCCCGGACGGCCCCGCGCCGCCTACGCGCCGTGAGAGCCTCACC
AAGAAAGCTAAGCGCTTCCTCGCCAACCTGGTGCCGCGCAACGCTAAGTTCTTCAAGCAACGCTCCAGGT
CATGTCACGACCTCTCTGTGCTC**TGA**GCCACGGTCGCCATGGTCACTGCAGTCGCCATGGTCACCGTGCC
CTCCGCTCGCCCCCTCACCCCACTCCTGTCCGTCTAGGAAACCAAAAATACCCAGGATGCCCTGGTGTGA
GCGGGAGGCGGGGACGGGTAGCTGGTAGGTCCACCACCACCACCTCTCCTGGTCTTAACAGCCGACCATT
CACAGAGCCTCAAGACCTGCAAGTCAGGGAAGAAAACCGTGCTGCAAGGT**ATTTTTT**ATTGTTATTATTA
ACTTGCAAGAAGCCACCTCTCCCGGAAAGACACTCCAAAAACTAGAACCGAAAAAGTGCTTTGTAGCCTC
CTGGATGGAGCTGACCTCCTTGGTACTCGGAACATGCCTTACCTTTAAGTAAGTTTTAAAGGTAAAAACC
CAGAGATGACTTTTCCAGAGATAACTGAAAATTATCCTCTGTCCTGGTCCACTTTGCCCTTAAGAAGTTC
TTCAGAGAAAGGGCTGGAAGTTATTCTTGAAATTGGACTCTGACTTACCATTCTAGTATGGCACGTTTCC
TTTAAGGTTATTGATGTGACGATGTGGGCAGACTCTCTGCATTTAGATGCCCTAAGGTAGATCTGAGGGC
TGGCAGCCCCTTTGCCTAGGGACTAGGAGGCACCAGCAAGGGCCACCCTTTCCCCTTTCCAATAAAtttt
tttttCTATTGCCACGTG

Rem2-ΔSm:
TAATACGACTCACTATAGATTAGCATATGTGACCTCATCAGGAGGCGGGACAATTTCCCCAGGTGTCTGG
AGCGGGGAGGGGTGGGGAATATGATgggggggggCAGTATTTAAAGGGAAAAGCTGACAGTGCTGCTGAG
TGAGGAACCGGTGCTCTGAGCCGCTGGGCTGCACTCGCACATGCACGCCG**ATG**CACACGGACCTTGACAC
CGACATGGACATGGACACAGAAACCGTAGCACTTTGTTCTTCCAGCAGCCGCCAGGCCTCCCCACTGGGG
ACACCCACACCAGAAGCAGATACTACACTTCTGAAACAGAAGCCAGAGAAACTGTTAGCAGAGTTGGACC
TGAGCGGGCCTCCTCCTGCTCCTGGGGTCCCCAGACGAAGAGGAAGCATGCCCGTGCCCTACAAACACCA
GCTGCGGCGGGCCCAAGCTGTAGATGAACTTGACTGGCCACCCCAGGCCTCCCCCTCTGGCTCCTCTGAC
TCCTTGGGCTCAGGGGAGGCAGCCCTTACCCAAAAAGATGGCGTCTTTAAGGTCATGCTCGTGGGGGAGA
GTGGCGTGGGCAAGAGCACTCTAGCGGGCACTTTTGGAGGTCTCCAGGGAGACCATGCTCACGAGATGGA
GAACTCAGAGGACACCTATGAGAGACGGATCATGGTGGACAAAGAAGAAGTGACTTTAATTGTTTATGAC
ATCTGGGAACAGGGAGATGCAGGAGGATGGCTGCAGGATCACTGCCTTCAGACGGGGGATGCCTTTCTCA
TCGTCTTCTCAGTGACAGATCGACGAAGCTTCTCTAAAGTTCCAGAAACCCTTCTTCGGCTCCGGGCTGG
GAGGCCCCACCATGACCTACCTGTCATCCTTGTTGGAAATAAGAGTGACCTGGCCCGCTCCCGGGAGGTA
TCACTGGAGGAGGGTCGCCATCTGGCTGGGACGCTGAGCTGCAAGCACATCGAGACGTCGGCCGCTCTCC

ACCACAACACTCGTGAGCTCTTCGAGGGTGCTGTGCGTCAGATCAGGCTGCGGCGGGGCCGGGGTCATGC
CGGGGGCCAGCGACCCGAACCTAGCAGCCCGGACGGCCCCGCGCCGCCTACGCGCCGTGAGAGCCTCACC
AAGAAAGCTAAGCGCTTCCTCGCCAACCTGGTGCCGCGCAACGCTAAGTTCTTCAAGCAACGCTCCAGGT
CATGTCACGACCTCTCTGTGCTC**TGA**GCCACGGTCGCCATGGTCACTGCAGTCGCCATGGTCACCGTGCC
CTCCGCTCGCCCCCTCACCCCACTCCTGTCCGTCTAGGAAACCAAAAATACCCAGGATGCCCTGGTGTGA
GCGGGAGGCGGGGACGGGTAGCTGGTAGGTCCACCACCACCACCTCTCCTGGTCTTAACAGCCGACCATT
CACAGAGCCTCAAGACCTGCAAGTCAGGGAAGAAAACCGTGCTGCAAGGT**ACCCCCT**ATTGTTATTATTA
ACTTGCAAGAAGCCACCTCTCCCGGAAAGACACTCCAAAAACTAGAACCGAAAAAGTGCTTTGTAGCCTC
CTGGATGGAGCTGACCTCCTTGGTACTCGGAACATGCCTTACCTTTAAGTAAGTTTTAAAGGTAAAAACC
CAGAGATGACTTTTCCAGAGATAACTGAAAATTATCCTCTGTCCTGGTCCACTTTGCCCTTAAGAAGTTC
TTCAGAGAAAGGGCTGGAAGTTATTCTTGAAATTGGACTCTGACTTACCATTCTAGTATGGCACGTTTCC
TTTAAGGTTATTGATGTGACGATGTGGGCAGACTCTCTGCATTTAGATGCCCTAAGGTAGATCTGAGGGC
TGGCAGCCCCTTTGCCTAGGGACTAGGAGGCACCAGCAAGGGCCACCCTTTCCCCTTTCCAATAAAtttt
tttttCTATTGCCACGTG
